# Fitness translocation: improving variant effect prediction with biologically-grounded data augmentation

**DOI:** 10.1101/2024.12.17.628831

**Authors:** Adrien Mialland, Shuzo Fukunaga, Riku Katsuki, Yunfei Dong, Hideki Yamaguchi, Yutaka Saito

**Author notes:** National Institute of Advanced Industrial Science and Technology (Adrien Mialland, Yutaka Saito).

## Abstract

Data scarcity limits the characterization of protein fitness landscapes and the development of accurate variant effect prediction models. To address this challenge, we introduce *fitness translocation*, a data augmentation strategy that generates synthetic variants for a target protein by leveraging variant fitness data previously measured in homologous proteins. Using embeddings from protein language models, the method computes the difference between each homolog variant and its wild type and applies these offsets to the target wild-type embedding to create synthetic variants in embedding space. We illustrate the utility of fitness translocation in the context of variant effect prediction on three protein families: IGPS, GFP, and SARS-CoV-2 spike proteins, across different models and training data sizes. Fitness translocation consistently improves predictive performance, particularly under limited training data, and is effective even when augmenting with remote homologs sharing as little as 35% sequence identity. These results illustrate how biologically grounded data augmentation can expand and diversify protein fitness landscapes, supporting more data-efficient protein engineering. The code is available at https://github.com/adrienmialland/ProtFitTrans.

## 1 Introduction

Understanding the relationship between protein sequence and function is central to protein engineering. Mapping this relationship accurately enables the rational design of proteins with novel or enhanced functions and supports the interpretation of mutations in both natural and engineered contexts [1]. In this setting, fitness refers to a quantitative measure of how well a protein performs a desired function such as enzymatic activity, binding affinity, or fluorescence. The fitness landscape describes the conceptual mapping from sequences to their corresponding fitness values, illustrating how sequence variations influence functional performance [2]. However, characterizing this landscape requires sufficient experimental observations of sequence–function relationships.

Exploring the fitness landscape experimentally is inherently limited by the vastness of sequence space. Even mutating a modest number of amino acid sites generates a combinatorial explosion of possible variants, 20^*k*^ for *k* sites, making comprehensive measurements infeasible. While modern technologies such as deep mutational scanning and high-throughput screening allow the creation of large protein variant libraries [3–5], measuring fitness for every sequence remains costly and often less scalable than library construction [4]. As a result, most experimentally characterized fitness landscapes are sparsely sampled. Machine learning models have been widely applied to learn the mapping from sequence to fitness using available experimental data [1, 6–9]. This task, known as variant effect prediction, enables in silico exploration of unmeasured sequences but is strongly constrained by the size and diversity of the training data. Limited coverage often prevents models from generalizing beyond the regions already observed, highlighting the need for strategies that can effectively expand and diversify the available training data.

Data augmentation involves generating synthetic training data from existing measurements to increase dataset diversity and improve model generalization without requiring additional experiments. In fields such as computer vision and natural language processing, approaches like image transformations or sentence paraphrasing are widely used to enhance model performance [10, 11]. In protein variant effect prediction, however, direct analogs of these techniques are less straightforward, and effective augmentation strategies remain underexplored. A biologically motivated alternative is to leverage information from homologous proteins, which often share structural and functional features due to common ancestry. Recent studies suggest that fitness landscapes may be partially conserved across homologs [12], indicating that variant fitness measurements from one homolog could be leveraged to inform predictions in another.

Here, we introduce a data augmentation strategy called *fitness translocation*, which generates synthetic variants for a target protein by transferring variant fitness information from homologous proteins. Given variant fitness data from a homolog, embeddings derived from protein language models (pLMs) [13, 14] are used to compute the difference between each variant and its wild type, capturing the mutational change as a vector offset. These offsets are then applied to the embedding of the target wild type to produce synthetic variants in embedding space, effectively augmenting the available target training data. We illustrate the utility of fitness translocation in the context of variant effect prediction, evaluating it on three protein families with diverse biological functions and experimental fitness assays: IGPS homologs [12], GFP homologs [15], and SARS-CoV-2 spike proteins [16], covering enzymatic activity, fluorescence intensity, and cell entry or receptor binding. Using various prediction models and a range of training data sizes, we show that fitness translocation consistently improves prediction performance, particularly in low-data regimes, and we present algorithms to identify which homolog datasets are most beneficial for augmentation. These results highlight the potential of biologically grounded data augmentation to enable more data-efficient exploration of protein fitness landscapes.

## 2 Methods

### 2.1 Problem Setting and Overview

Accurately characterizing protein fitness landscapes is challenging because only a small fraction of possible variants can typically be measured due to practical constraints. This sparse sampling limits our ability to predict the fitness of unobserved variants, a task known as variant effect prediction, in which models are trained on available experimental data to generalize across sequence space. To improve performance in such low-data scenarios, we propose to augment available training data using fitness-labeled variants from homologous proteins with related sequences and functions. While homologs differ from the target protein in sequence, we assume that they often share similar fitness landscape due to their common ancestry, providing an opportunity to transfer variant fitness information from homologs to the target.

Our proposed method, named fitness translocation, uses pLMs to represent sequences in an embedding space (Figure 1). In this space, we analyze how mutations shift the embedding of a homolog’s wild-type sequence. These shifts, referred to as mutation offsets, are then applied to the target wild type to generate synthetic variants in the embedding space. These synthetic variants are integrated with real variants to expand training data for prediction models.

**Figure 1.**
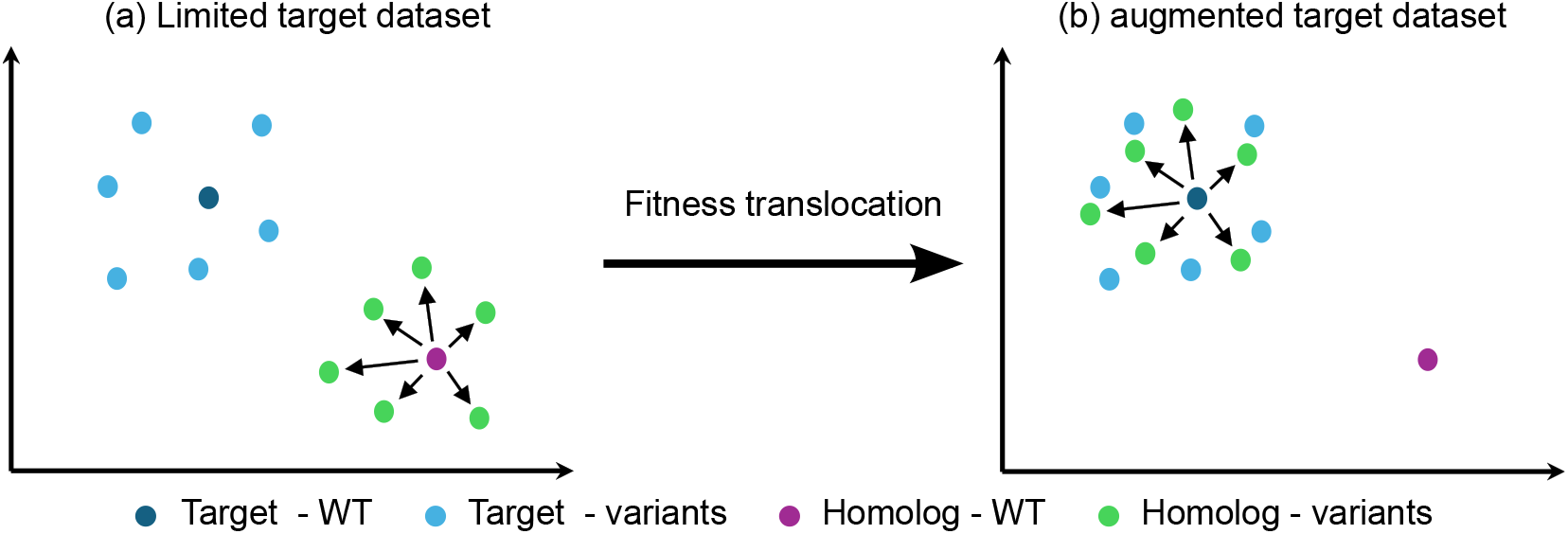
Concept of Fitness Translocation. WT: Wild-Type. (a) The embeddings of the homolog’s variants shift the wild-type embedding. (b) When fitness translocation is applied, these shifts, or mutation offsets, are applied to the target wild-type to generate synthetic variants.

### 2.2 Protein Language Model Embeddings

To represent protein sequences in a form suitable for machine learning and data augmentation, we use pLMs, deep learning models pretrained on large corpora of natural protein sequences. pLMs are used to calculate a sequence embedding, a fixed-length numerical vector representing sequence information. Because pLMs are pretrained on diverse protein families, they enable meaningful comparisons between sequences even across homologs, providing the foundation for fitness translocation. We used ESM-2 [14] or ESM-1v [17] pLMs (esm2_t33_650M_UR50D and esm1v_t33_650M_UR90S_1 respectively), but other existing pLMs can be used (e.g. [18, 19]).

### 2.3 Fitness Translocation

Fitness translocation is a data augmentation method that generates synthetic training data for a target protein by transferring variant fitness data obtained in a homologous protein. Given a homolog, we first compute embeddings for both its wild-type and variants using pLM. For each variant, we calculate a mutation offset, defined as the difference between the variant embedding and the wild-type embedding (Figure 1). This offset captures how the mutation shifts the representation of the homolog in the embedding space. To generate a synthetic variant for the target protein, we add the mutation offset from each homolog variant to the target wild-type embedding. The resulting synthetic embedding simulates how an analogous mutation might affect the target protein, under the assumption that the fitness landscape is at least partially conserved between the homolog and the target. We associate the synthetic variant with the same fitness value as the original homologous variant, normalized by the corresponding wild-type fitness. While this may not reflect the true fitness of the unmeasured target variant, it provides a training signal grounded in evolutionarily related data.

The set of synthetic variants, each represented as an embedding paired with a fitness label, is combined with (added to) the original target training data. By augmenting the target dataset in this way, fitness translocation increases the diversity of training data without requiring new experimental measurements. In addition, it operates only in the embedding space and requires no sequence alignment between the homolog and the target protein. Synthetic variants are generated from all homolog variants, independent of residue identity or position, which simplifies applicability across homologs with varying sequence similarity. This approach is expected to be beneficial in low-data regimes where variant effect prediction models struggle to generalize from limited training data.

### 2.4 Prediction Models and Training Setup

To evaluate the effect of data augmentation by fitness translocation, we trained supervised regression models to predict the variant fitness from a sequence embedding. Each training sample consists of a sequence embedding, either from a real variant or a synthetic variant generated via fitness translocation, and the corresponding fitness label. When training with augmented data, synthetic variants were added to the training data alongside the measured variants from the target protein. We tested three types of regression models: support vector regression (SVR), random forest regression (RF), and linear regression with L1 regularization (Lasso). Hyperparameters were selected through nested cross-validation (CV) [20] on training data.

For each target protein, we trained models under multiple training sizes to simulate low to moderate data scenarios. We used held-out test data from the same protein to evaluate prediction with and without augmentation. In all settings, test data contained only real variants that were never seen during training, ensuring a fair comparison between augmentation and baseline conditions.

### 2.5 Dataset

We evaluated fitness translocation using three protein families with experimentally measured variant fitness data: IGPS homologs, GFP homologs, and SARS-CoV-2 spike proteins. Each family includes multiple related proteins, allowing us to examine augmentation across homologs with varying levels of sequence identity (Table 1) and similarity (Table S1.1), and biological context. Detailed characteristics are provided in Tables S1.2 and S1.3 (fitness statistics, assay type, sequence lengths, and mutated sites) and Figures S1.1–S1.3 display fitness distributions for all homologs.

**Table 1.** Summary of the dataset. The pairwise sequence identities and similarities are computed using EMBOSS Needle and EMBOSS Water [21], respectively for global and local alignment.

| Protein Family | Homolog | Number of Variants | EMBOSS Needle |  |  | EMBOSS Water |  |  |
| --- | --- | --- | --- | --- | --- | --- | --- | --- |
|  |  |  | SsIGPS | TmIGPS | TtIGPS | SsIGPS | TmIGPS | TtIGPS |
| IGPS | SsIGPS | 1497 | 100% | 35.5% | 39.6% | 100% | 37.1% | 42.9% |
|  | TmIGPS | 1502 | 35.5% | 100% | 35.0% | 37.1% | 100% | 36.4% |
|  | TtIGPS | 1489 | 39.5% | 35.0% | 100% | 42.9% | 36.4% | 100% |
|  |  |  | amacGFP | cgreGFP | ppluGFP | amacGFP | cgreGFP | ppluGFP |
| GFP | amacGFP | 35500 | 100% | 42.7% | 17.9% | 100% | 45.1% | 25.2% |
|  | cgreGFP | 26165 | 42.7% | 100% | 18.5% | 45.1% | 100% | 23.2% |
|  | ppluGFP | 32260 | 17.9% | 18.5% | 100% | 25.2% | 23.2% | 100% |
|  |  |  | XBB.1.5 | BA.2 | – | XBB.1.5 | BA.2 | – |
| SARS-CoV-2 Spike | XBB.1.5 | 7029 | 100% | 98.9% | – | 100% | 98.9% | – |
|  | BA.2 | 7129 | 98.9% | 100% | – | 98.9% | 100% | – |

IGPS homologs [12]: This set includes three homologs of imidazole glycerol phosphate synthase (IGPS) originated from *Thermogama maritima* (TmIPGS), *Sulfolobus Solfataricus* (SsIGPS), *Streptococus thermopjilus* (TtIGPS). Variant fitness data are measured for single-point mutants of each IGPS homolog based on growth rate under histidine-limited conditions, which represents the enzymatic activity of IGPS.

GFP homologs [15]: This family includes green fluorescent protein (GFP) originated from *Aequorea macrodactyla, Hydrozoa* (amacGFP), *Clytia gregaria, Hydrozoa* (cgreGFP), *Pontellina plumata, Copepoda* (ppluGFP). The fitness is defined by fluorescence intensity, measured for variants of each GFP homolog.

SARS-CoV-2 spike proteins [16]: This dataset includes spike protein variants from the two SARS-CoV-2 omicron strains: *XBB*.*1*.*5* and *BA*.*2* Variant fitness is quantified by ACE2 binding affinity or cell entry efficiency.

For each family, we designated one protein as the target for variant effect prediction and used one or more homologs as augmentation sources for fitness translocation. Variant fitness values were normalized by the wild-type fitness of the respective protein.

### 2.6 Homolog Selection Algorithms

When multiple homologs are available for a given protein family, not all are necessarily beneficial for data augmentation. We developed the homolog selection algorithm to identify which homologs are most informative for fitness translocation to a given target. It evaluates, across multiple train/validation splits, which homologs consistently improve model performance when added to the target dataset. Given a target protein, a set of homologs, a number *N* of data split, and a significance threshold *α*, the first selection stage measures the average change in predictive accuracy (denoted Δ_*µ*_) when each individual homolog is translocated into the target data (Figure 1). We then compute whether this change is statistically significant across the data splits, retaining only the homologs that yield a significant positive effect. In the second stage, these selected homologs are further evaluated in combination, translocating one homolog after another from the highest to the lowest scoring. Each translocated homolog is kept only if it improves the performance of the previous combination, in order to identify the optimal set for fitness translocation. The formal description of our algorithm is shown in Algorithm S2 (Supplementary Data).

## 3 Results

We evaluated our data augmentation strategy, fitness translocation, on 60 configurations: (4 SARS-CoV-2 + 3 IGPS + 3 GFP) targets ×2 pLMs × 3 predictors. For a given configuration (associated with a target protein), 26 target training sizes from 45 up to 1125 variants were considered to undergo fitness translocation. For a given target training size, the homolog selection algorithm was applied to identify the optimal set of homologs, followed by the translocation of all possible sets of homologs for comparisons. The performances of the variant effect prediction tasks were evaluated using Spearman’s correlation between true and predicted fitness values over train/validation splits, and a number of *N* = 50 splits were considered to extract the corresponding sample of Spearman’s correlation. Two samples of Spearman’s correlation were then statistically compared using a one-sided paired t-test to evaluate whether the mean difference indicated an increase in performances, and a significant threshold of *α* = 0.05 was used. Therefore, the corresponding minimal value that a paired t-test can reliably detect depends on the distribution of the samples, and is given by the following formula [22]:

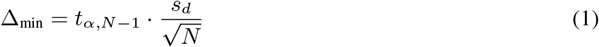

Where *α* is the significance threshold, *N* is the number of train/validation splits, *t*_*α*,*N* −1_ is the critical value of the Student’s t-distribution for *N* − 1 degrees of freedom at the significance level *α*, and *s*_*d*_ is the standard deviation of the paired differences. Consequently, the minimum detectable improvement Δ_min_ decreases as *N* increases and depends on the ability of a predictor to provide stable performances over multiple splits.

### 3.1 Embedding-space effects of fitness translocation

Figure 2 shows the effect of fitness translocation using Principal Component Analysis (PCA) on IGPS embeddings. Homolog embeddings initially separated are evenly aggregated after translocation. Given a target protein, the synthetic variants from remaining homologs are evenly positioned near the target. SARS-CoV-2 spike embeddings show similar behaviour (Figure S3.2), while GFP embeddings show less coherent aggregation, partially overlapping but mostly separated (Figure S3.3).

**Figure 2.**
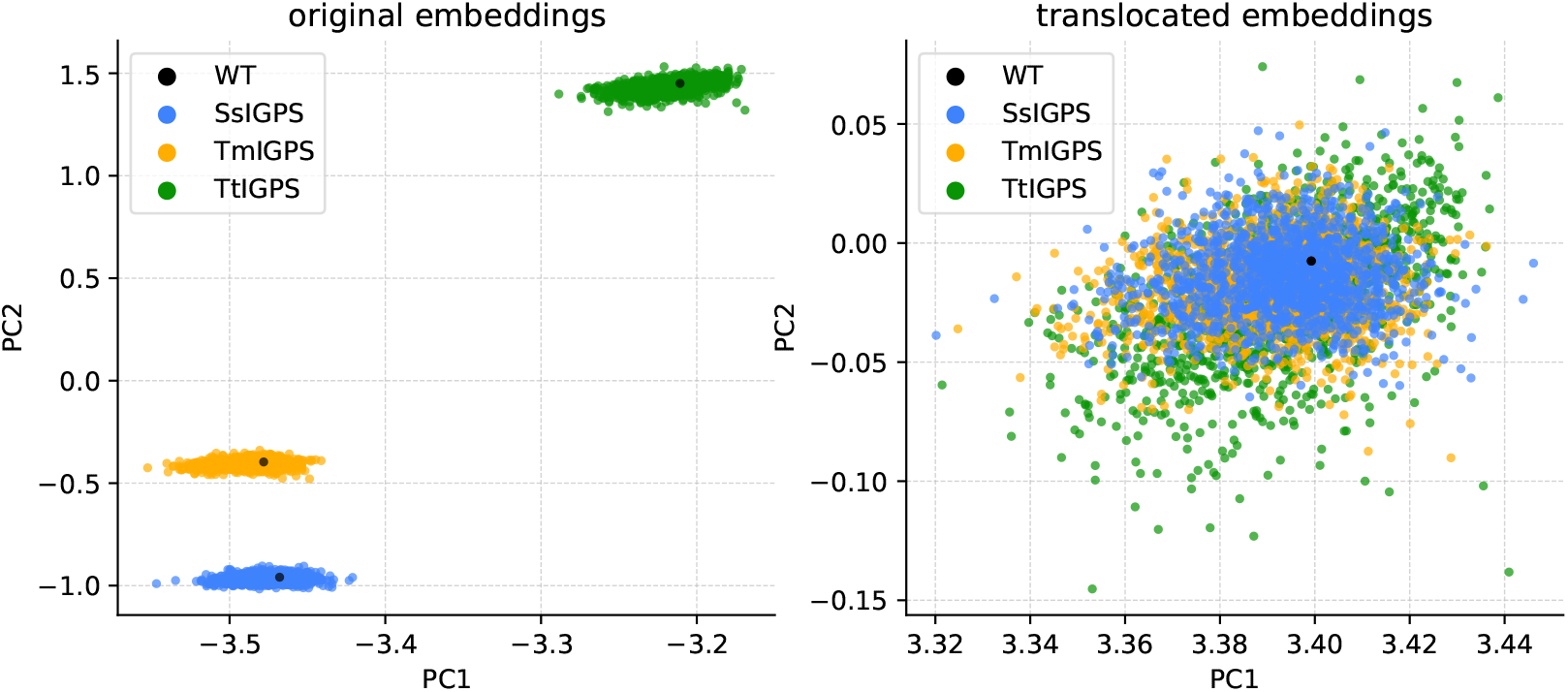
Effect of fitness translocation on IGPS embeddings shown with PCA. Variant embeddings from different homologs are initially separated in embedding space but evenly aggregated after fitness translocation, reflecting the transfer of mutational effects to the target protein.

### 3.2 Fitness Translocation Evaluation

Figure 3, 4, and 5 respectively show representative results of IGPS, GFP, and SARS-CoV-2 spike protein cell entry (ACE2 binding available in the Supplementals Figure S19.1, as it shows similar behaviour to cell entry) for all possible sets of translocated homologs, using the ESM2 pLM and the SVR predictor. The optimal set of homologs selected through the homolog selection algorithm, are then highlighted with a red line over all target training sizes. The additional results are reported in the Supplementary Data.

**Figure 3.**
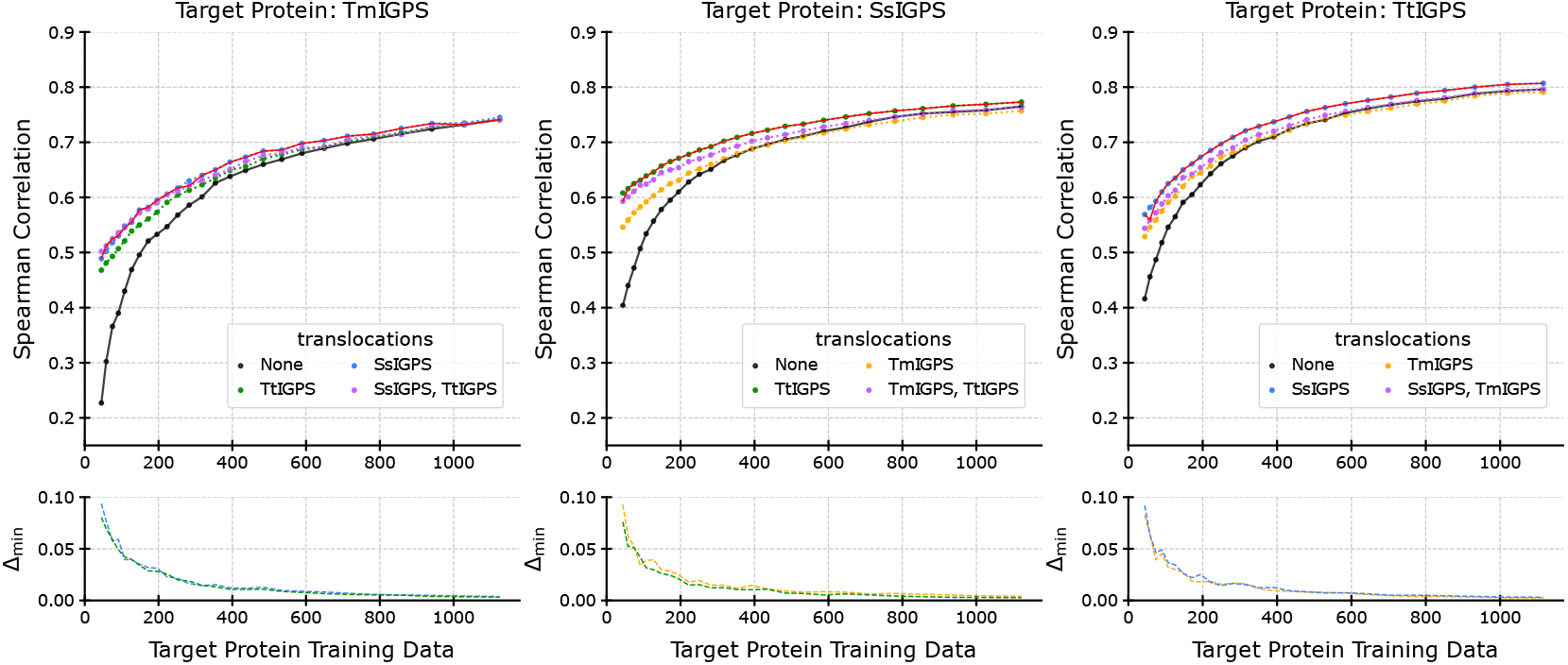
**Top:** Results for IGPS, ESM2 pLM, and SVR predictor. All combination of homologs yielded significant improvement, especially for smaller target training sizes. The optimal set of homolog selected by our method is highlighted with the red line, which shows its effectiveness for maximizing the improvement. **Bottom:** minimum average paired difference in performances Δ_min_ that can be detected (Equation 1). It is inversely proportional to the training size and depends also on *N* .

**Figure 4.**
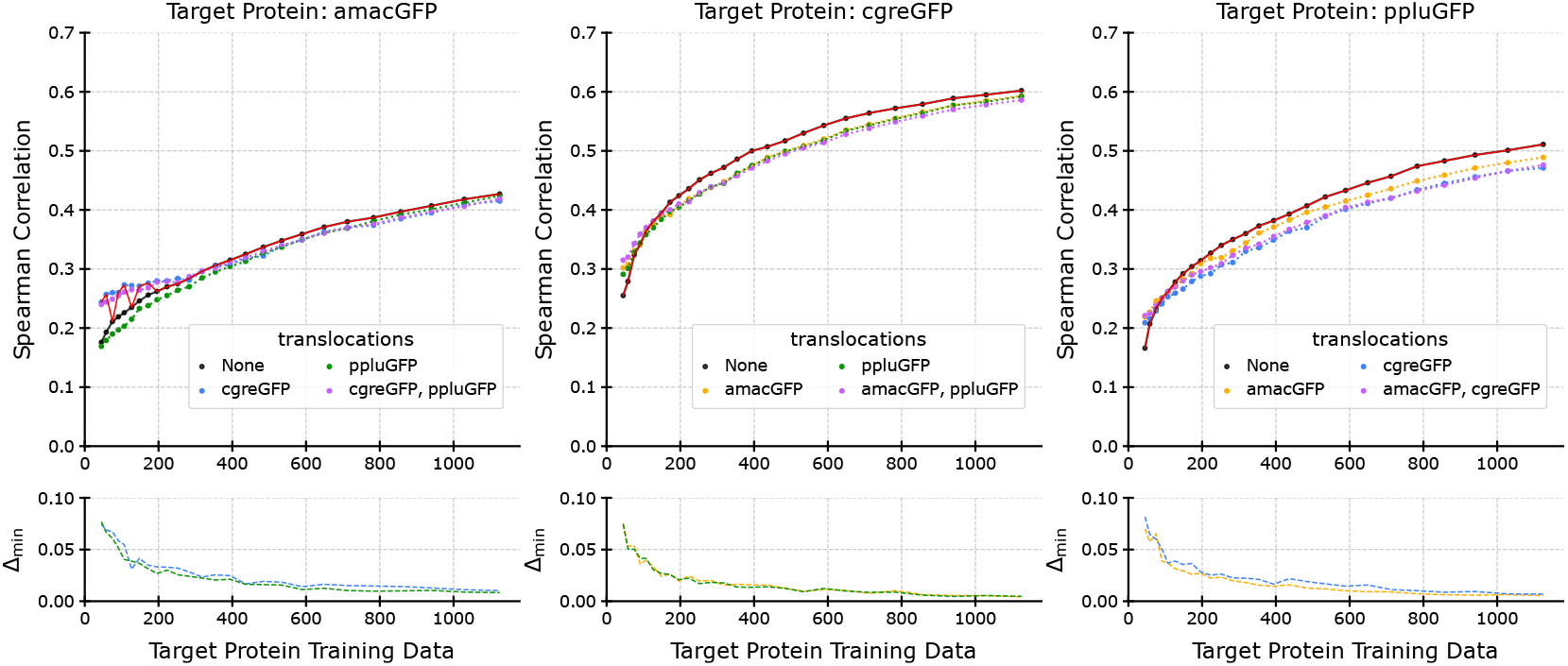
**Top:** Results for GFP, ESM2 pLM, and SVR predictor. Limited improvement could be achieved, and only for small target training size, but few other configurations still yielded substantial increase in performances (e.g Figure S17.2). The optimal set of homolog selected by the homolog selection algorithm is highlighted with the red line, and when no improvement was achieved, the original dataset was consistently selected as the best one. **Bottom:** minimum average paired difference in performances Δ_min_ that can be detected (Equation 1). It is inversely proportional to the training size and depends also on *N*. The red line fluctuations arise from statistical instability, where increasing *N* would reduce the uncertainty and allow for smaller Δ_min_ detection.

**Figure 5.**
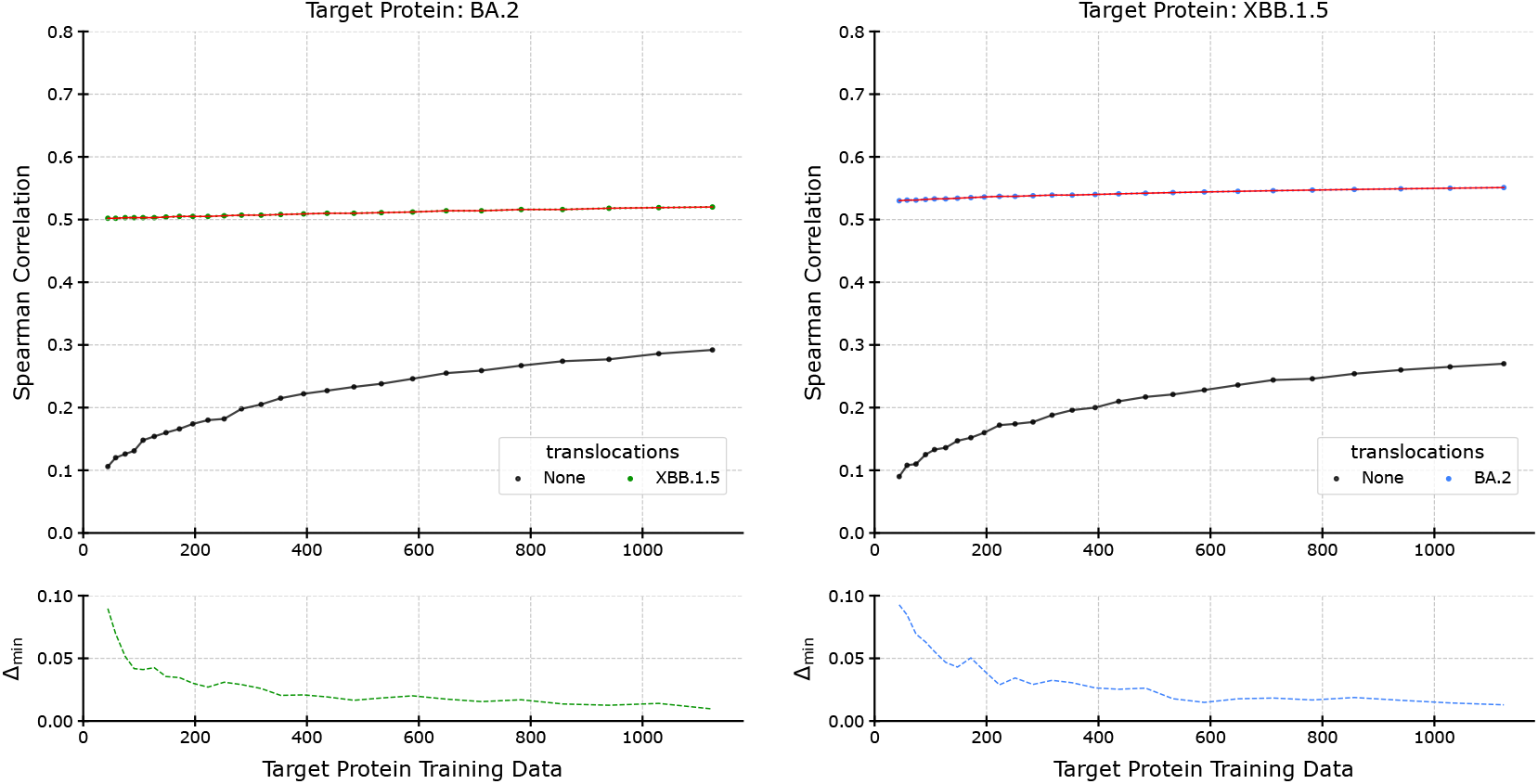
**Top:** Results for SARS-CoV-2 spike protein cell entry, ESM2 pLM, and SVR predictor. It achieved the highest increase in performances among all protein family, which allowed the homolog selection algorithm to easily select the translocated homolog, as highlighted with the red line. **Bottom:** minimum average paired difference in performances Δ_min_ that can be detected (Equation 1). It is inversely proportional to the training size and depends also on *N* .

These results reveal that incorporating additional protein data in a target protein dataset can substantially improve the performance of a variant effect prediction task. The effect was particularly visible for smaller sizes and was the biggest for SARS-CoV-2 cell entry (Figure S6, S10), followed by SARS-CoV-2 ACE2 binding (Figure S7, S11), IGPS (Figure S8, S12) and GFP (Figure S9, S13) families. Consistent improvement could be achieved with IGPS and SARS-CoV-2 spike proteins, with the lowest sequence identity of 35% observed between TmIGPS and TtIGPS (Table 1). Improvement on GFP family was limited and occurred for smaller target training sizes in most cases, although some configurations yielded substantial improvements (Figure S21.2). Besides, the homolog selection algorithm was able to reliably select among the best performing set of homologs, that achieves a significant increase in performances on the target protein variants effect prediction task. Especially, when the relative improvement between different homologs combinations was small, the lowest scoring combinations were systematically excluded (e.g Figure 3, S16.1). Also, in the case where no improvement could be achieved, the original target dataset was consistently selected as the best one (e.g Figure 4, S17.3).

This behaviour stems from the use to the one-sided paired t-test. It naturally establishes an adaptive upper threshold and minimizes the influences of the statistical fluctuations from non-effective combinations, which could be selected accidentally. Therefore, the initial selection of homologs is improved. In cases where the improvement was too small to reach statistical significance, increasing *N* may benefit the selection, to decrease the minimum detectable improvement Δ_min_. In particular, the variations observed for smaller target training sizes in GFP family (Figure S17, S21), arose from the limited improvements it provided, which could not consistently reach statistical significance, while showcasing reduced but noticeable improvements. In other words, statistical fluctuations could make the average (over *N*) improvement appear inconsistent, preventing detection when the true average is close to Δ_min_, whereas a larger *N* would smooth out these fluctuations and decrease Δ_min_. In addition, for all configurations that we evaluated (Figure S14, S15, S16, S17, S18, S19, S20, S21), the minimum detectable improvement Δ_min_ followed the expected trend expressed by Equation 1. For the smallest target training size, the sample size *N* = 50 resulted in Δ_min_ between 0.1 and 0.07, which then rapidly decreased as the target training size increased, to reach between 0.017 and 0.005 for the biggest target training sizes. In other words, the fitness translocation method naturally allows to detect smaller improvements as the target training size increases.

### 3.3 Evaluation Under Grouped Splitting

Our fitness translocation method uses a standard random splitting strategy for cross-validation, assuming no hypothesis about the fitness distribution. We provide additional evaluations under grouped splitting on IGPS proteins. Because IGPS mutation sites evenly span the eight *β*-strands, variants were grouped by *β*-strand: for each nested cross-validation iteration, four strands were randomly assigned to the target test set, while the homolog training set included only the remaining four strands. SVR predictors were trained with each homolog successively used as the target and using the previously best-performing homolog for augmentation with similar hyperparameters. Results (Table S5.1) show that fitness translocation still provides substantial improvements, with similar trends to random splitting but with reduced magnitude.

### 3.4 Ablation Study on Homolog Selection Algorithm

We conducted an ablation study to assess the contribution of individual components in the homolog selection procedure. In the first ablation scenario, we removed the statistical testing in the first selection stage and used the cross-validation performance of the target training data as a selection threshold. The second stage was kept unchanged to sort out the initial selection. So, in situations where each translocated homolog individually yielded substantial improvements in the first stage, this ablation scenario and the original selection method led to similar final sets, as the second stage operated on similar sets (e.g. Figure S28, S40). However, when an individual homolog achieved poor performances in the first stage, the lack of paired t-test did not allow to effectively manage the statistical fluctuations (e.g. Figure S32.2, S43.2, S30.2), often resulting in suboptimal selections with poorer generalization capability.

The second ablation scenario starts with the same configuration as the first scenario – removing the statistical testing in the first selection stage. However, we also removed the second selection stage that sorts out the initial selected set, which resulted in all homologs from the first stage being selected as the final set. this approach led to similar considerations as the first ablation scenario, with the additional downside that the lack of sorting could not allow for the initial selection of homologs to be refined (e.g Figure S29.3, S40.3, S43.3), resulting in the worst results.

### 3.5 Performance Trends Across Homolog Combinations

The assessment of all possible combination showed that the results were dependent on the type of predictor and the combination being translocated, with consistency across various configurations.

#### Absolute Amplitudes Trend

In all family − target − pLM − predictor configurations, the effect of the translocation eventually reached a saturation point as the target training sizes increased (Figure S8, S9, S10, S11), or indicated a trend toward saturation (Figure S6, S7, S10, S11). In other words, the performance difference between with and without fitness translocation becomes small as the target training size increases. The SARS-COV-2 spike proteins for cell entry had 12 configurations tested (2 targets × 2 pLMs × 3 predictors, Figure S6, S10) and had the biggest improvement, followed by SARS-CoV-2 spike proteins for ACE2 bindings that also had 12 configurations tested (Figure S7, S11). These results were consistent across all configurations and were substantial for all 26 target training sizes. The IGPS family had 18 configurations tested (3 targets × 2 pLMs × 3 predictors, Figure S8, S12) and had the second biggest improvement. It was consistent across target − pLM configurations, and reaches saturation toward the higher target training sizes, but was dependent on the predictor. Specifically, the RF predictor resulted in decreased performances for the higher target training sizes (Figure S8.3, S12.3). The GFP family had 18 configurations tested (Figure S10, S13) and had the lowest performances. It was consistent across pLMs models, but showed varying trend over the target training sizes, and was dependent on both the target and the predictor. However, some configurations still provided substantial improvement for all 26 target training sizes (Figure S9.2, S13.2). Besides, as an overall trend, the final performances were mainly dependent on the content of the translocated combination and the type of predictor, where specific combinations could either increase or decrease the original performances. The effects of the pLMs were mostly marginal and mostly acted on the overall magnitude of the results, rather than the inherent capabilities of the additional datasets. However, the GFP family did not exhibit a clear pattern, showing varying performances depending of the choice of species used as the target (Figure S9, S13).

#### Relative Amplitudes Trend

We assessed the relative amplitude between pairs of translocated combinations for IGPS and GFP families. For a given target, the relative amplitude between each pair showed a repeatable trend over each pLMs and each predictor. In other word, the translocated combinations could reliably be classified from the highest to the lowest performing. This was especially visible when the relative amplitude was more pronounced, and a representative example can be seen with IGPS family and SsIGPS species used as the target (Figure 3). The best improvement over both ESM model and all 3 predictors was consistently observed with the translocation of TtIGPS, followed by TmIGPS + TtIGPS together, and then by TmIGPS alone (Figure S8, S12). Even for the RF predictor, that performed poorer than the SVR and Lasso predictor, no significant disruption of the overall trend could be noticed. Also, it should be specified that when the relative amplitude between pairs of combinations is not substantial (e.g. Figure S8.3, S12.3), the overall trend can probably be assessed as a whole as any impact on the trend, over different configurations, may be considered as statistical variations. Besides, and more interestingly, the relative amplitudes tended to be conserved over different targets. For instance, in IGPS family the translocation of TmIGPS consistently performed the worst over all targets that used it, and over all ESM and predictor configurations (Figure S8, S12). Finally, despite the reduced improvement that fitness translocation provided on GFP family, it still exhibited similar trends in relative amplitude between the translocated combinations (Figure S9, S13).

## 4 Discussion

This study demonstrates the potential of fitness translocation, a biologically-grounded data augmentation method that addresses the challenge of experimental data scarcity in protein engineering. We show how variant effect prediction task can be improved by leveraging fitness landscapes of related proteins, enabling the construction of augmented datasets that improve the prediction performance on a target protein. These findings highlight how leveraging homologous fitness data can improve the predictive performance by making efficient use of prior experimental efforts.

### Comparison with Traditional Methods

data augmentation methods have been commonly used in various modern machine learning approaches, demonstrating their effectiveness in fields such as computer vision [23, 11] or natural language processing (NLP) [24, 25]. These methods range from basic manipulations, such as rotation or cropping, to the generation of synthetic data using advanced methods. However, these approaches are difficult to apply on protein data, which are subject to complex sequence-function relationships and where even a single residue mutation can significantly alter the properties of a given protein [26]. Recent investigations explored the advantages of image and text related data augmentation methods for protein sequence prediction [27]. The authors suggest that while improvement can be achieved, the performance gains may be suboptimal and that innovative methods tailored to protein data are required. Therefore, our proposed fitness translocation method provides a simple framework for protein data augmentation, enabling the use of protein data and fitness landscapes from past experiments, without relying on synthetic data and with no alteration of the sequence-function pairs.

Fitness translocation also differs conceptually from commonly used mutation scoring approaches such as sequence conservation and MSA-based models, which rely on evolutionary statistics from aligned homologous sequences, or pLM log-odds (zero-shot) prediction, which compares the likelihood of a variant sequence to the wild-type using a protein language model without experimental data. In contrast, fitness translocation leverages fitness values previously measured in homologous proteins within a supervised learning framework. Operating directly in embedding space, it does not require explicit sequence alignment between the homolog and the target protein. Fitness translocation is therefore complementary to zero-shot approaches, enabling the reuse of experimentally characterized variants from related proteins to improve supervised prediction on the target protein.

### Fitness Translocation Evaluation

The generalizability of the fitness translocation method across multiple configurations, as shown by our results, makes it a reliable and easy to apply data augmentation approach for enhancing target protein datasets. Even for the GFP protein family, where improvements were not consistent, significant gains were observed under specific conditions (Figure S9.2, S13.2). This suggests that certain protein species may only benefit from the translocation of a subsets of homologs, highlighting the potential advantage of increasing the number of homolog’s datasets considered for fitness translocation. However, this may result in a significant increase in processing time if all possible combinations are evaluated. With a number of *m* homologs, each one of them being either included or not, this would require a number of 2*m* combinations to be evaluated. However, the homolog selection algorithm in our method allows to effectively prioritize the most promising combinations and reducing the number of evaluations to a maximum of 2*m*. This upper limit is only maximized if all homologs are selected during the first selection stage. The use of a one-sided paired t-test can further improve both the processing times and the final performances, by reducing the impact of statistical variations and only considering the meaningful improvements. In that regard, Equation 1 provides a simple solution to set a relevant minimum detectable improvement Δ_min_. Although increasing *N* does not scale linearly with Δ_min_, which is proportional to 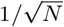, the use of *N* = 50 in this study was sufficient to achieve a Δ_min_ below 0.1 for the smallest target training size, with Δ_min_ rapidly decreasing as the training size increased (Figures 3, 4, 5). In addition, Δ_min_ also depends on *s*_*d*_, which is fixed by a given configuration. So, for a specific configuration, using the one-sided paired t-test with our homolog selection algorithm and a reasonably high number *N* can both provide reliable and meaningful results in a reduced processing time. Besides, embeddings computation did not represent a prohibitive cost (Table S3.1). The longest sequences (SARS-CoV-2 Spike) required approximately 0.123 seconds per sequence on an NVIDIA A6000 GPU and 2.861 seconds on a CPU, while the largest dataset (GFP) required about 35 minutes on GPU and 13 hours on CPU.

### Biological Validity of Fitness Translocation

Recent investigations into phylogenetically divergent IGPS proteins with the TIM barrel fold, showing only 35-39.6% sequence identity (Table 1), have explored the evolutionary constraints imposed by sequence and structural requirements [12]. The study revealed unexpected long-range allosteric pathways linking distal residues to the active site, which likely play a critical role in maintaining function. Also, a significant correlation was observed between the fitness landscapes of the distant orthologs. The authors suggested that amino-acid preferences at a given position in the structure are mostly conserved during evolution. These findings propose a general mechanism by which evolutionary pressures contribute to the conservation of fitness landscapes, where such pressures maintain critical biophysical and functional features of the protein, despite sequence divergence. Additionally, previous studies on closely related homologs (influenza virus nucleoproteins), with 94% sequence identity, reported similar results on amino-acid preferences at specific positions [28], further supporting the notion that fitness landscapes are shaped by conserved structural and functional properties, even for low sequence identity. So, the results from the study on IGPS led the authors to interpret landscapes conservation as the translocation of fitness landscapes in the sequence space.

Building on these findings, our study employed pre-trained pLMs to generate latent representations of protein sequences, leveraging their ability to capture intricate structural and functional features encoded in amino acids [13]. These models are pre-trained on large-scale protein sequence datasets containing billions of amino acids and learn internal representations by predicting masked amino acids. This allows the model to infer context-dependent features in a self-supervised manner, producing embeddings that represent the biological properties of protein sequences, while preserving the sequence-function relationship [29]. Our findings show that the translocation of the fitness landscape effectively take advantage of the conserved properties between protein species, with the possibility to generalize beyond the particular TIM barrel fold.

### Implications for Protein Engineering

By introducing sequence diversity in datasets for machine learning models, fitness translocation has the potential to improve model generalization capabilities across diverse contexts, facilitating applications such as enzyme engineering or therapeutic development. In the case of directed evolution, the properties of a protein sequence are iteratively improved by mimicking the process of natural selection. It involves several rounds of mutagenesis, expression, and selection, where each round builds upon the previous one to construct a library of sequences that contains high quality variants. Its effectiveness is therefore dependent on the selection process, where a machine learning model should predict the best set of sequences to include in subsequent round [8]. The cost and time required for directed evolution make the fitness translocation method particularly suited to increase the model capabilities, select high quality variants, and limit the number of rounds. More generally, many applications that rely on sequence-function mappings of protein variants would likely benefit from fitness translocation. As an example, modern methods for designing artificial proteins with specific functions, or *de novo* protein design, often involves generative models trained on embeddings from a pLM [30]. These methods may therefore benefit from an augmented dataset with fitness translocation.

## 5 Conclusion

In this study, we introduced *fitness translocation*, a data augmentation technique for protein variant effect prediction which leverages evolutionary conservation of fitness landscapes. This method enables the translocation of sequence space embedding features from homologs protein datasets to a target protein, enriching its training data with biologically relevant variations from previous protein engineering efforts. We also presented a systematic selection framework that identifies the most beneficial set of homologs for data augmentation. This approach significantly improved the target protein variant effect prediction and demonstrates the utility of fitness translocation, particularly in scenarios with limited data. The implementation of the method is available at https://github.com/adrienmialland/ProtFitTrans.

## Supporting information

supplemental: Algorithm, Tables, Figures

datasets: sequences and fitnesses

## 6 Data Availability

The data underlying this article are available in the article and in its online supplementary material. The code is available at https://github.com/adrienmialland/ProtFitTrans

## 7 Supporting Information

Algorithm S2 and Figure and Tables S1.1-S51.3 (PDF)

## 9 Author contributions statement

AM and YS developed the method and wrote the paper. AM wrote the code and performed the computational experiments. SF, RK, HY, and YD participated in the preliminary data analysis. All authors read and approved the final version of the manuscript.

## 10 Acknowledgments

This work was supported by JSPS KAKENHI (22H03691), AMED grant (JP19ak0101122), NEDO grant (JPNP14004), JACI Prize for Encouraging Young Researcher. The computations were partially performed on the NIG supercomputer at ROIS National Institute of Genetics, the ABCI supercomputer at AIST, and the SQUID supercomputer at Cybermedia Center, Osaka University through the HPCI System Research Project (hp230057, hp240075).

