## supplemental: Algorithm, Tables, Figures for "Fitness translocation: improving variant effect prediction with biologically-grounded data augmentation"

#### Supplemental S1: Protein Identity, Similarity, Fitness, and mutations.

Table S1.1: Summary of the dataset. The pairwise sequence similarities are computed using EMBOSS Needle and EMBOSS Water [16], respectively for global and local alignment.

| Protein Family | Homolog | Number of Variants | EMBOSS Needle | | | EMBOSS Water | | |
|---|---|---|---|---|---|---|---|---|
| | | | SsIGPS | TmIGPS | TtIGPS | SsIGPS | TmIGPS | TtIGPS |
| IGPS | SsIGPS | 1497 | 100% | 56.7% | 55.8% | 100% | 58.8% | 60.1% |
| | TmIGPS | 1502 | 56.7% | 100% | 49.1% | 58.8% | 100% | 51.2% |
| | TtIGPS | 1489 | 55.8% | 49.1% | 100% | 60.1% | 51.2% | 100% |
| | | | amacGFP | cgreGFP | ppluGFP | amacGFP | cgreGFP | ppluGFP |
| GFP | amacGFP | 35500 | 100% | 62.6% | 29.9% | 100% | 65.9% | 41.5% |
| | cgreGFP | 26165 | 62.6% | 100% | 32.3% | 65.9% | 100% | 38.4% |
| | ppluGFP | 32260 | 29.9% | 32.3% | 100% | 41.5% | 38.4% | 100% |
| | | | XBB.1.5 | BA.2 | – | XBB.1.5 | BA.2 | – |
| SARS-CoV-2 Spike | XBB.1.5 | 7029 | 100% | 98.8% | – | 100% | 98.9% | – |
| | BA.2 | 7129 | 98.9% | 100% | – | 98.9% | 100% | – |

Table S1.2: Summary of the dataset. Fitness range and mean $pm$ SD for each homolog alongside their corresponding assay.

| Protein | Homolog | Assay | Fitness Range | Mean ± SD |
|---|---|---|---|---|
| IGPS | SsIGPS | Enzymatic Activity | -3.392 – 1.239 | -0.549 ± 0.611 |
| | TmIGPS | Enzymatic Activity | -2.157 – 1.951 | -0.687 ± 0.608 |
| | TtIGPS | Enzymatic Activity | -2.223 – 0.862 | -0.461 ± 0.608 |
| GFP | amacGFP | Fluorescence | -1.318 – 0.301 | -0.320 ± 0.454 |
| | cgreGFP | Fluorescence | -1.726 – 0.106 | -0.850 ± 0.770 |
| | ppluGFP | Fluorescence | -1.455 – 0.219 | -0.355 ± 0.517 |
| Sars-CoV-2 spike | XBB.1.5 | Cell Entry | -6.862 – 0.340 | -1.406 ± 2.066 |
| | BA.2 | Cell Entry | -6.619 – 0.175 | -0.895 ± 1.792 |
| | XBB.1.5 | ACE2 Binding | -3.726 – 4.013 | 0.036 ± 0.525 |
| | BA.2 | ACE2 Binding | -7.446 – 5.283 | 0.238 ± 0.867 |

Table S1.3: Summary of the dataset. Sequence and mutation informations for each homolog.

| Protein | Homolog | Num. of Variants | Seq. lenght | Num. of Mut. Sites | Coverage Type | Num. of Mutations |
|---|---|---|---|---|---|---|
| IGPS | SsIGPS | 1497 | 222 | 80 | Saturation | 20 AA/site |
| | TmIGPS | 1502 | 221 | 80 | Saturation | 20 AA/site |
| | TtIGPS | 1489 | 220 | 80 | Saturation | 20 AA/site |
| GFP | amacGFP | 35500 | 244 | 237 | Error-prone PCR | avg 3/variant |
| | cgreGFP | 26165 | 235 | 235 | Error-prone PCR | avg 3/variant |
| | ppluGFP | 32260 | 222 | 221 | Error-prone PCR | avg 3/variant |
| SARS-CoV-2 spike | XBB.1.5 | 7029 | 1253 | 1234 | Targeted | avg 2/variant |
| | BA.2 | 7129 | 1253 | 1240 | Targeted | avg 2/variant |
| | XBB.1.5 | 7029 | 1253 | 1149 | Targeted | avg 2/variant |
| | BA.2 | 7129 | 1253 | 1196 | Targeted | avg 2/variant |

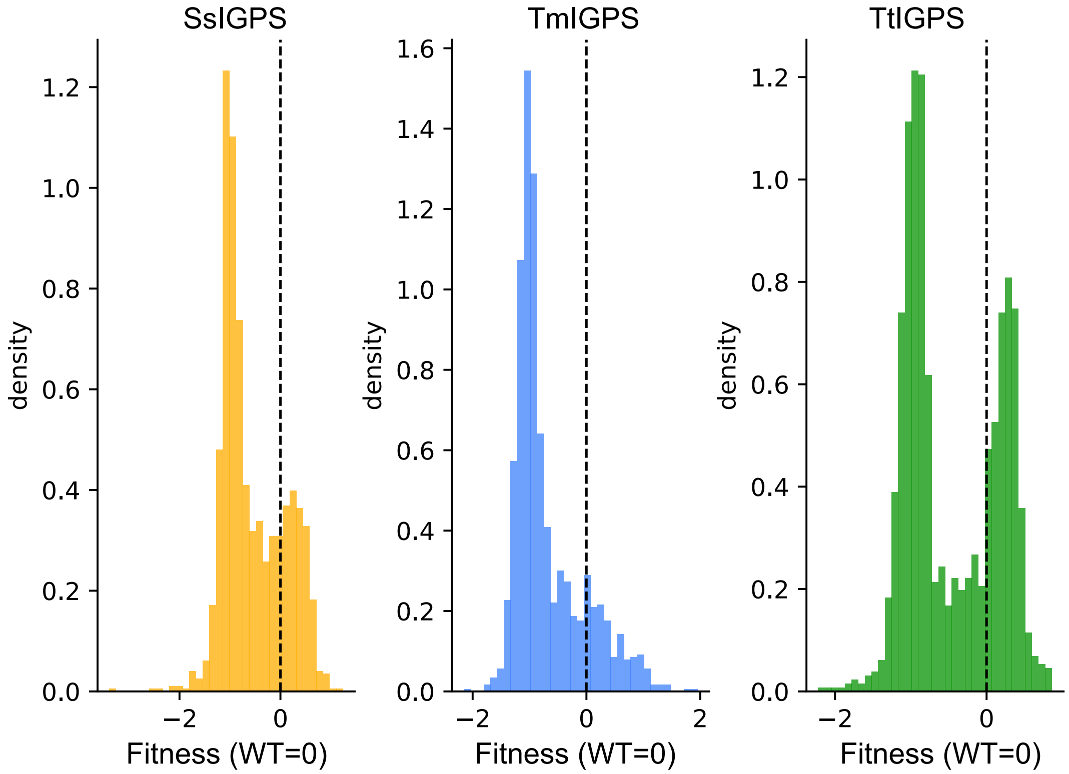


Figure S1.1: IGPS orthologs fitness distributions. The dotted vertical line is the WT=0 fitness

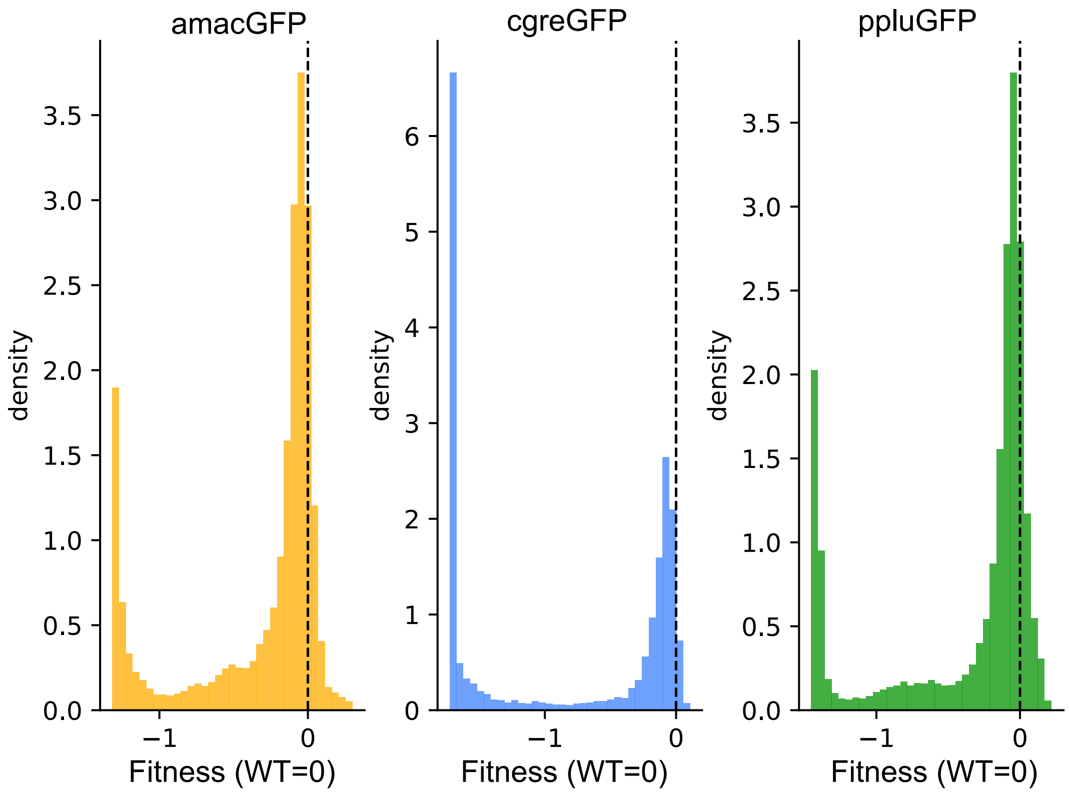


Figure S1.2: GFP orthologs fitness distributions. The dotted vertical line is the WT=0 fitness

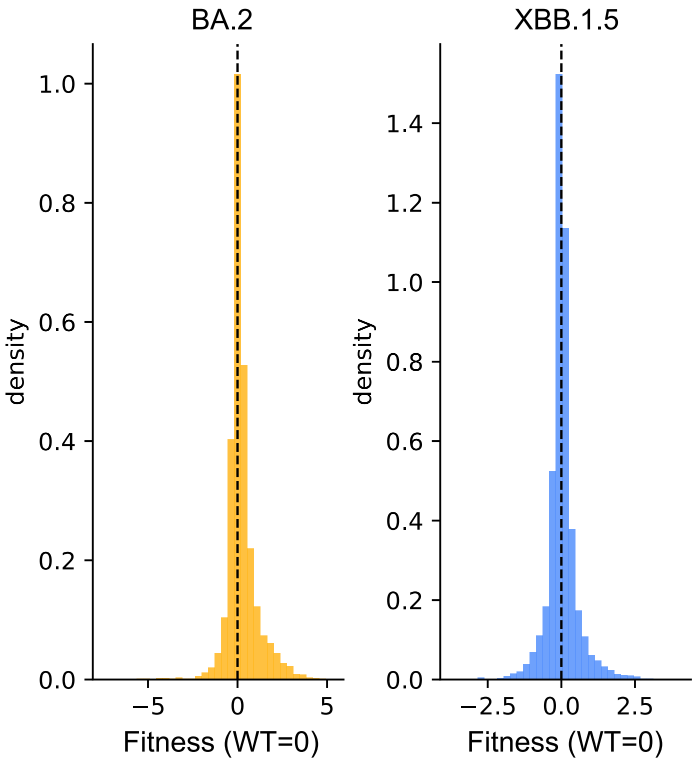


Figure S1.3: SARS-CoV-2 orthologs (ACE2 binding assay) fitness distributions. The dotted vertical line is the WT=0 fitness

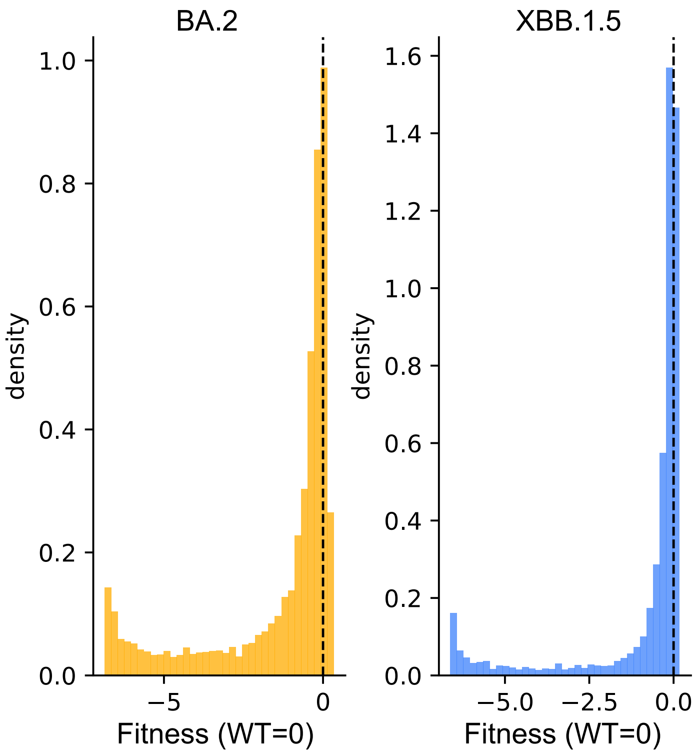


Figure S1.4: SARS-CoV-2 orthologs Spike (cell entry assay) fitness distributions. The dotted vertical line is the WT=0 fitness

#### Supplemental S2: homolog selection algorithms

**Algorithm 1** homolog selection algorithm

**Require:** target, homologs, *N*, $\alpha$
  `differences` $\leftarrow$ []
  `significant` $\leftarrow$ []

  **for** $n \leftarrow 1$ to *N* **do** ▷ Compute paired differences
    *train, validation* $\leftarrow$ *split_dataset*(*target*)
    *t_result* $\leftarrow$ *evaluate_model*(*train*, *validation*)
    **for all** *hom* $\in$ *homologs* **do**
      *trans* $\leftarrow$ *translocate*(*target*, *hom*)
      *result* $\leftarrow$ *evaluate_model*(*train*||*trans*, *validation*)
      append (*hom*, *result* $-$ *t_results*) to `differences`
    **end for**
  **end for**

  **for all** *hom* $\in$ *homologs* **do** ▷ Compute differences significance
    $\sigma_e \leftarrow$ *standard_error*(`differences`, *hom*)
    $\Delta\mu \leftarrow$ *mean_difference*(`differences`, *hom*)
    **if** *p-value*$(\Delta\mu, \sigma_e, N) < \alpha$ **then**
      append (*hom*, $\Delta\mu$) to `significant`
    **end if**
  **end for**

  $hom_{\text{best}}, \Delta\mu_{\text{best}} \leftarrow$ *extract_best*(`significant`)
  `significant` $\leftarrow$ *remove_best*(`significant`)

  *candidate_homs* $\leftarrow$ *sort_by_*$\Delta\mu$(`significant`)
  `optimal_set` $\leftarrow hom_{\text{best}}$

  **for all** *hom* $\in$ *candidate_homs* **do** ▷ Sort out candidate combinations
    *new_results* $\leftarrow$ []
    **for** $n \leftarrow 1$ to *N* **do**
      *trans* $\leftarrow$ *translocate*(*target*, *hom*, `optimal_set`)
      *result* $\leftarrow$ *evaluate_model*(*train*||*trans*, *validation*)
      append *result* to *new_results*
    **end for**

    $\Delta\mu_{\text{new}} \leftarrow mean($*new_results*$)$

    **if** $\Delta\mu_{\text{new}} > \Delta\mu_{\text{best}}$ **then**
      $\Delta\mu_{\text{best}} \leftarrow \Delta\mu_{\text{new}}$
      add *hom* to `optimal_set`
    **end if**
  **end for**

  **return** `optimal_set`

#### Supplemental S3: Original and Translocated Embedding Comparison

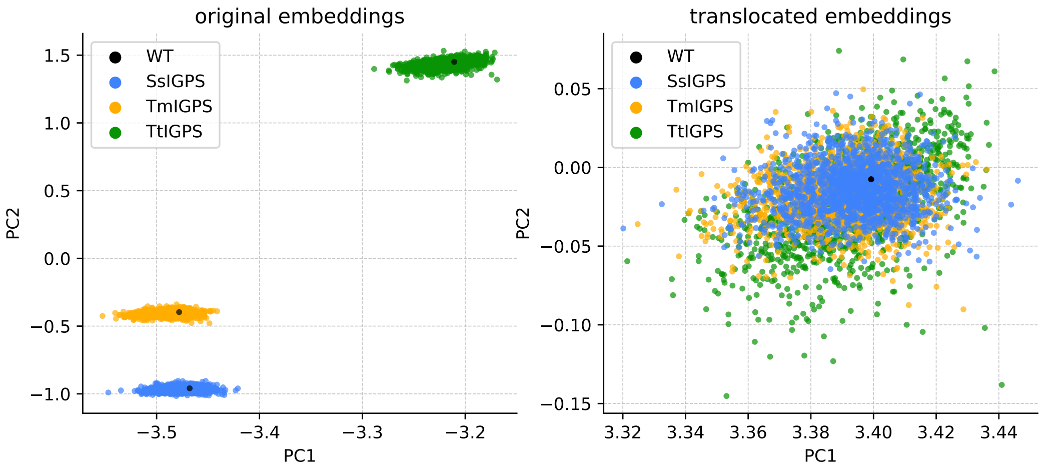


Figure S3.1: Principal Component Analysis (PCA) of IGPS protein embeddings, before and after fitness translocation. Variant embeddings from different homologs are initially separated in embedding space but evenly aggregated after fitness translocation, reflecting the transfer of mutational effects to the target protein.

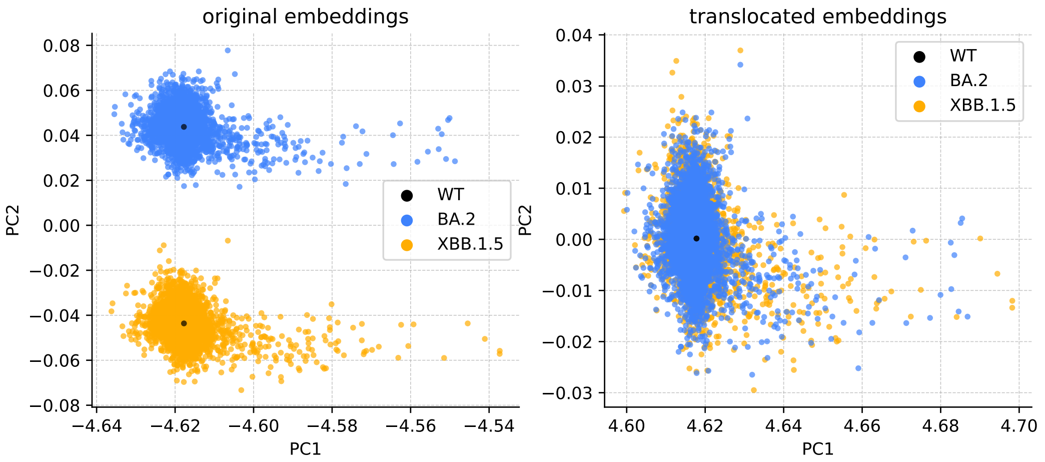


Figure S3.2: Principal Component Analysis (PCA) of SARS-CoV-2 Spike protein embeddings, before and after fitness translocation. Variant embeddings from different homologs are initially separated in embedding space but evenly aggregated after fitness translocation, reflecting the transfer of mutational effects to the target protein.

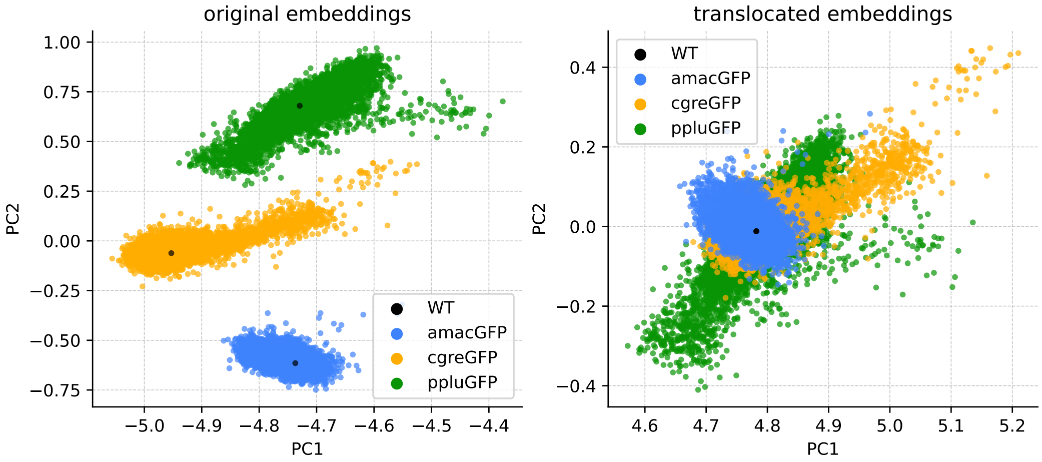


Figure S3.3: Principal Component Analysis (PCA) of GFP protein embeddings, before and after fitness translocation. Variant embeddings from different homologs are initially separated in embedding space but partially aggregated after fitness translocation, reflecting the transfer of mutational effects to the target protein.

#### Supplemental S4: Embeddings Generation Computational Cost

Table S4.1: Computational cost to obtain sequence embeddings for each protein family. The time/sequence increase with sequence length and dataset length. The longest dataset to process was GFP sequences and took less 35 minutes to obtain with one standard NVIDIA A6000 GPU and around 13h00 with a CPU.

| **Protein Family** | **Total seq** | **Seq lenght** | **GPU** | | **CPU** | |
|---|---|---|---|---|---|---|
| | | | Full time (s) | time/seq (s) | Full time (s) | time/seq (s) |
| IGPS | 4488 | 221 | 99 | 0.022 | 2226 | 0.496 |
| GFP | 93925 | 234 | 2066 | 0.022 | 47056 | 0.501 |
| SARS-CoV-2 Spike | 14158 | 1253 | 1741 | 0.123 | 40506 | 2.861 |

#### Supplemental S5: Cross-validation Strategies Comparison - Random Splitting vs Grouped $\beta$-Strand Splitting.

Table S5.1: IGPS fitness translocation results. Comparison between random splitting cross-validation and grouped $\beta$-strands splitting cross-validation.

| | | | Target training sizes | | | | | | |
|---|---|---|---|---|---|---|---|---|---|
| **Grouping** | **Target** | **Homolog** | 45 | 75 | 108 | 148 | 196 | 252 | 318 |
| random | SsIGPS | | 0.404 | 0.472 | 0.534 | 0.578 | 0.610 | 0.642 | 0.667 |
| | SsIGPS | TtIGPS | 0.608 | 0.625 | 0.639 | 0.657 | 0.671 | 0.686 | 0.702 |
| $\beta$-strands | SsIGPS | | 0.395 | 0.474 | 0.529 | 0.572 | 0.614 | 0.645 | 0.663 |
| | SsIGPS | TtIGPS | 0.518 | 0.551 | 0.587 | 0.607 | 0.635 | 0.659 | 0.674 |
| random | TmIGPS | | 0.227 | 0.366 | 0.430 | 0.496 | 0.533 | 0.568 | 0.601 |
| | TmIGPS | SsIGPS | 0.489 | 0.518 | 0.548 | 0.577 | 0.595 | 0.617 | 0.640 |
| $\beta$-strands | TmIGPS | | 0.221 | 0.365 | 0.425 | 0.490 | 0.528 | 0.563 | 0.597 |
| | TmIGPS | SsIGPS | 0.414 | 0.448 | 0.489 | 0.542 | 0.567 | 0.597 | 0.615 |
| random | TtIGPS | | 0.416 | 0.487 | 0.546 | 0.591 | 0.623 | 0.661 | 0.690 |
| | TtIGPS | SsIGPS | 0.569 | 0.593 | 0.625 | 0.650 | 0.673 | 0.697 | 0.721 |
| $\beta$-strands | TtIGPS | | 0.411 | 0.492 | 0.542 | 0.590 | 0.623 | 0.658 | 0.687 |
| | TtIGPS | SsIGPS | 0.471 | 0.527 | 0.576 | 0.607 | 0.639 | 0.673 | 0.699 |

#### Supplemental S6: SARS-CoV-2 Spike protein Cell Entry, ESM-1v, SVR - Lasso - RF, No-Selection

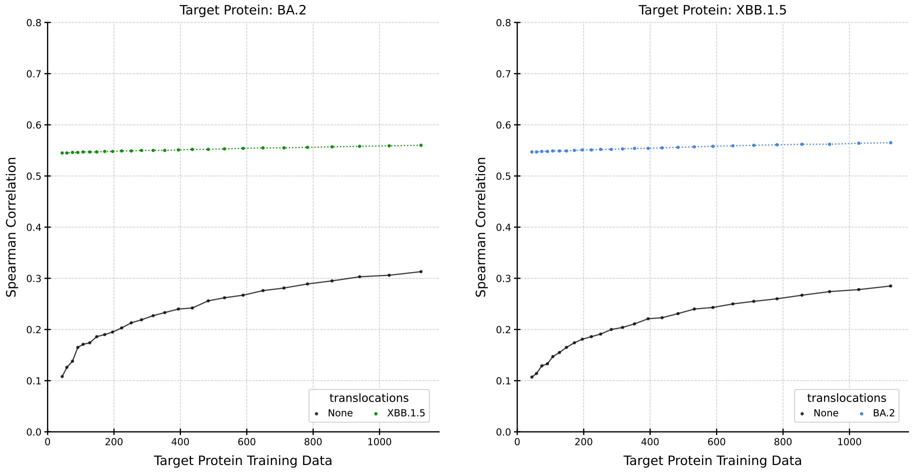


Figure S6.1: SARS-CoV-2 spike protein, ESM-1v pLM, and SVR predictor.

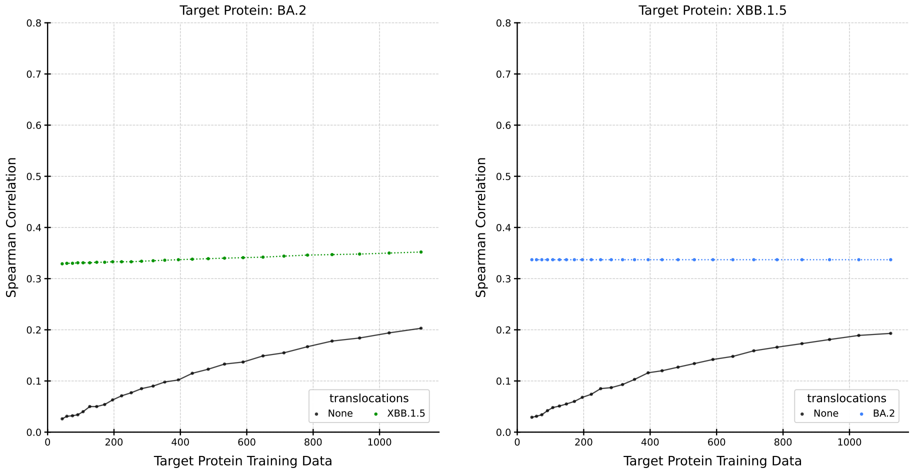


Figure S6.2: SARS-CoV-2 spike protein, ESM-1v pLM, and Lasso predictor.

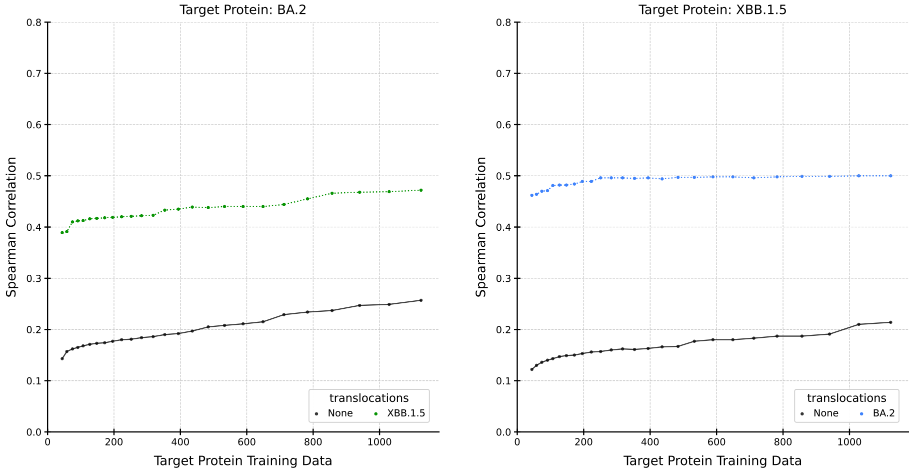


Figure S6.3: SARS-CoV-2 spike protein, ESM-1v pLM, and RF predictor.

#### Supplemental S7: SARS-CoV-2 Spike protein ACE2 Bindings, ESM-1v, SVR - Lasso - RF, No-Selection

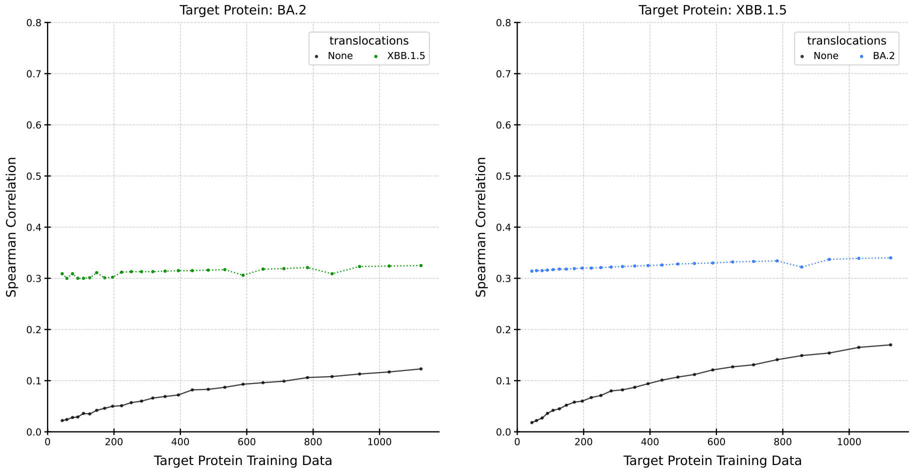


Figure S7.1: SARS-CoV-2 spike protein ACE2 Binding, ESM-1v pLM, and SVR predictor.

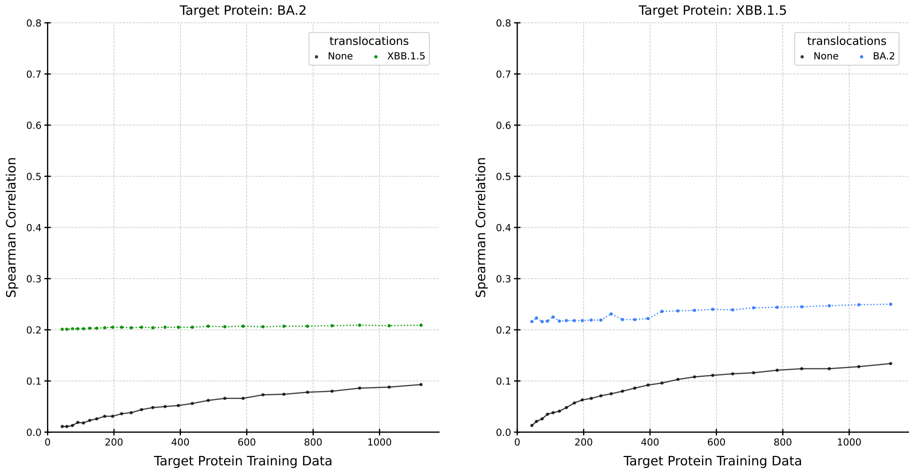


Figure S7.2: SARS-CoV-2 spike protein ACE2 Binding, ESM-1v pLM, and Lasso predictor.

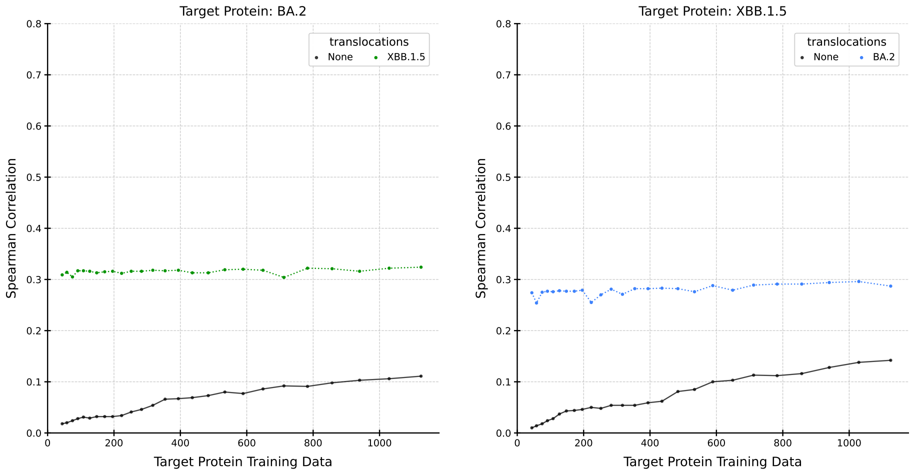


Figure S7.3: SARS-CoV-2 spike protein ACE2 Binding, ESM-1v pLM, and RF predictor.

### Supplemental S8: IGPS, ESM-1v, SVR - Lasso - RF, No-Selection

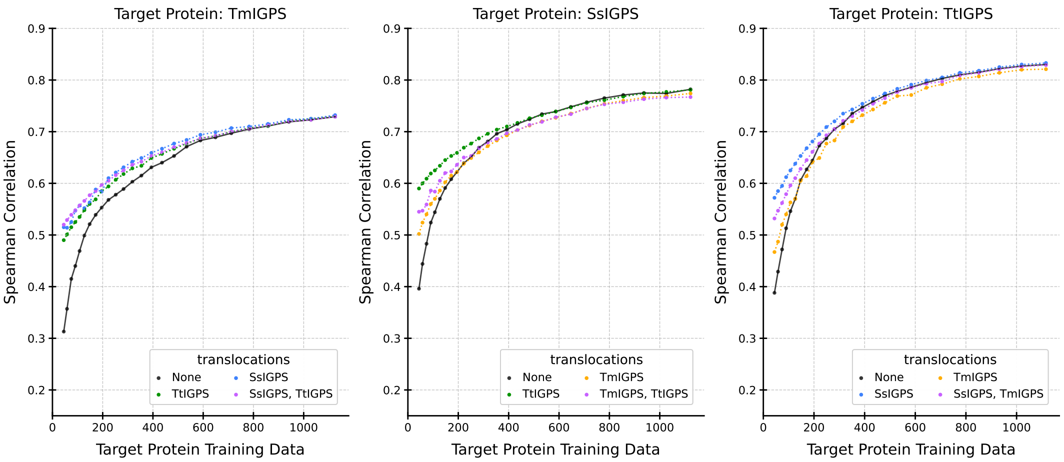


Figure S8.1: IGPS orthologs, ESM-1v pLM, and SVR predictor.

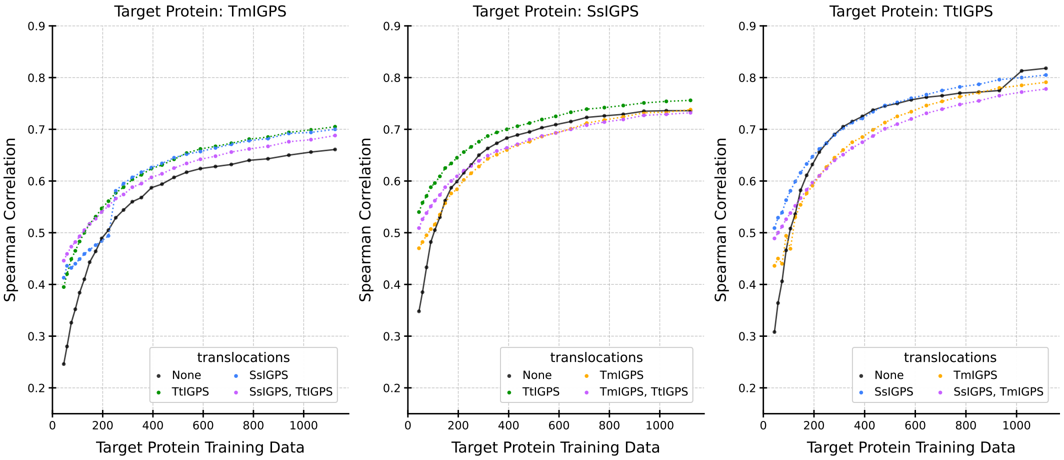


Figure S8.2: IGPS orthologs, ESM-1v pLM, and lasso predictor.

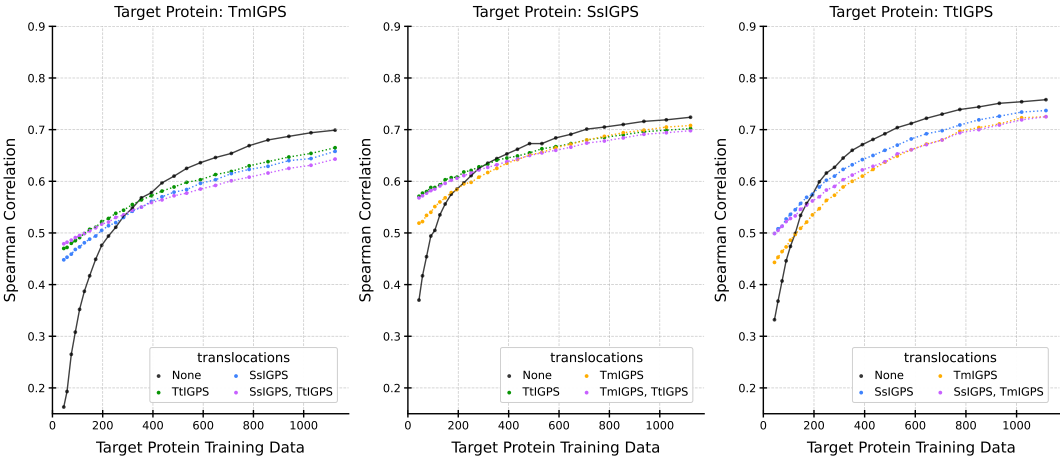


Figure S8.3: IGPS orthologs, ESM-1v pLM, and RF predictor.

### Supplemental S9: GFP, ESM-1v, SVR - Lasso - RF, No-Selection

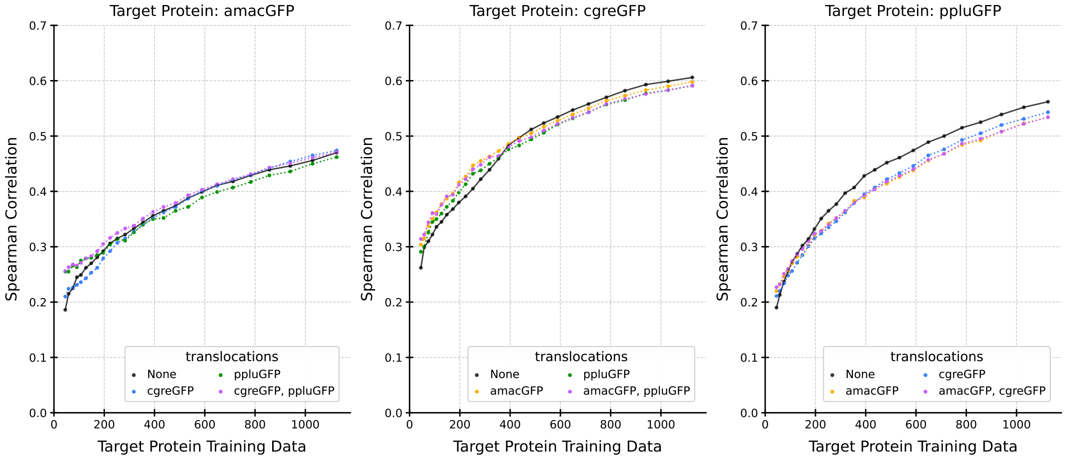


Figure S9.1: GFP orthologs, ESM-1v pLM, and SVR predictor.

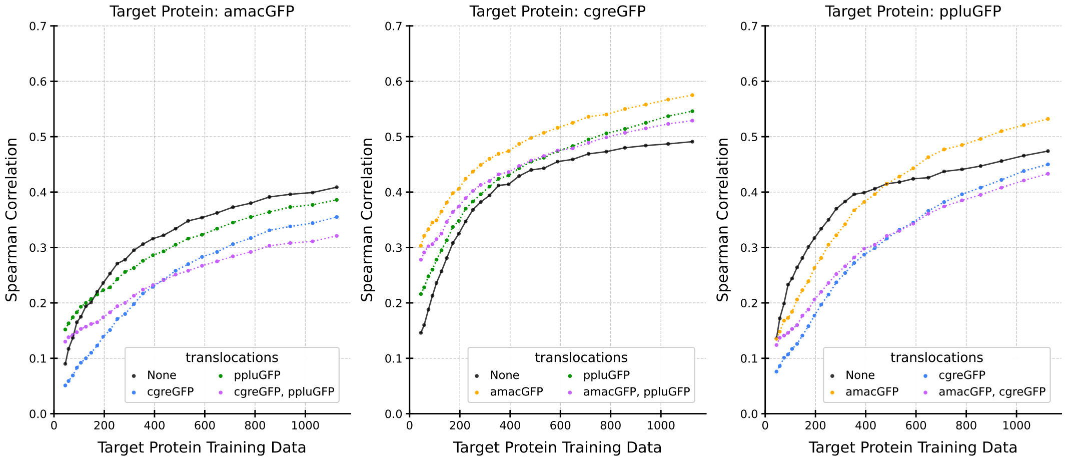


Figure S9.2: GFP orthologs, ESM-1v pLM, and Lasso predictor.

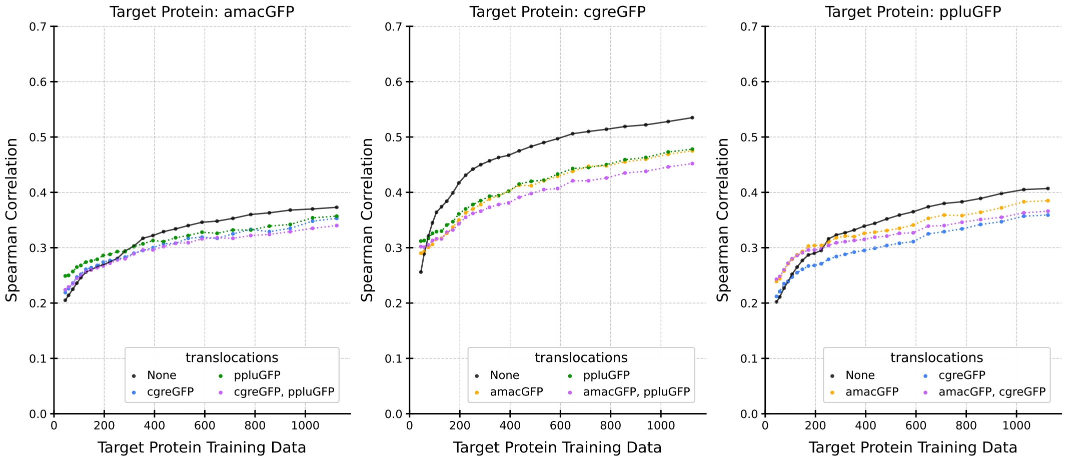


Figure S9.3: GFP orthologs, ESM-1v pLM, and RF predictor.

### Supplemental S10: SARS-CoV-2 Spike protein Cell Entry, ESM2, Lasso - SVR - RF, No-Selection

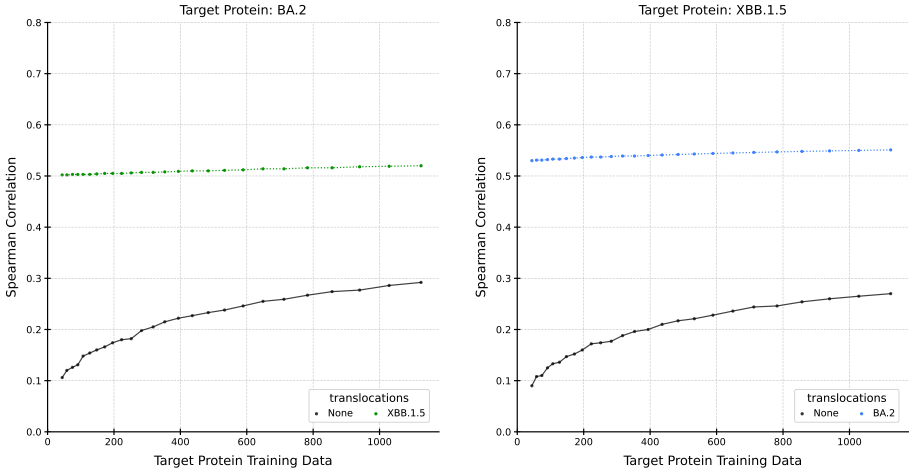


Figure S10.1: SARS-CoV-2 spike protein Cell Entry, ESM2 pLM, and SVR predictor.

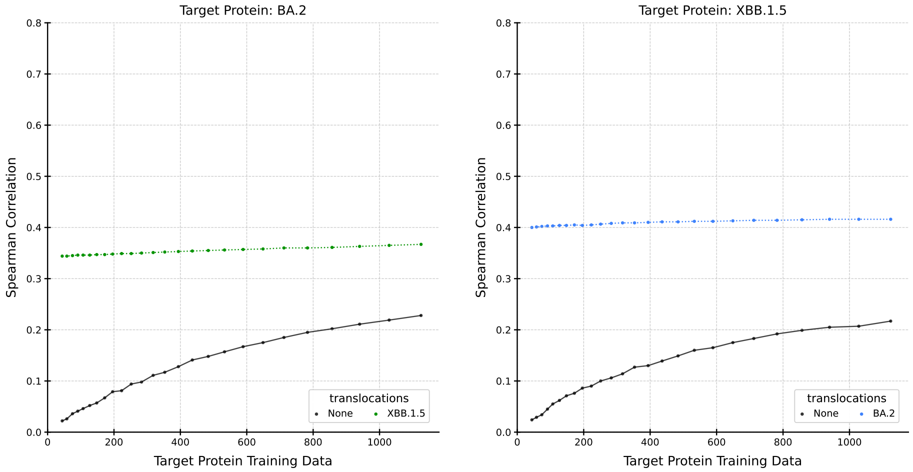


Figure S10.2: SARS-CoV-2 spike protein Cell Entry, ESM2 pLM, and Lasso predictor.

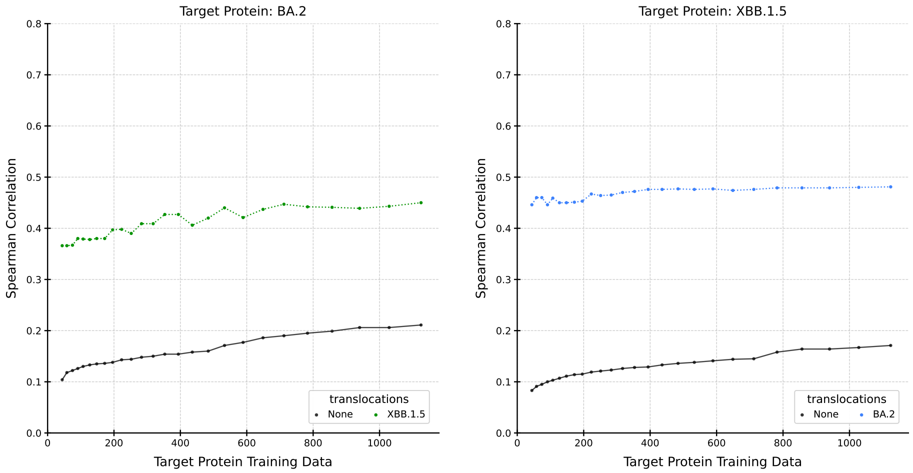


Figure S10.3: SARS-CoV-2 spike protein Cell Entry, ESM2 pLM, and RF predictor.

#### Supplemental S11: SARS-CoV-2 Spike protein ACE2 Binding, ESM2, Lasso - SVR - RF, No-Selection

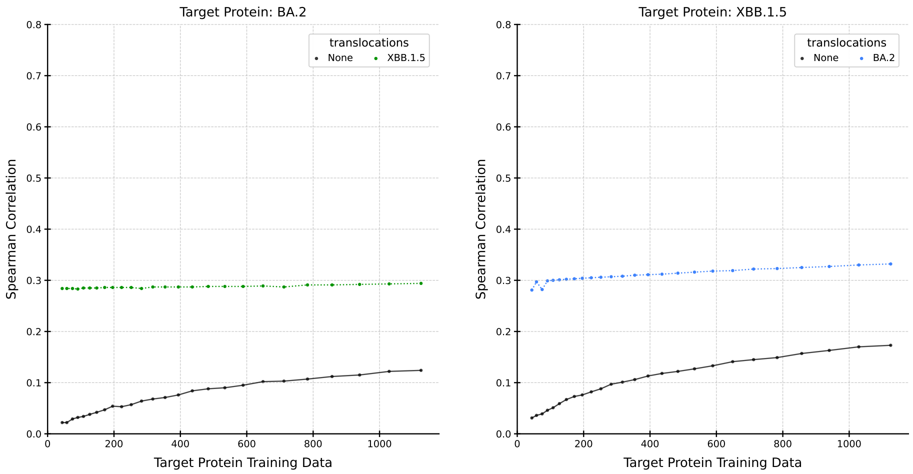


Figure S11.1: SARS-CoV-2 spike protein ACE2 Binding, ESM2 pLM, and SVR predictor.

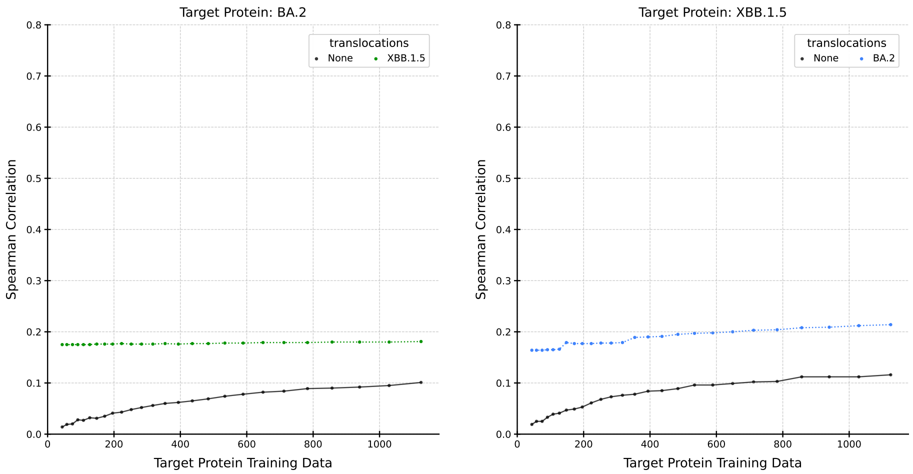


Figure S11.2: SARS-CoV-2 spike protein ACE2 Binding, ESM2 pLM, and Lasso predictor.

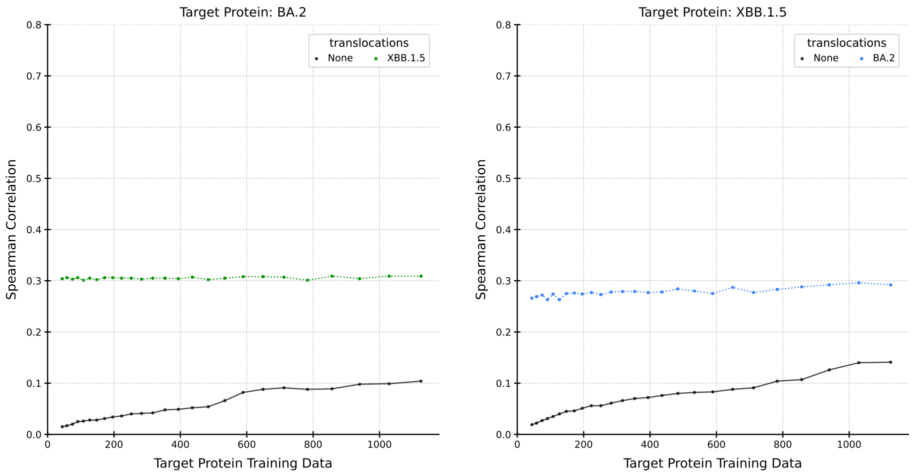


Figure S11.3: SARS-CoV-2 spike protein ACE2 Binding, ESM2 pLM, and RF predictor.

#### Supplemental S12: IGPS, ESM2, Lasso - SVR - RF, No-Selection

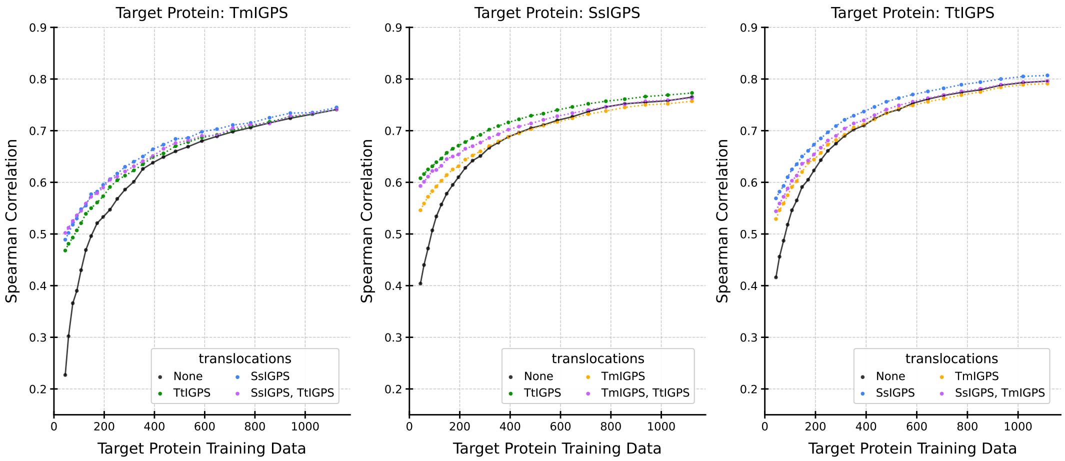


Figure S12.1: IGPS, ESM2 pLM, and SVR predictor.

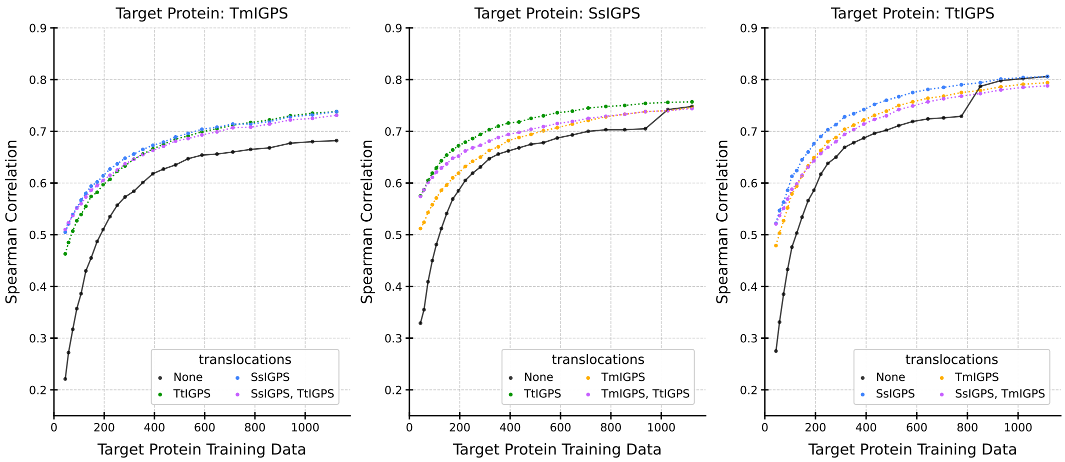


Figure S12.2: IGPS, ESM2 pLM, and Lasso predictor.

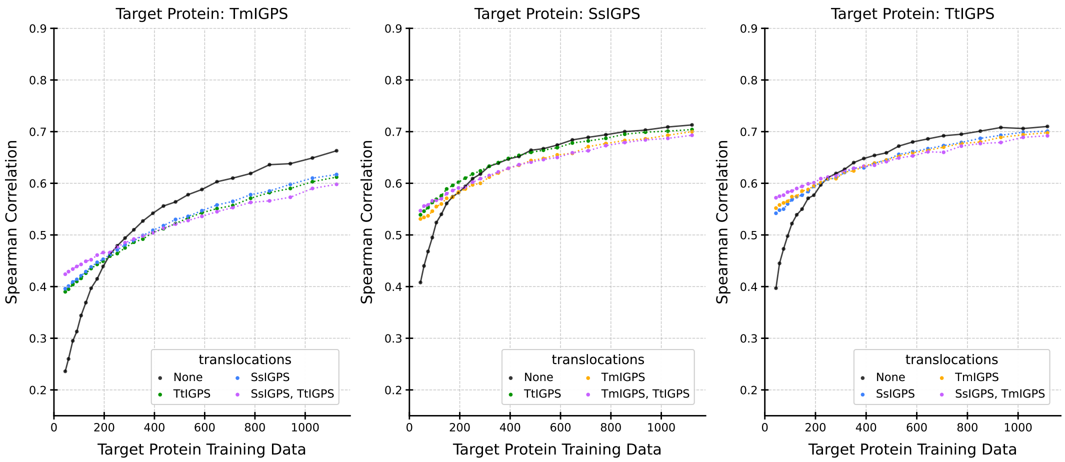


Figure S12.3: IGPS, ESM2 pLM, and RF predictor.

### Supplemental S13: GFP, ESM2, SVR - Lasso - RF, No-Selection

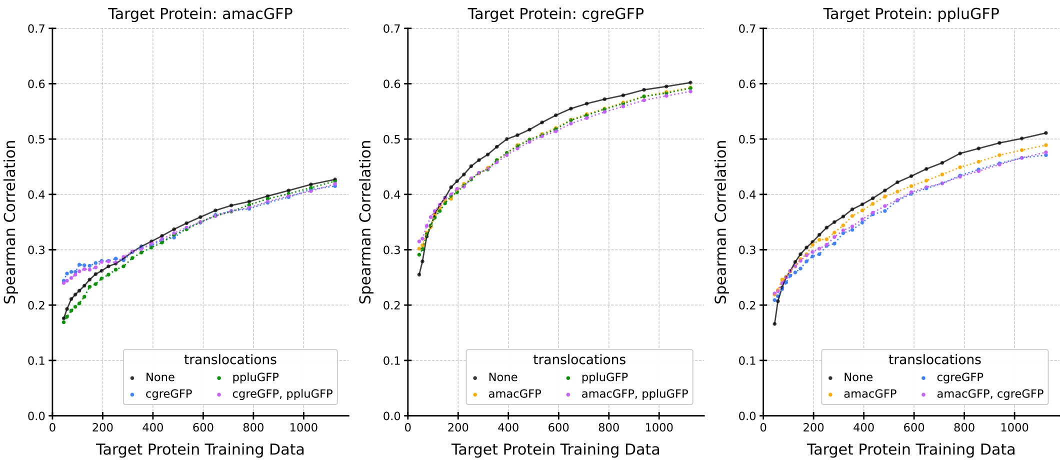


Figure S13.1: GFP, ESM2 pLM, and SVR predictor.

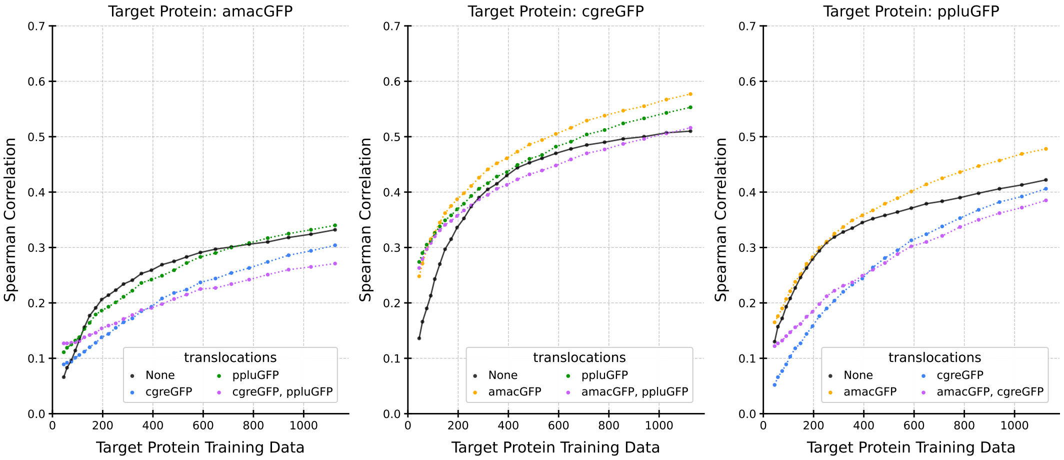


Figure S13.2: GFP, ESM2 pLM, and Lasso predictor.

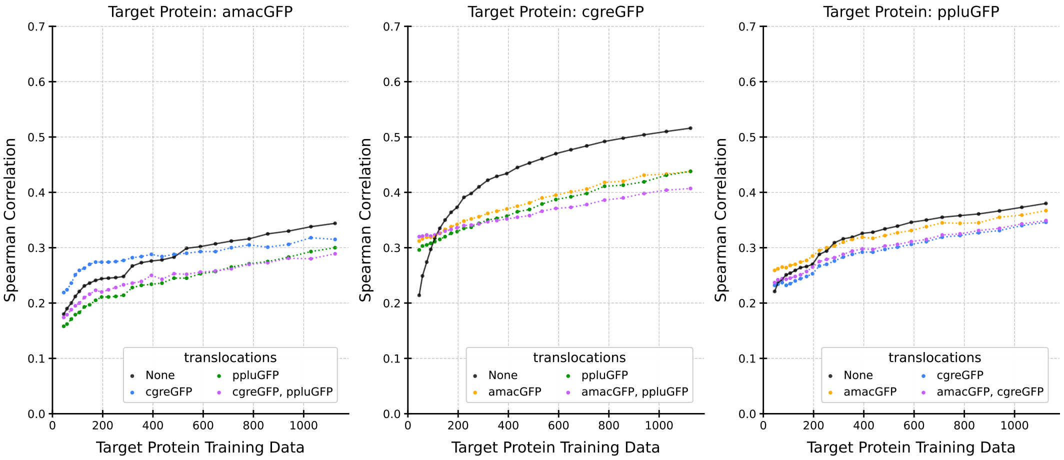


Figure S13.3: GFP, ESM2 pLM, and RF predictor.

#### Supplemental S14: SARS-CoV-2 Spike protein Cell Entry, ESM-1v, SVR - Lasso - RF, Statistical-Greedy

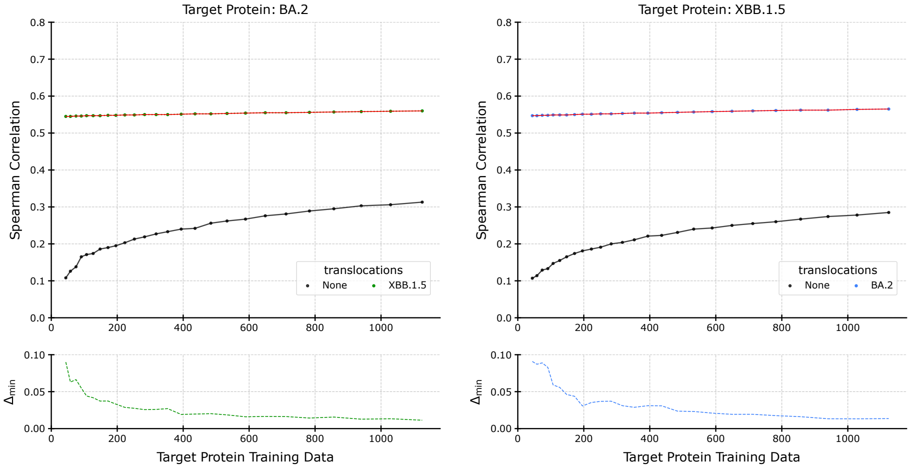


Figure S14.1: SARS-CoV-2 spike protein, ESM-1v pLM, and SVR predictor.

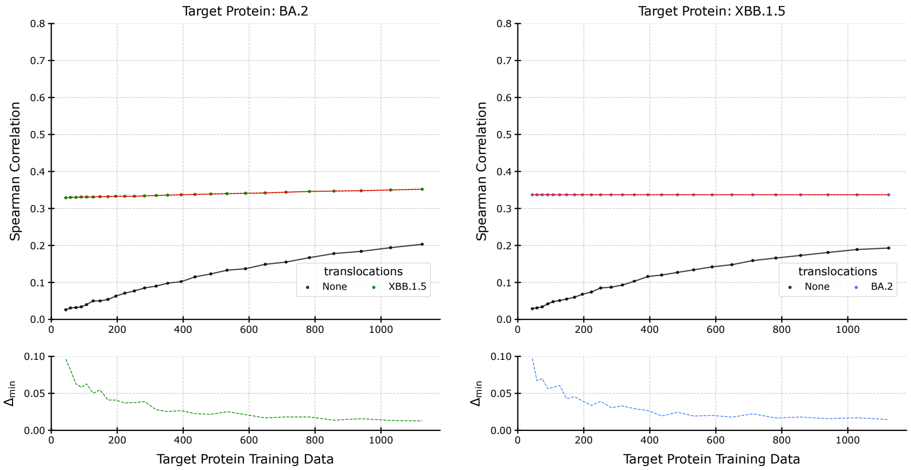


Figure S14.2: SARS-CoV-2 spike protein, ESM-1v pLM, and Lasso predictor.

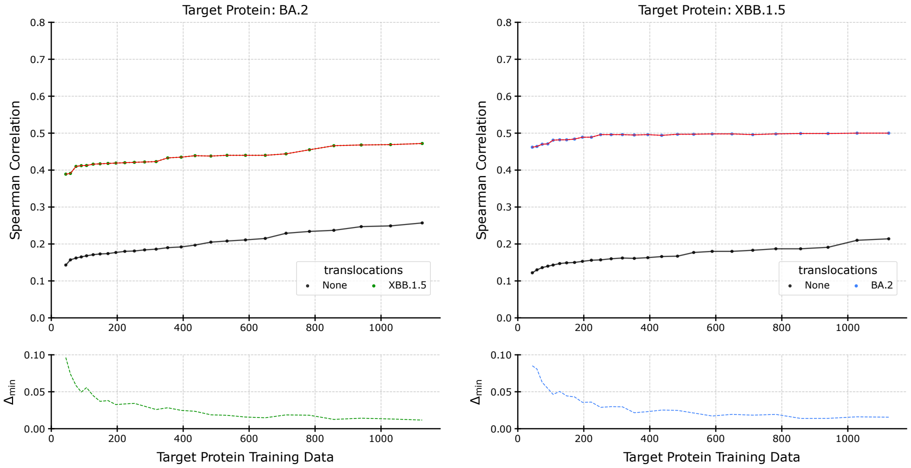


Figure S14.3: SARS-CoV-2 spike protein, ESM-1v pLM, and RF predictor.

#### Supplemental S15: SARS-CoV-2 Spike protein ACE2 Binding, ESM-1v, SVR - Lasso - RF, Statistical-Greedy

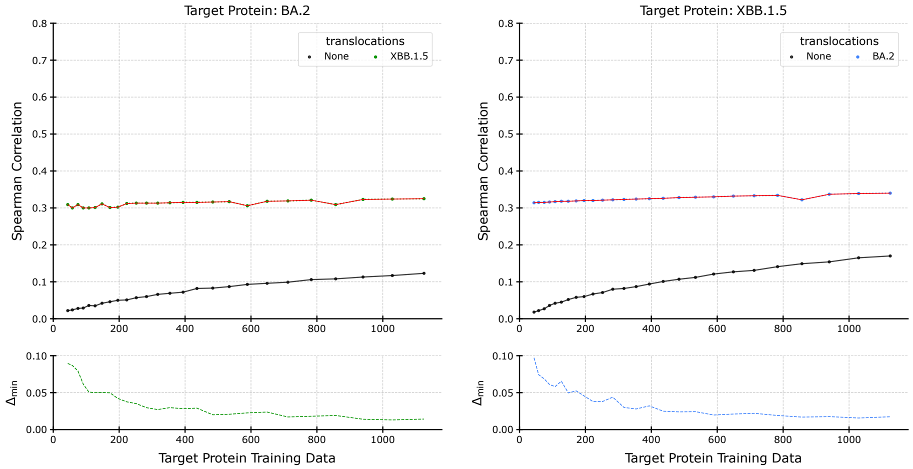


Figure S15.1: SARS-CoV-2 spike protein ACE2 Binding, ESM-1v pLM, and SVR predictor.

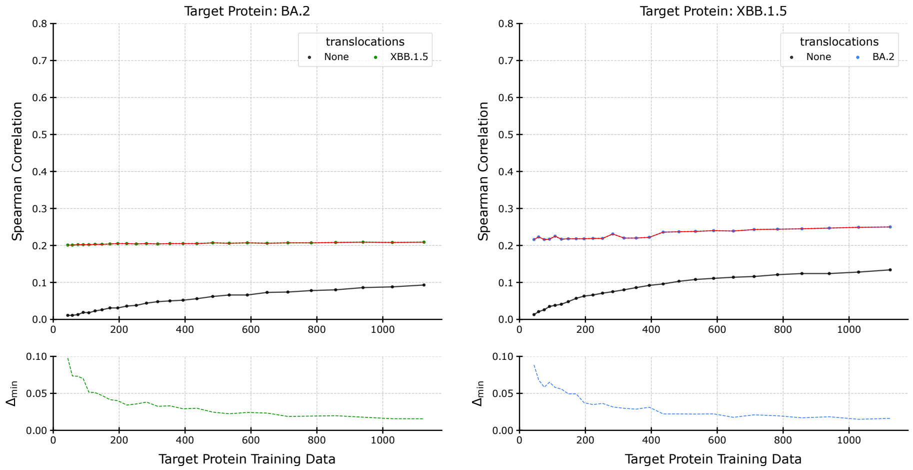


Figure S15.2: SARS-CoV-2 spike protein ACE2 Binding, ESM-1v pLM, and Lasso predictor.

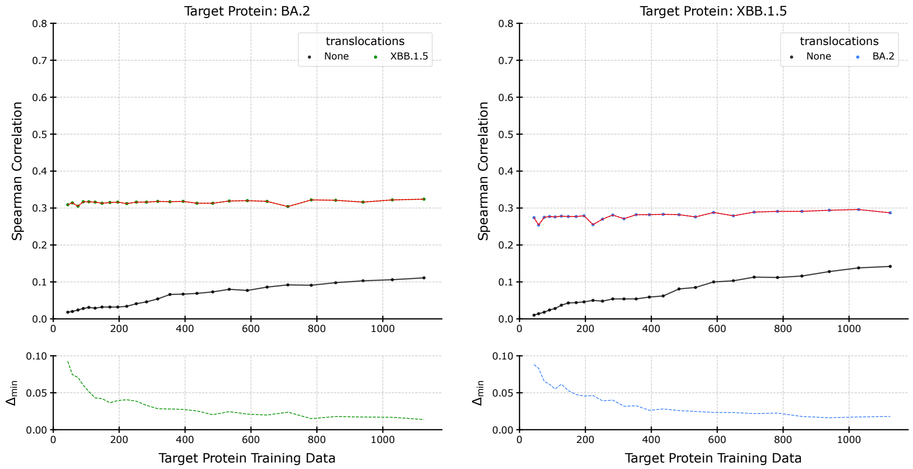


Figure S15.3: SARS-CoV-2 spike protein ACE2 Binding, ESM-1v pLM, and RF predictor.

#### Supplemental S16: IGPS, ESM-1v, SVR - Lasso - RF, Statistical-Greedy

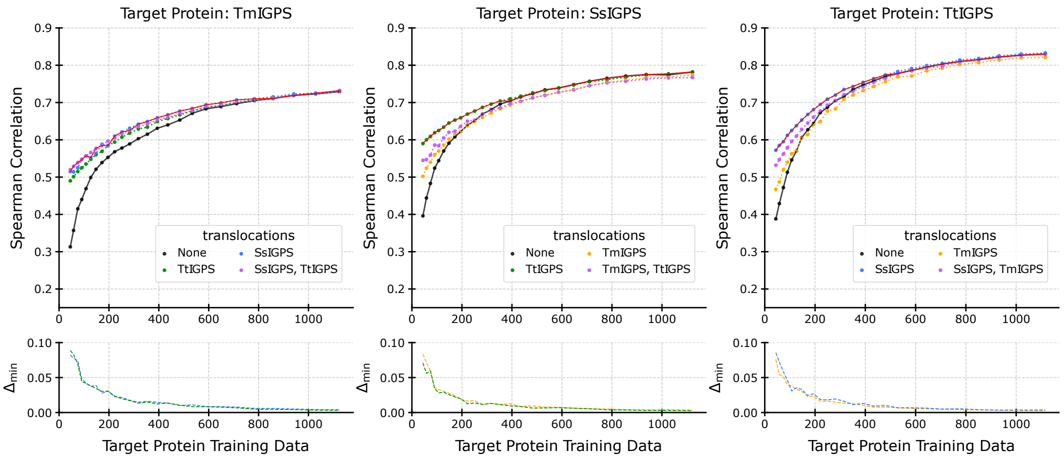


Figure S16.1: IGPS, ESM-1v pLM, and SVR predictor.

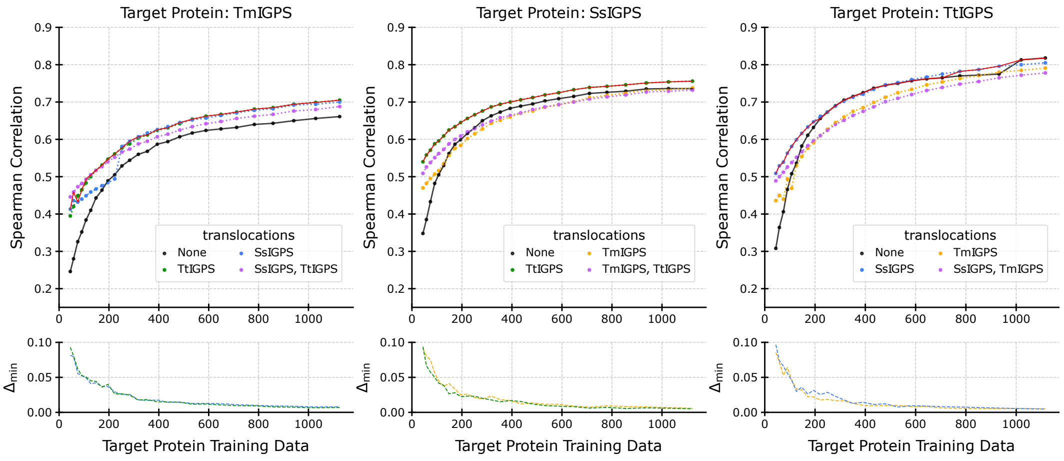


Figure S16.2: IGPS, ESM-1v pLM, and Lasso predictor.

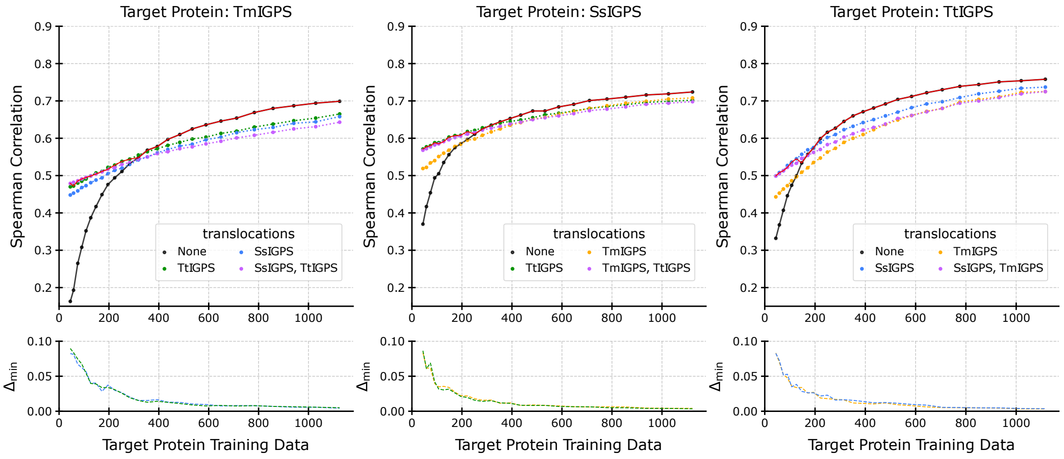


Figure S16.3: IGPS, ESM-1v pLM, and RF predictor.

#### Supplemental S17: GFP, ESM-1v, SVR - Lasso - RF, Statistical-Greedy

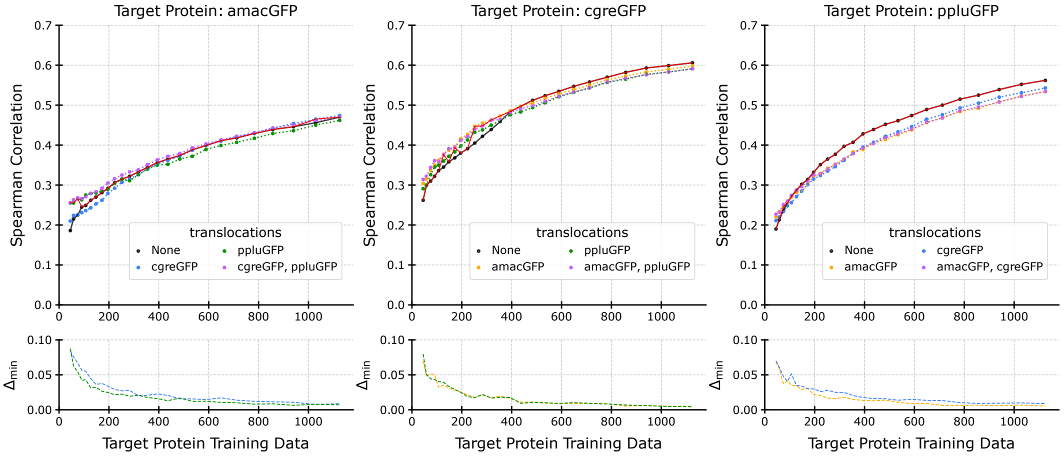


Figure S17.1: GFP, ESM-1v pLM, and SVR predictor.

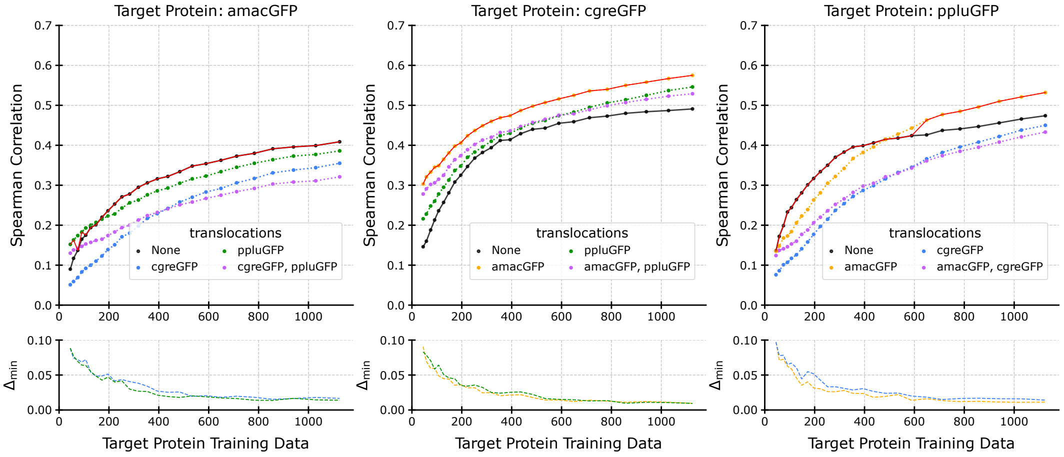


Figure S17.2: GFP, ESM-1v pLM, and Lasso predictor.

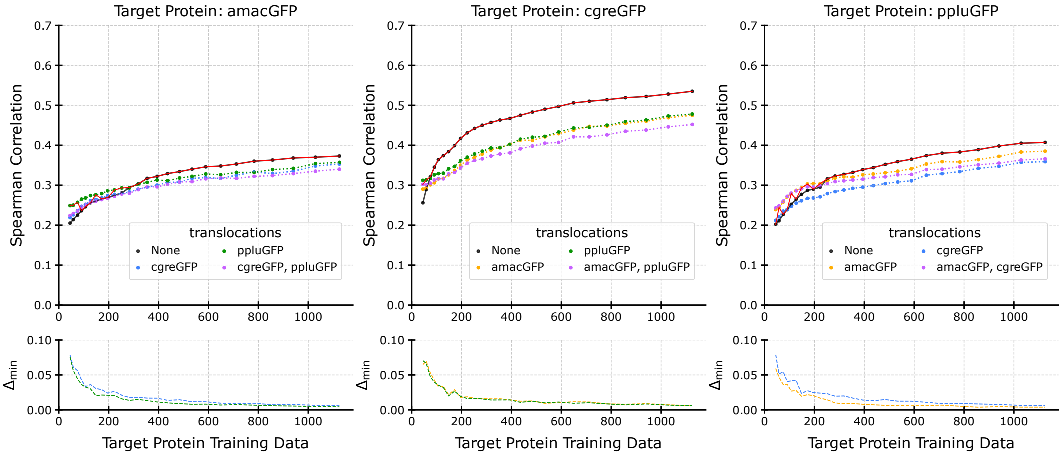


Figure S17.3: GFP, ESM-1v pLM, and RF predictor.

### Supplemental S18: SARS-CoV-2 Spike protein Cell Entry, ESM2, SVR - Lasso - RF, Statistical-Greedy

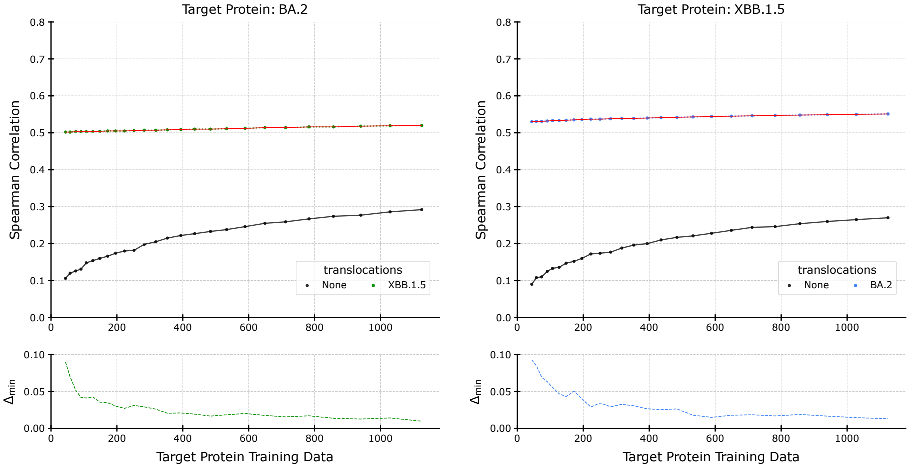


Figure S18.1: SARS-CoV-2 spike protein, ESM2 pLM, and SVR predictor.

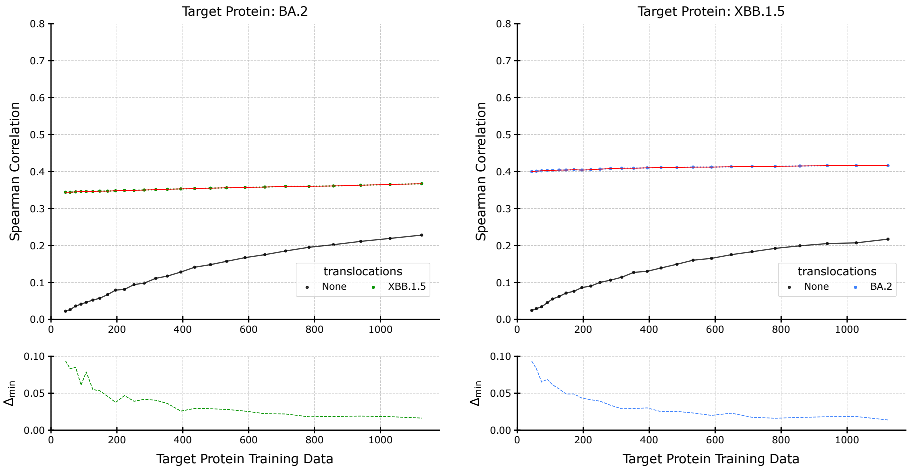


Figure S18.2: SARS-CoV-2 spike protein, ESM2 pLM, and Lasso predictor.

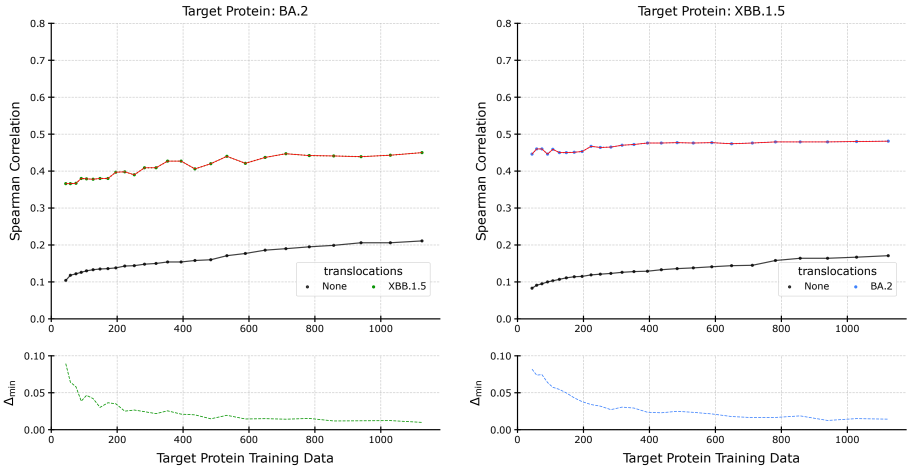


Figure S18.3: SARS-CoV-2 spike protein, ESM2 pLM, and RF predictor.

#### Supplemental S19: SARS-CoV-2 Spike protein ACE2 Binding, ESM2, SVR - Lasso - RF, Statistical-Greedy

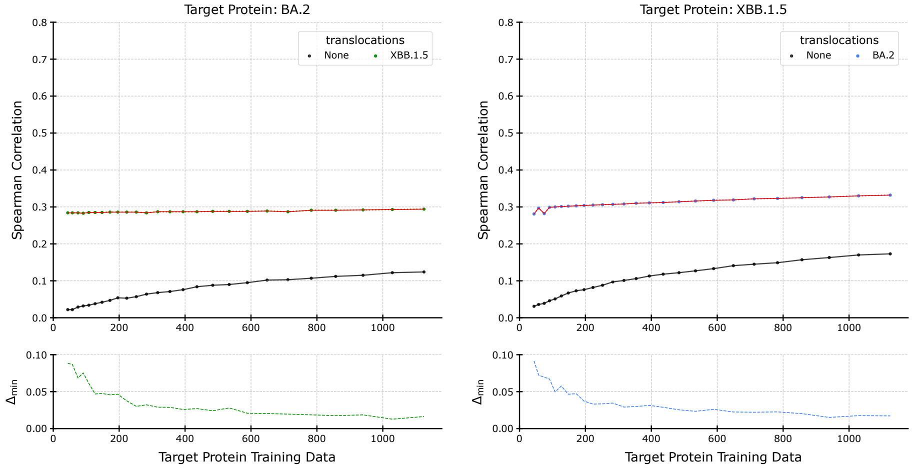


Figure S19.1: SARS-CoV-2 spike protein ACE2 Binding, ESM2 pLM, and SVR predictor.

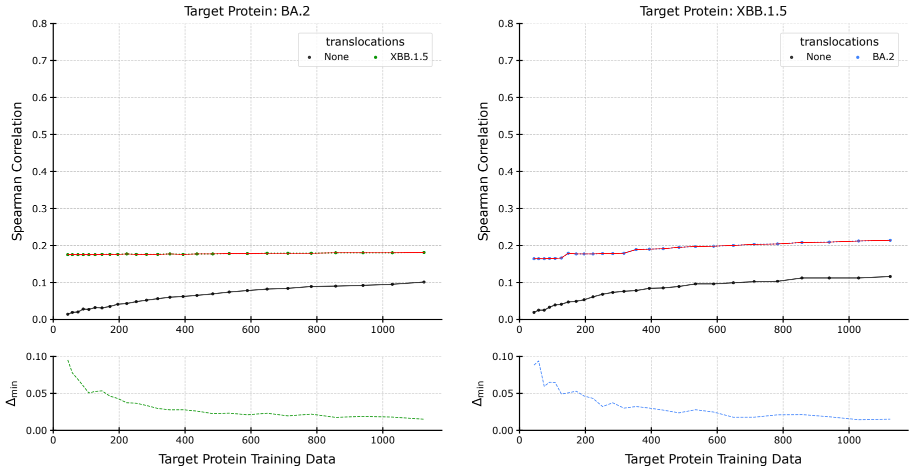


Figure S19.2: SARS-CoV-2 spike protein ACE2 Binding, ESM2 pLM, and Lasso predictor.

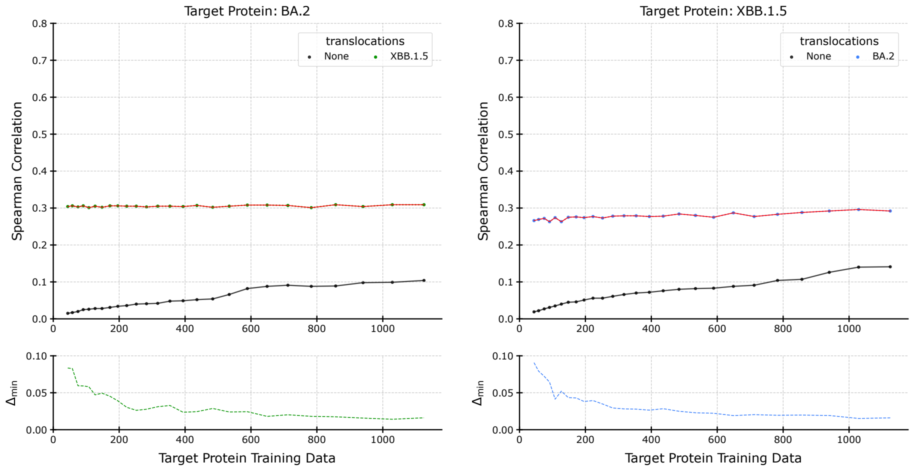


Figure S19.3: SARS-CoV-2 spike protein ACE2 Binding, ESM2 pLM, and RF predictor.

### Supplemental S20: IGPS, ESM2, SVR - Lasso - RF, Statistical-Greedy

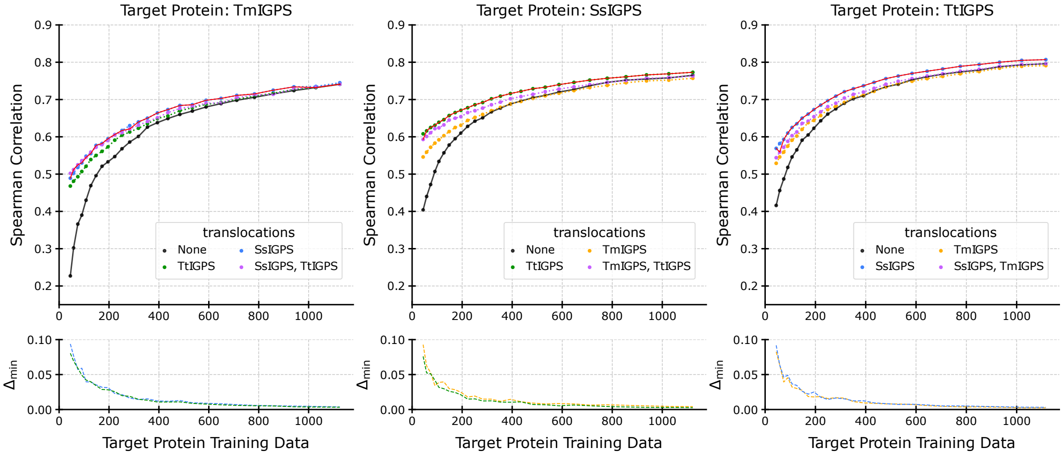


Figure S20.1: IGPS, ESM2 pLM, and SVR predictor.

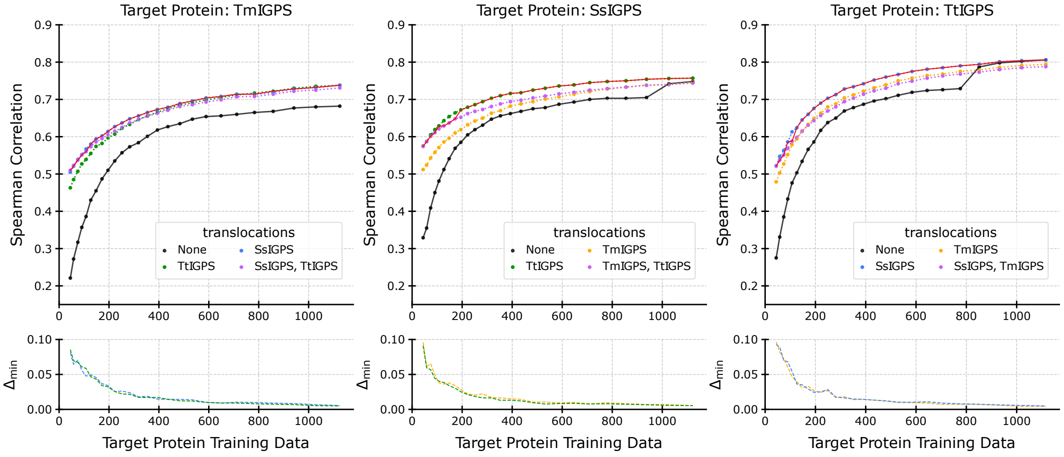


Figure S20.2: IGPS, ESM2 pLM, and Lasso predictor.

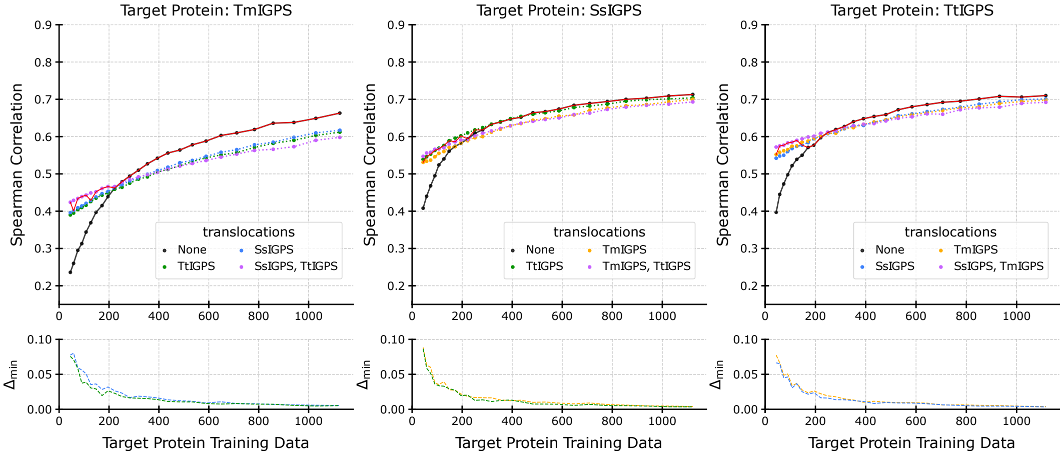


Figure S20.3: IGPS, ESM2 pLM, and RF predictor.

### Supplemental S21: GFP, ESM2, SVR - Lasso - RF, Statistical-Greedy

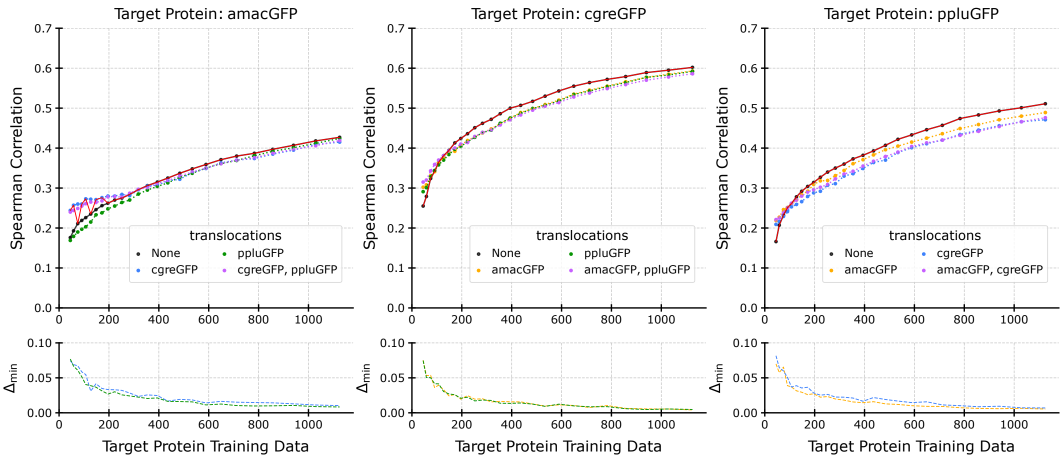


Figure S21.1: GFP, ESM2 pLM, and SVR predictor.

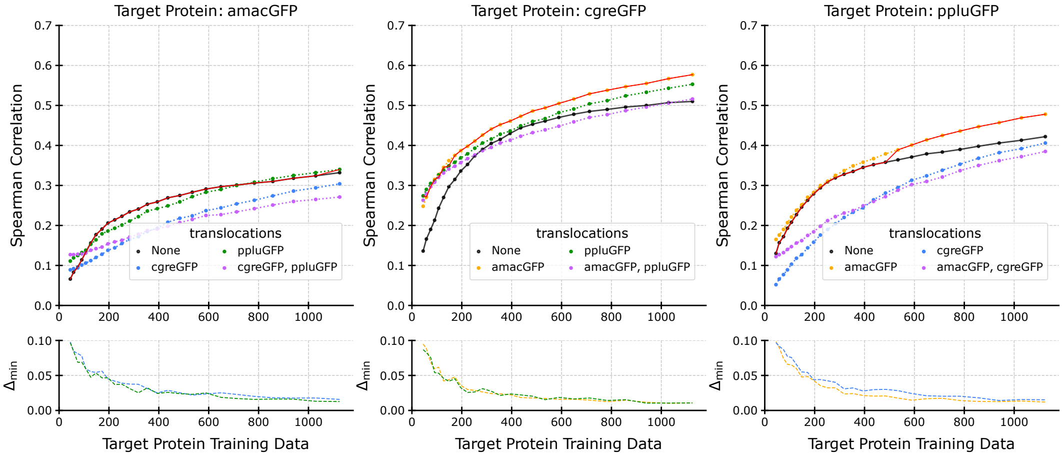


Figure S21.2: GFP, ESM2 pLM, and Lasso predictor.

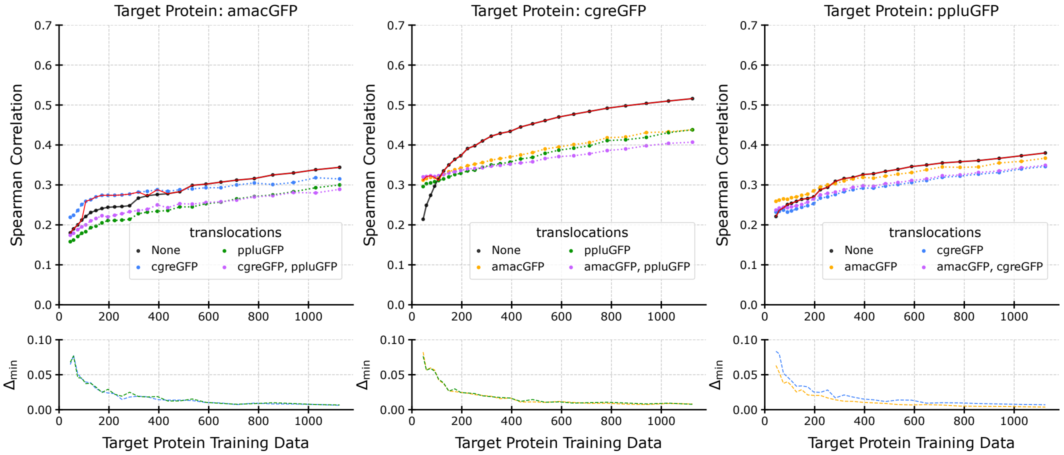


Figure S21.3: GFP, ESM2 pLM, and RF predictor.

#### Supplemental S22: SARS-CoV-2 Spike protein Cell Entry, ESM-1v, SVR, Statistical-Greedy - Individual-Greedy - Individual-Select

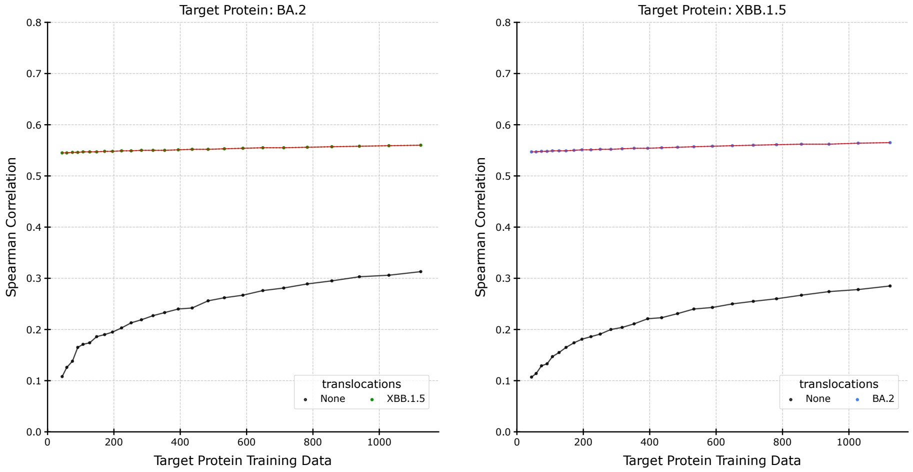


Figure S22.1: SARS-CoV-2 Spike protein Cell Entry, ESM-1v pLM, and SVR predictor.

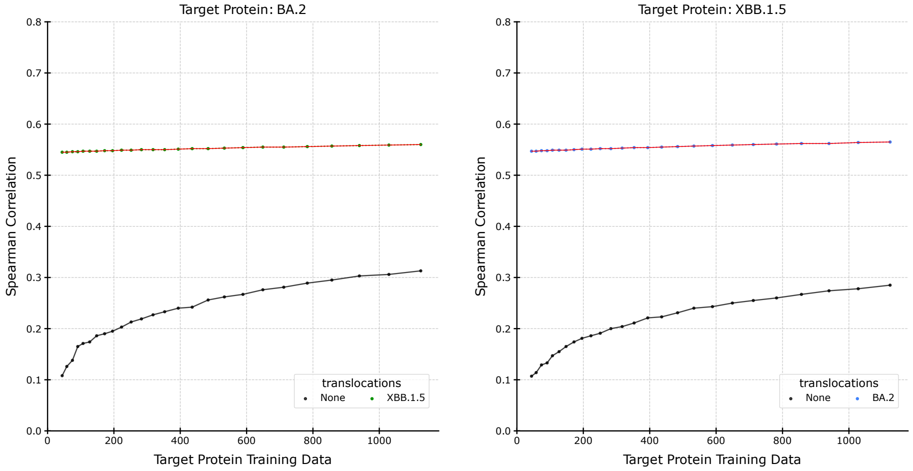


Figure S22.2: SARS-CoV-2 Spike protein Cell Entry, ESM-1v pLM, and SVR predictor.

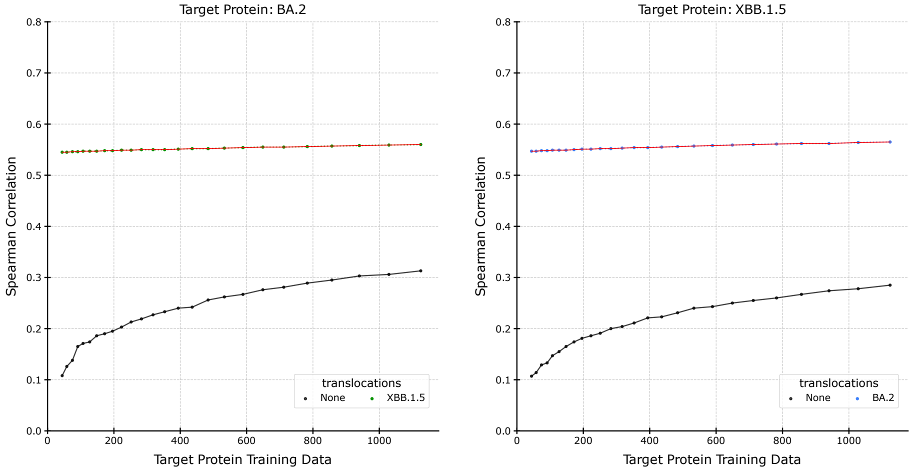


Figure S22.3: SARS-CoV-2 Spike protein Cell Entry, ESM-1v pLM, and SVR predictor.

#### Supplemental S23: SARS-CoV-2 Spike protein Cell Entry, ESM-1v, Lasso, Statistical-Greedy - Individual-Greedy - Individual-Select

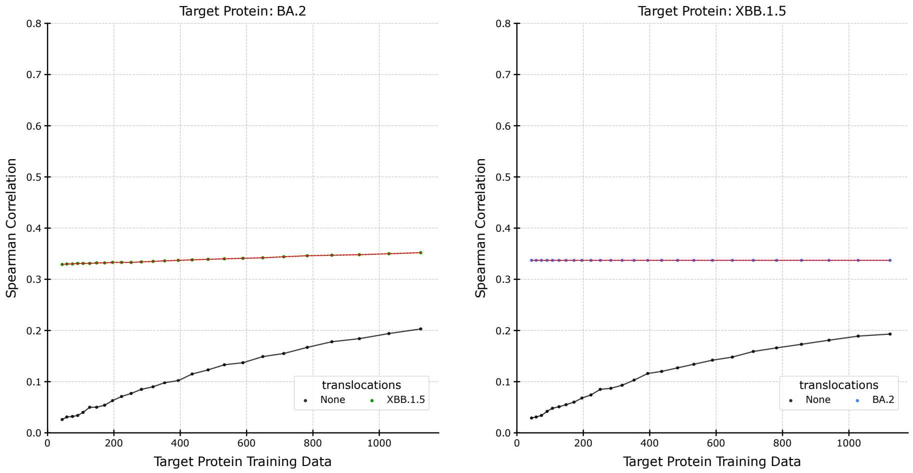


Figure S23.1: SARS-CoV-2 Spike protein Cell Entry, ESM-1v pLM, and Lasso predictor.

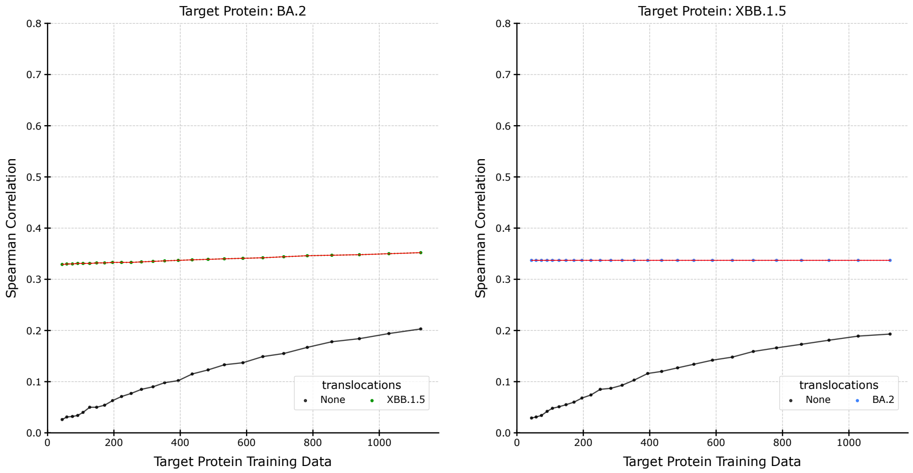


Figure S23.2: SARS-CoV-2 Spike protein Cell Entry, ESM-1v pLM, and Lasso predictor.

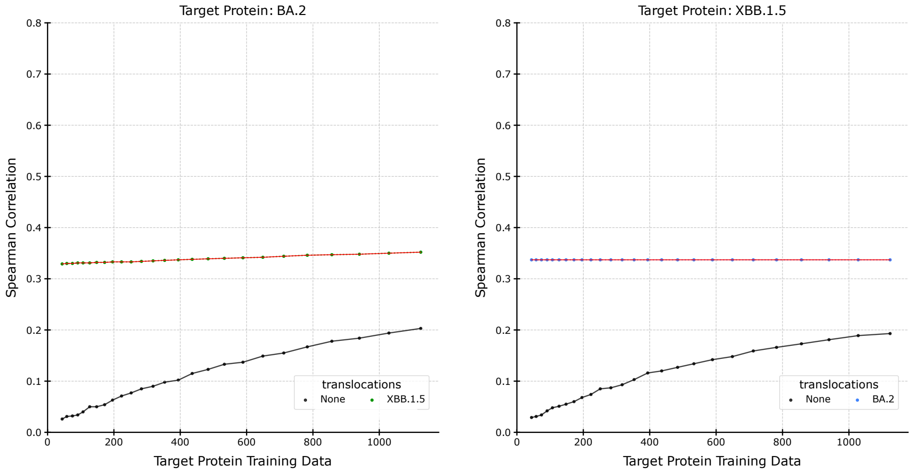


Figure S23.3: SARS-CoV-2 Spike protein Cell Entry, ESM-1v pLM, and Lasso predictor.

#### Supplemental S24: SARS-CoV-2 Spike protein Cell Entry, ESM-1v, RF, Statistical-Greedy - Individual-Greedy - Individual-Select

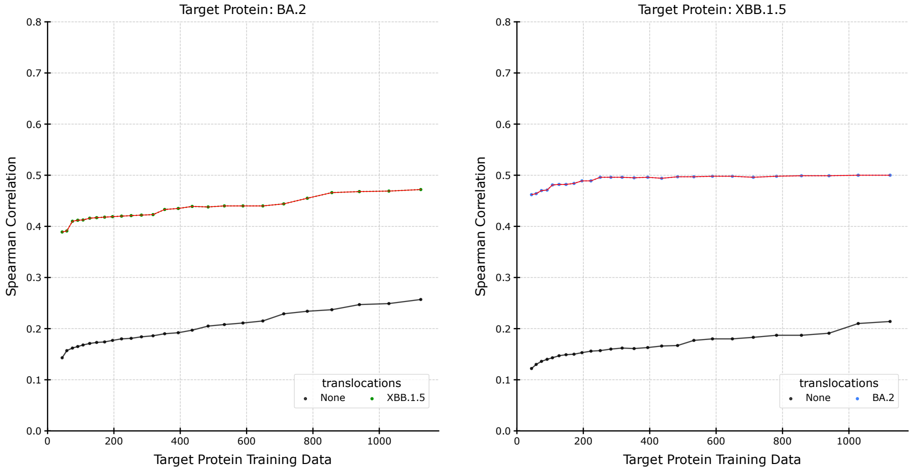


Figure S24.1: SARS-CoV-2 Spike protein Cell Entry, ESM-1v pLM, and RF predictor.

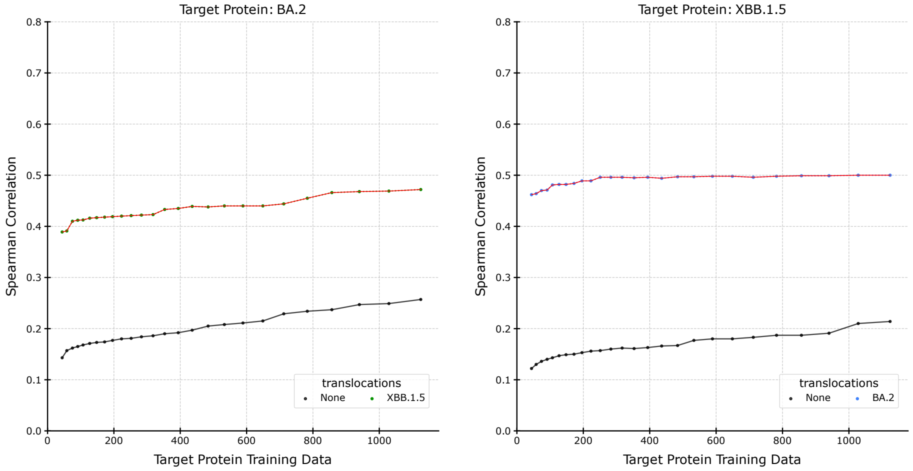


Figure S24.2: SARS-CoV-2 Spike protein Cell Entry, ESM-1v pLM, and RF predictor.

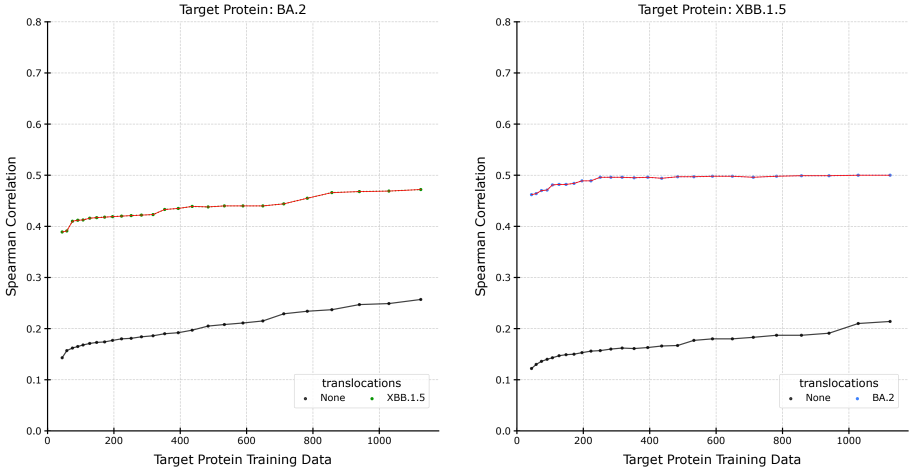


Figure S24.3: SARS-CoV-2 Spike protein Cell Entry, ESM-1v pLM, and RF predictor.

#### Supplemental S25: SARS-CoV-2 Spike protein ACE2 Binding, ESM-1v, SVR, Statistical-Greedy - Individual-Greedy - Individual-Select

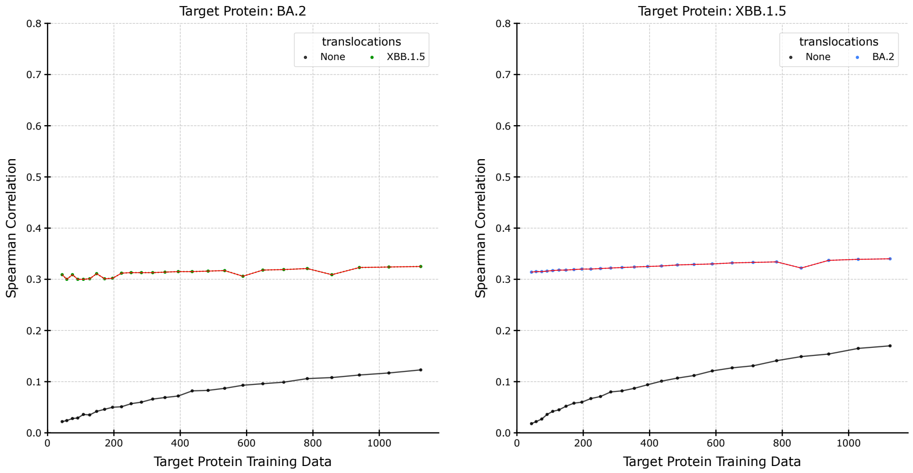


Figure S25.1: SARS-CoV-2 Spike protein ACE2 Binding, ESM-1v pLM, and SVR predictor.

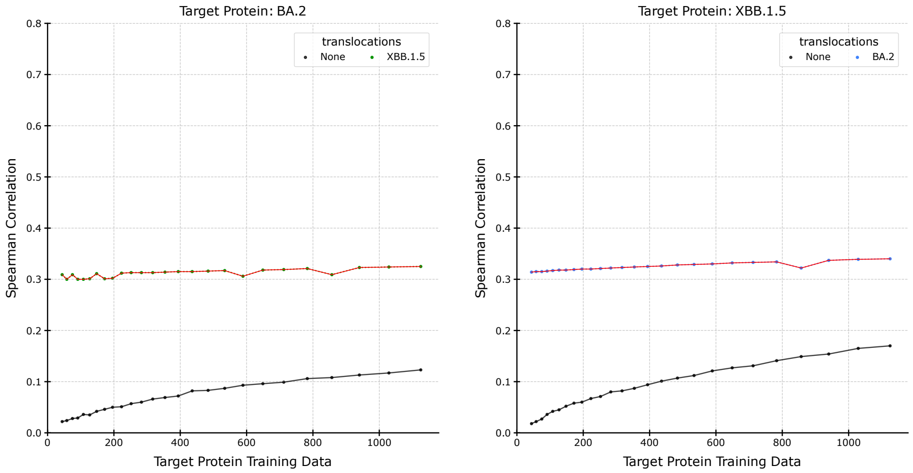


Figure S25.2: SARS-CoV-2 Spike protein ACE2 Binding, ESM-1v pLM, and SVR predictor.

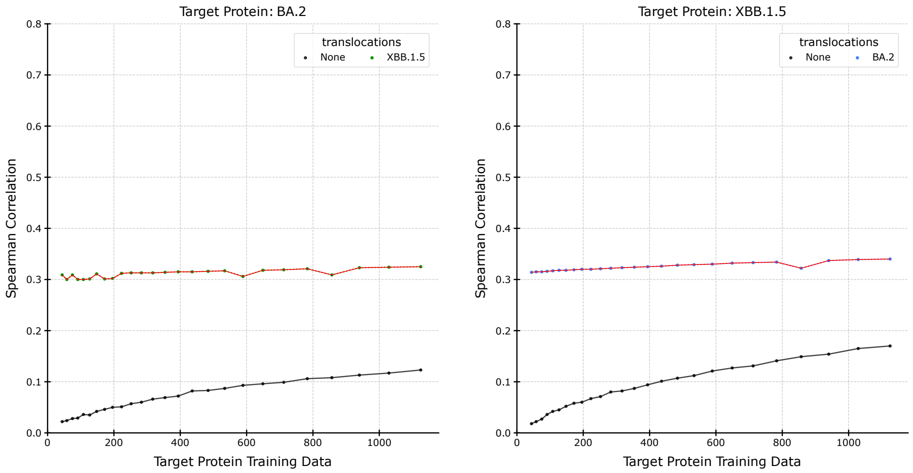


Figure S25.3: SARS-CoV-2 Spike protein ACE2 Binding, ESM-1v pLM, and SVR predictor.

#### Supplemental S26: SARS-CoV-2 Spike protein ACE2 Binding, ESM-1v, Lasso, Statistical-Greedy - Individual-Greedy - Individual-Select

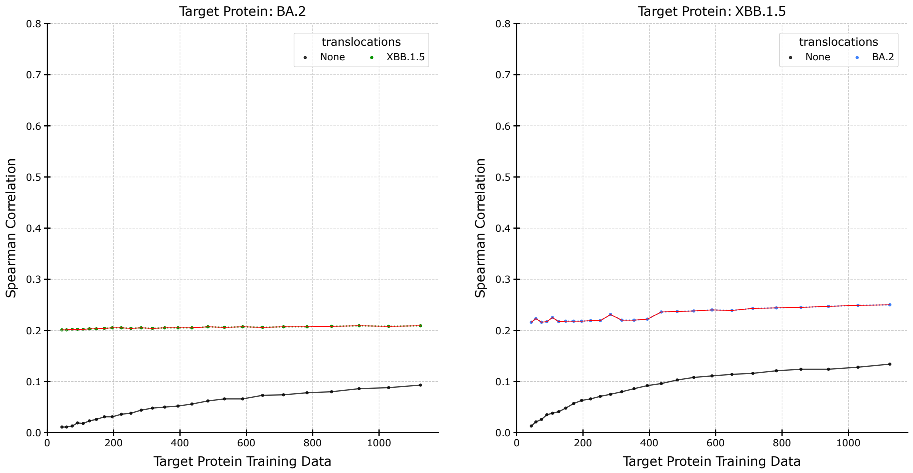


Figure S26.1: SARS-CoV-2 Spike protein ACE2 Binding, ESM-1v pLM, and Lasso predictor.

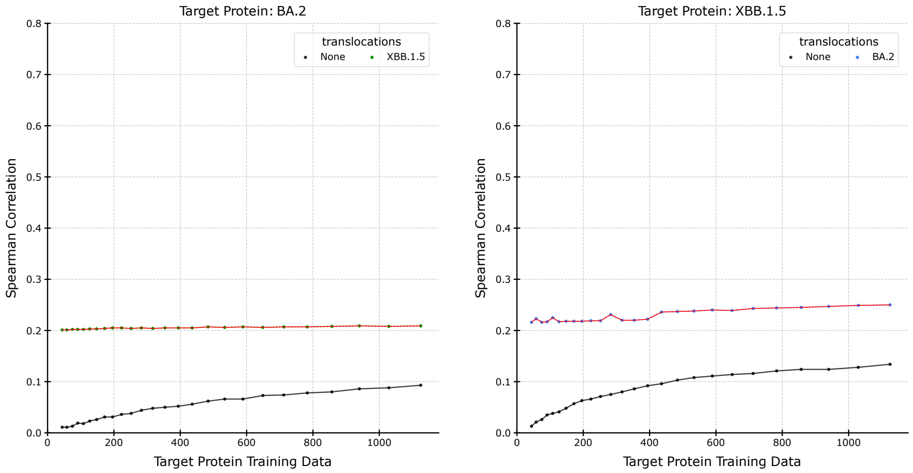


Figure S26.2: SARS-CoV-2 Spike protein ACE2 Binding, ESM-1v pLM, and Lasso predictor.

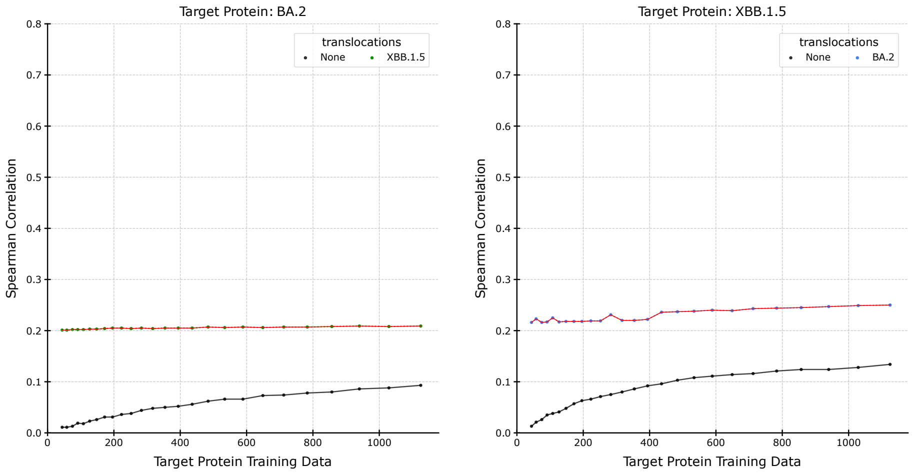


Figure S26.3: SARS-CoV-2 Spike protein ACE2 Binding, ESM-1v pLM, and Lasso predictor.

#### Supplemental S27: SARS-CoV-2 Spike protein ACE2 Binding, ESM-1v, RF, Statistical-Greedy - Individual-Greedy - Individual-Select

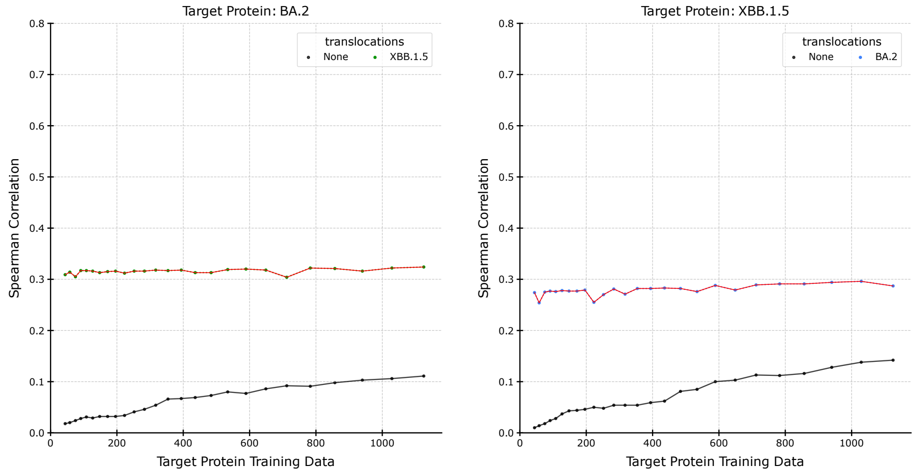


Figure S27.1: SARS-CoV-2 Spike protein ACE2 Binding, ESM-1v pLM, and RF predictor.

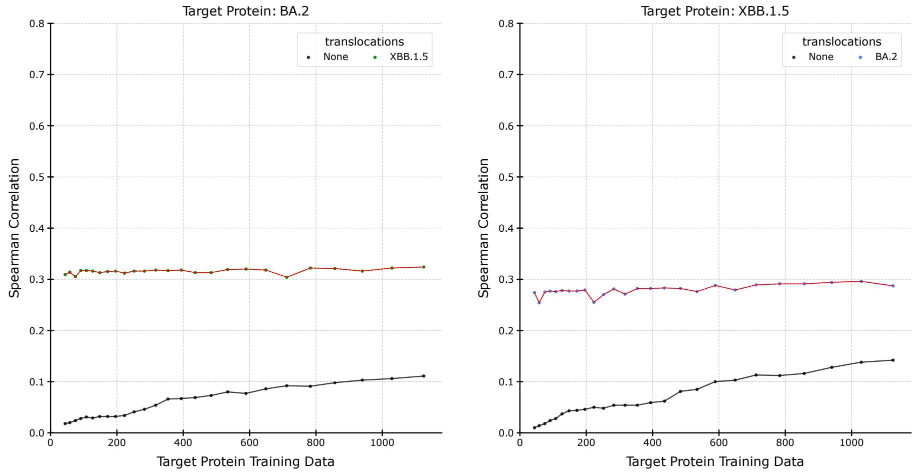


Figure S27.2: SARS-CoV-2 Spike protein ACE2 Binding, ESM-1v pLM, and RF predictor.

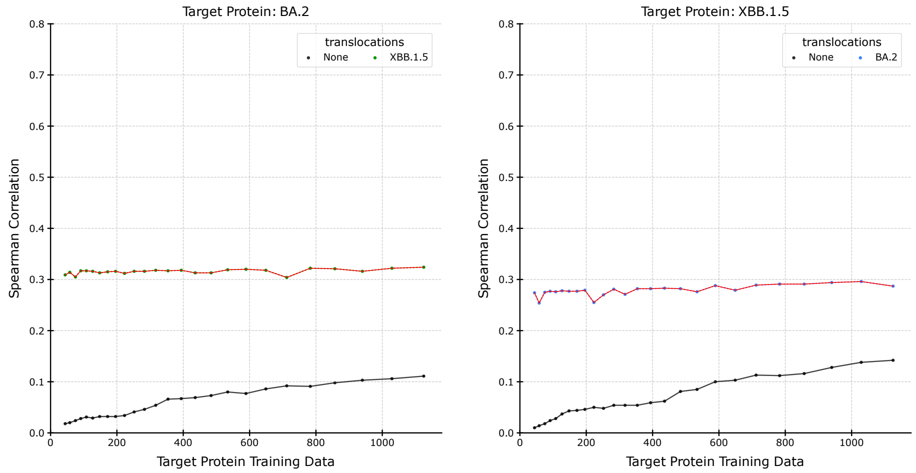


Figure S27.3: SARS-CoV-2 Spike protein ACE2 Binding, ESM-1v pLM, and RF predictor.

#### Supplemental S28: IGPS, ESM-1v, SVR, Statistical-Greedy - Individual-Greedy - Individual-Select

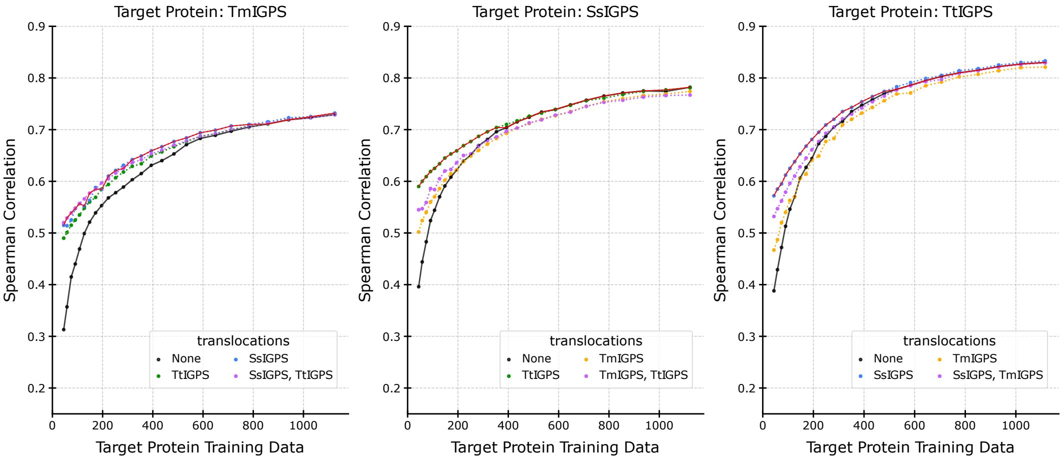


Figure S28.1: IGPS, ESM-1v pLM, and SVR predictor.

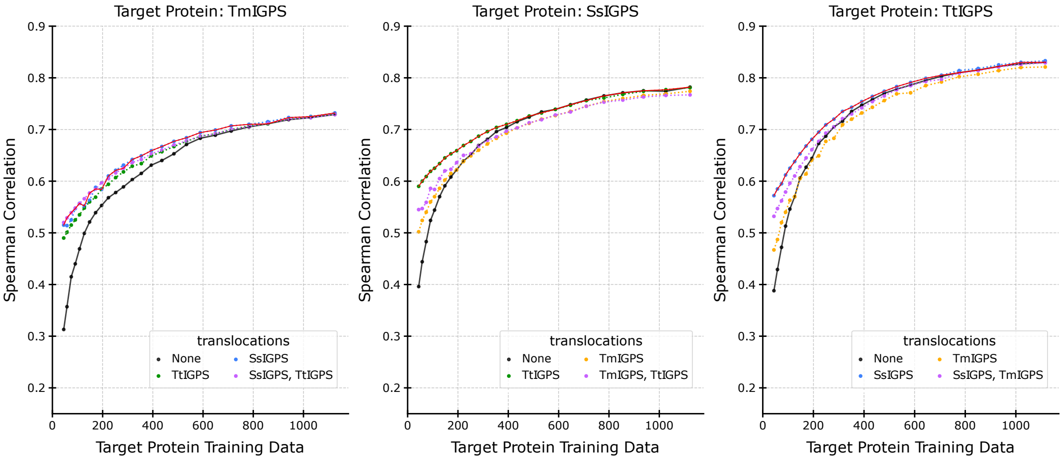


Figure S28.2: IGPS, ESM-1v pLM, and SVR predictor.

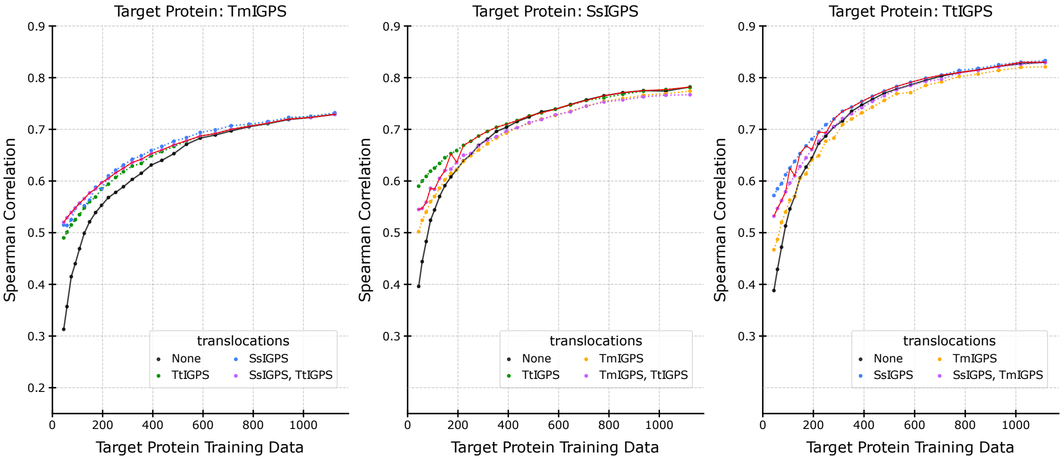


Figure S28.3: IGPS, ESM-1v pLM, and SVR predictor.

### Supplemental S29: IGPS, ESM-1v, Lasso, Statistical-Greedy - Individual-Greedy - Individual-Select

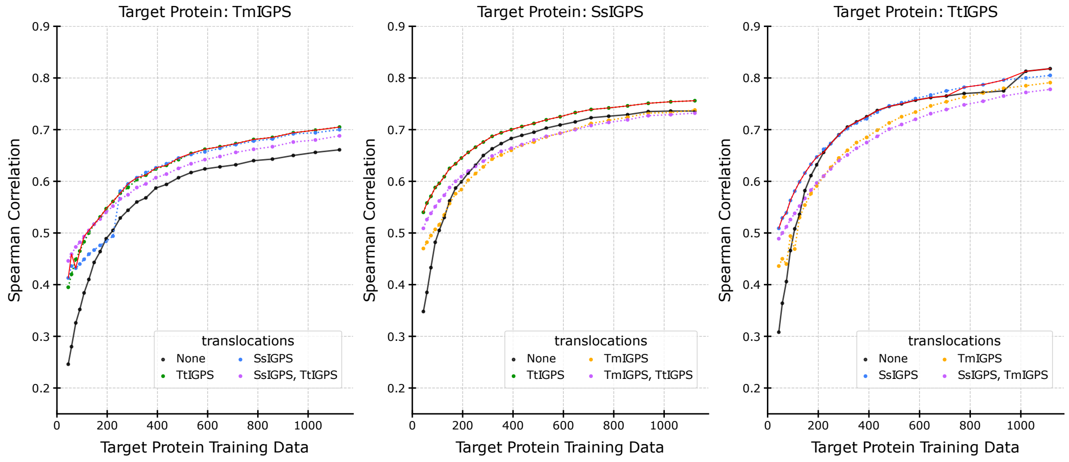


Figure S29.1: IGPS, ESM-1v pLM, and Lasso predictor.

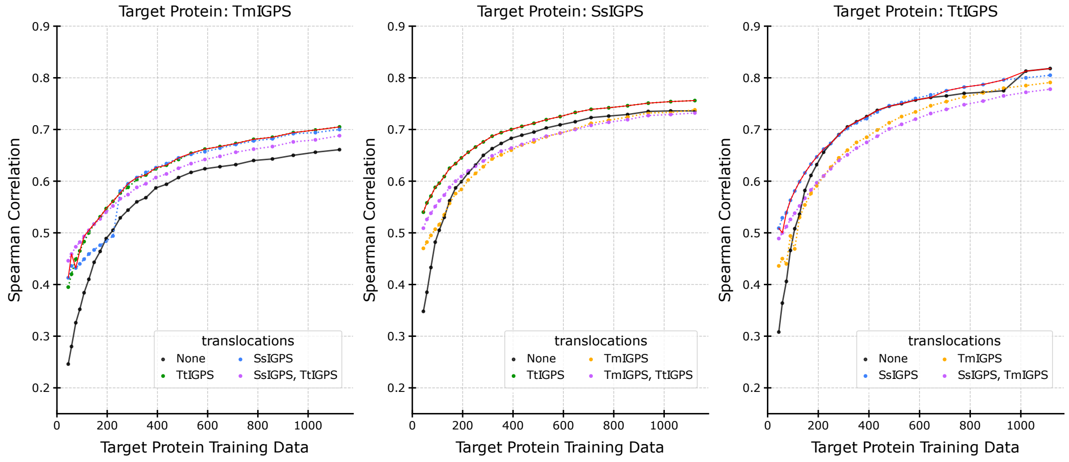


Figure S29.2: IGPS, ESM-1v pLM, and Lasso predictor.

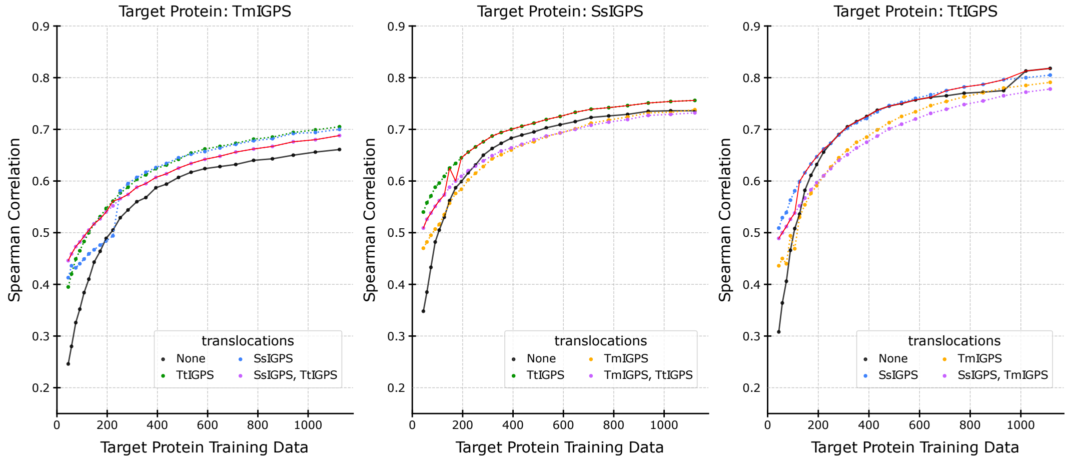


Figure S29.3: IGPS, ESM-1v pLM, and Lasso predictor.

#### Supplemental S30: IGPS, ESM-1v, RF, Statistical-Greedy - Individual-Greedy - Individual-Select

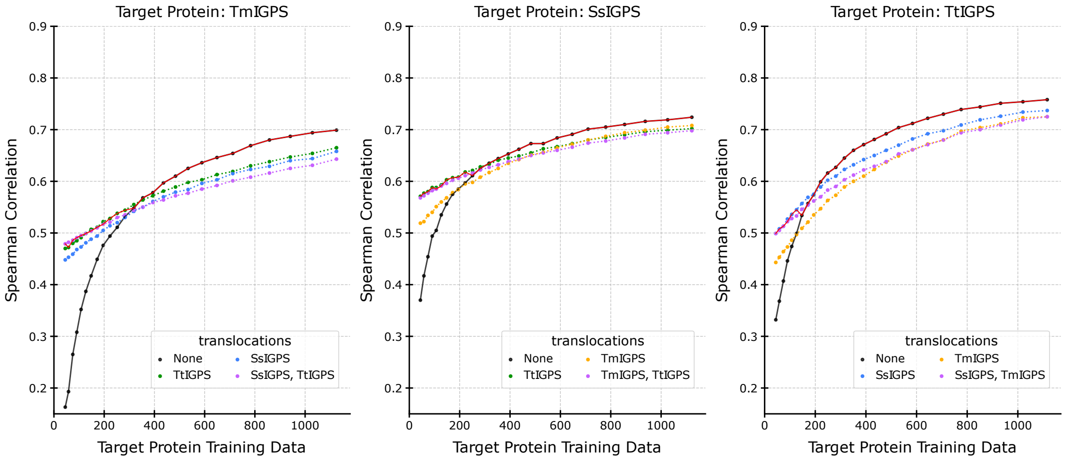


Figure S30.1: IGPS, ESM-1v pLM, and RF predictor.

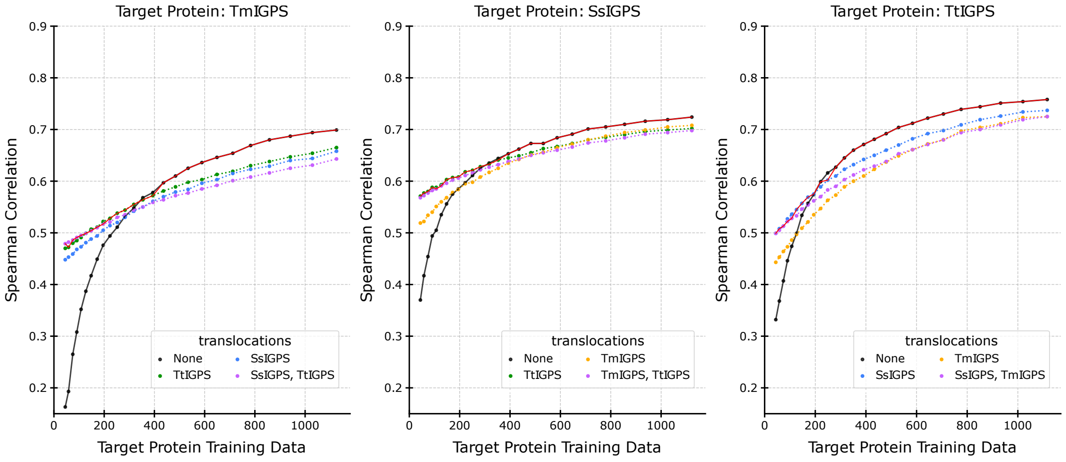


Figure S30.2: IGPS, ESM-1v pLM, and RF predictor.

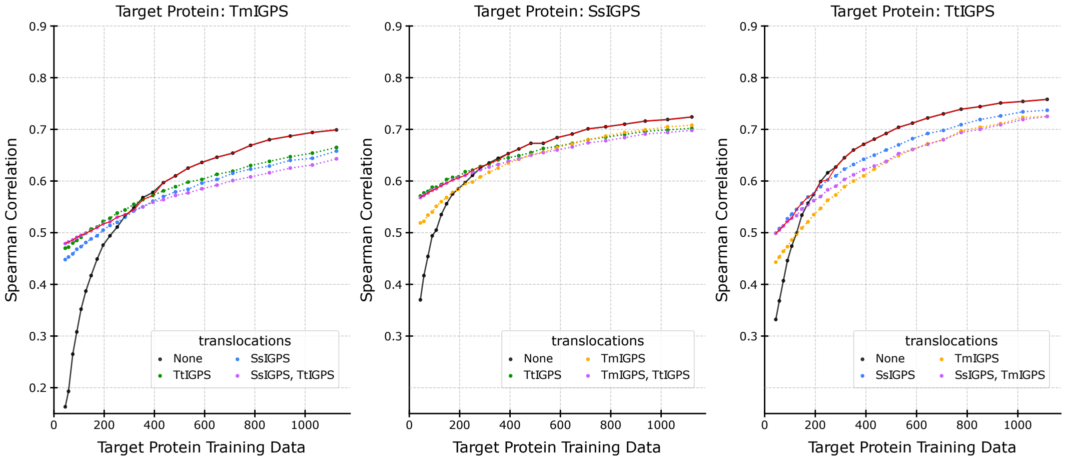


Figure S30.3: IGPS, ESM-1v pLM, and RF predictor.

#### Supplemental S31: GFP, ESM-1v, SVR, Statistical-Greedy - Individual-Greedy - Individual-Select

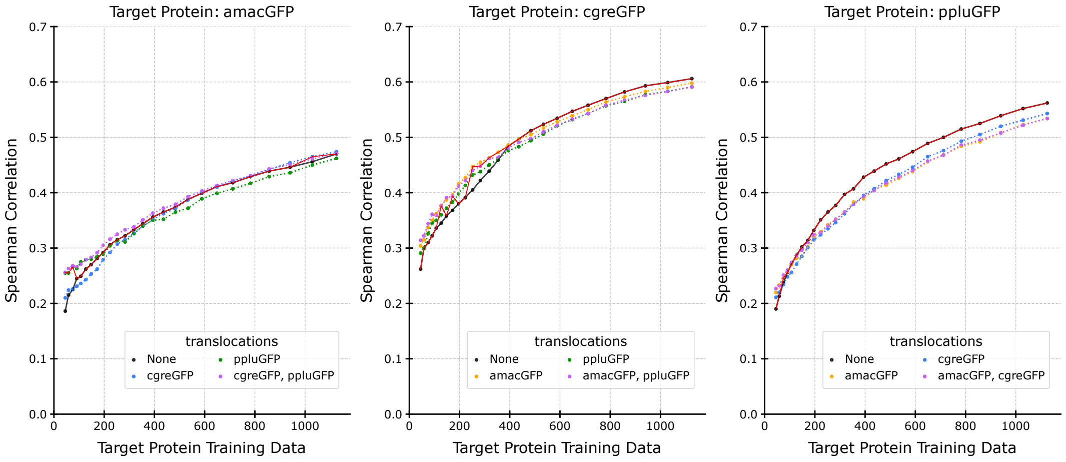


Figure S31.1: GFP, ESM-1v pLM, and SVR predictor.

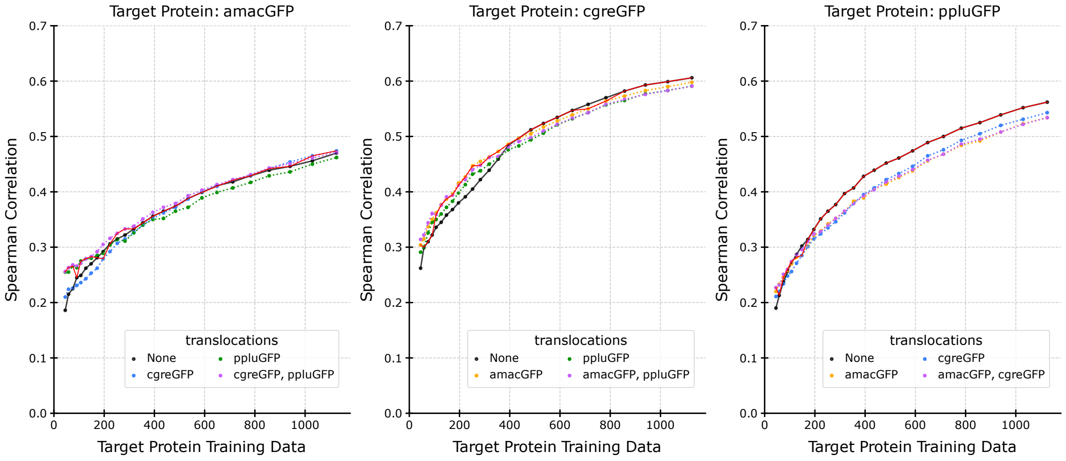


Figure S31.2: GFP, ESM-1v pLM, and SVR predictor.

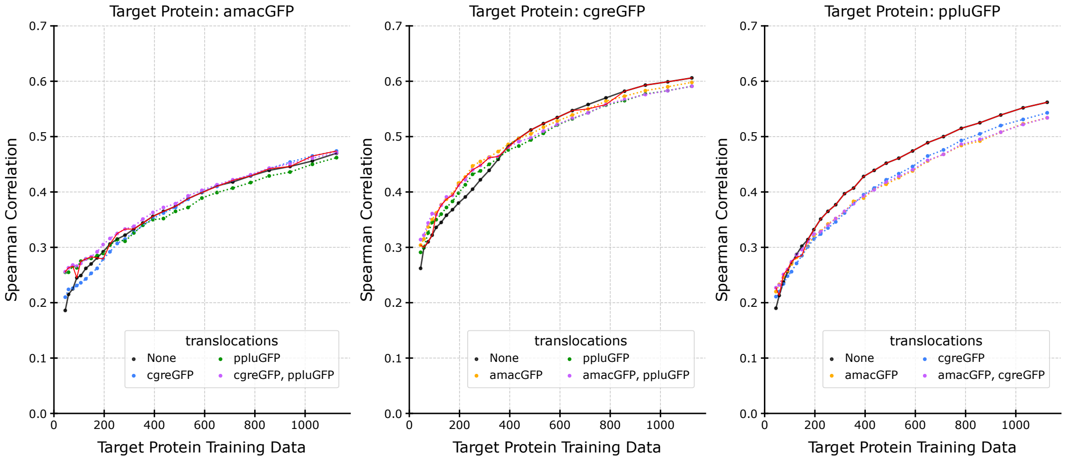


Figure S31.3: GFP, ESM-1v pLM, and SVR predictor.

#### Supplemental S32: GFP, ESM-1v, Lasso, Statistical-Greedy - Individual-Greedy - Individual-Select

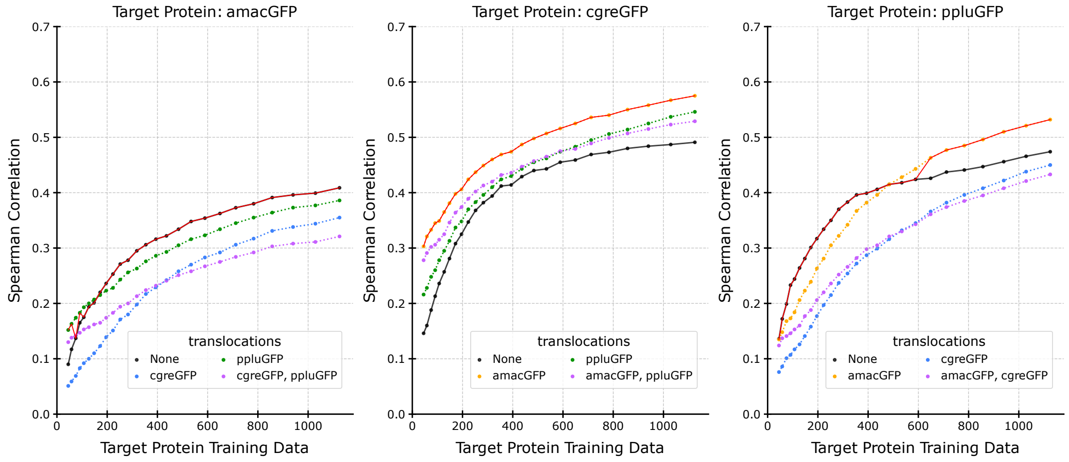


Figure S32.1: GFP, ESM-1v pLM, and Lasso predictor.

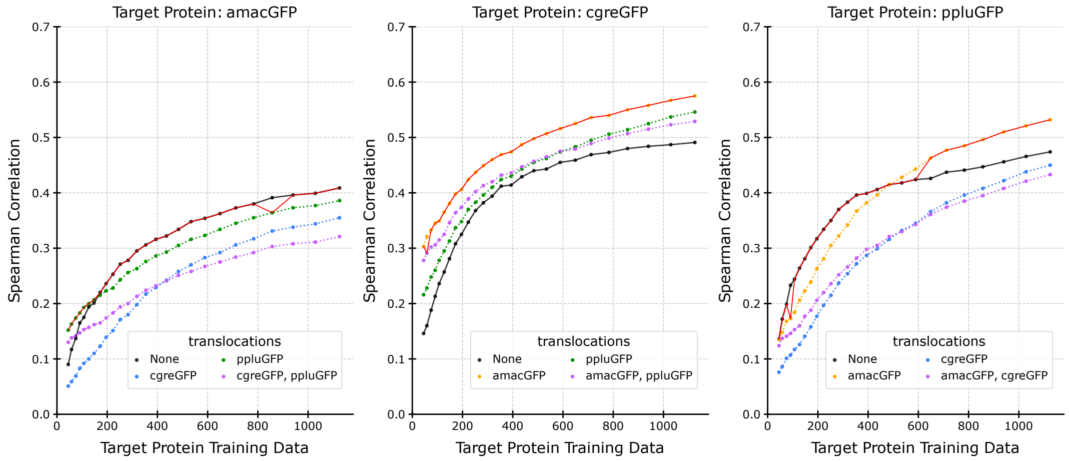


Figure S32.2: GFP, ESM-1v pLM, and Lasso predictor.

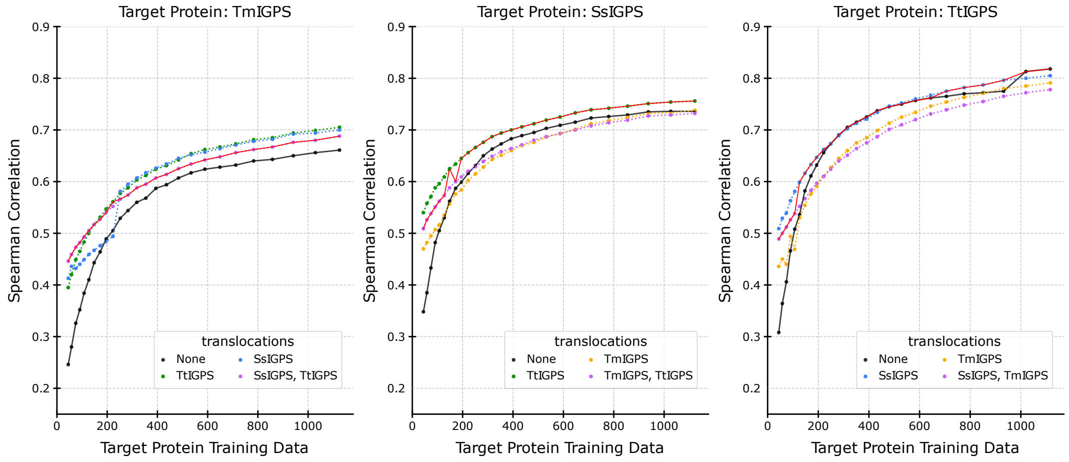


Figure S32.3: GFP, ESM-1v pLM, and Lasso predictor.

#### Supplemental S33: GFP, ESM-1v, RF, Statistical-Greedy - Individual-Greedy - Individual-Select

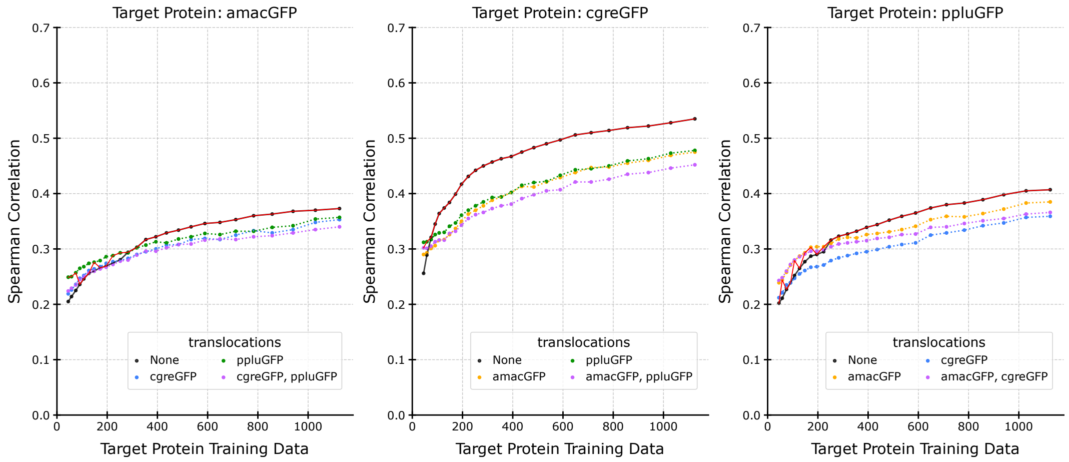


Figure S33.1: GFP, ESM-1v pLM, and RF predictor.

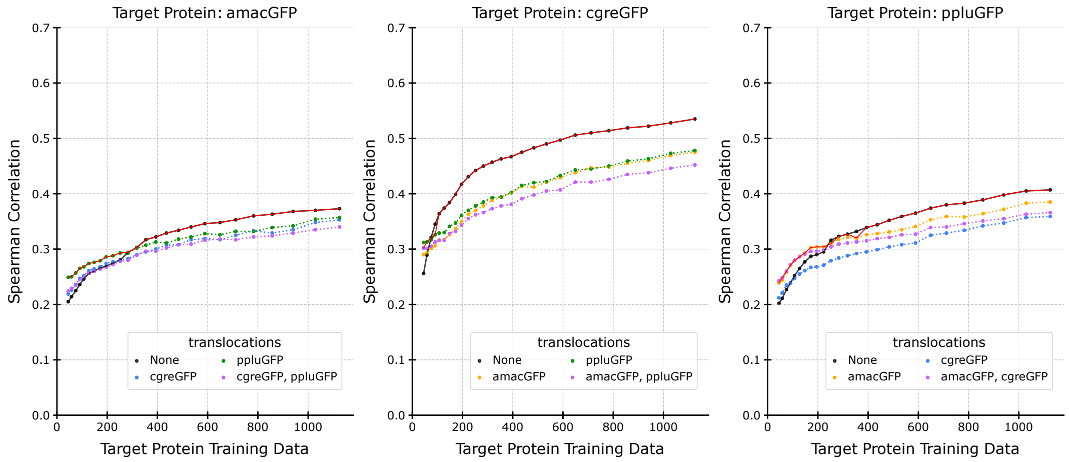


Figure S33.2: GFP, ESM-1v pLM, and RF predictor.

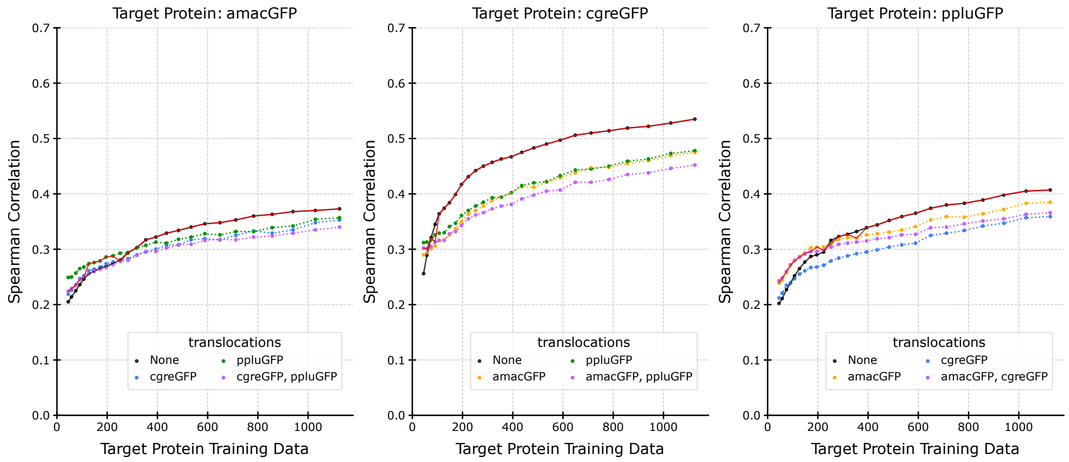


Figure S33.3: GFP, ESM-1v pLM, and RF predictor.

#### Supplemental S34: SARS-CoV-2 Spike protein Cell Entry, ESM2, SVR, Statistical-Greedy - Individual-Greedy - Individual-Select

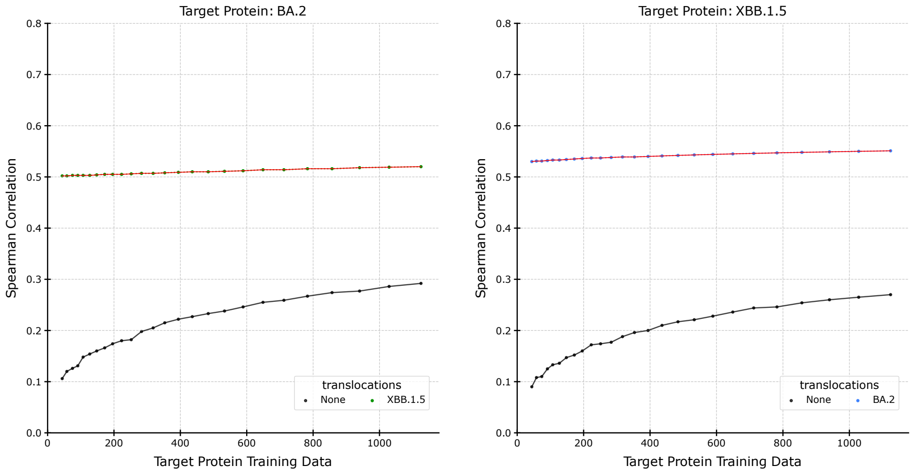


Figure S34.1: SARS-CoV-2 Spike protein Cell Entry, ESM2 pLM, and SVR predictor.

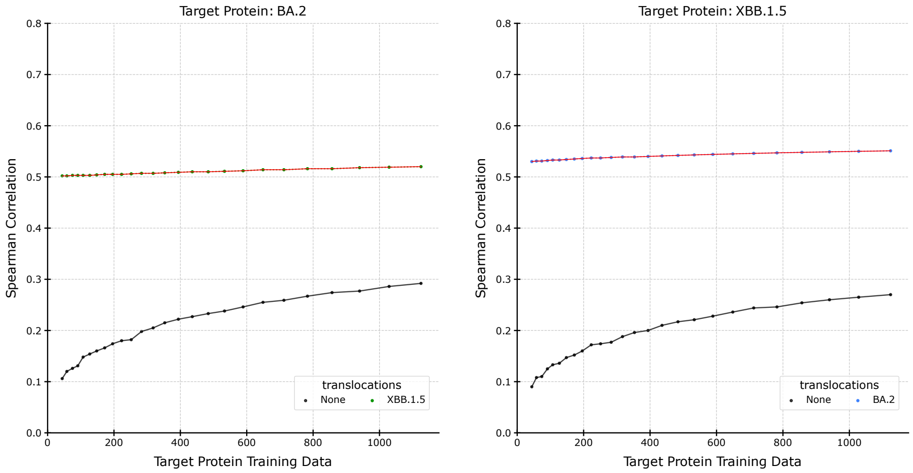


Figure S34.2: SARS-CoV-2 Spike protein Cell Entry, ESM2 pLM, and SVR predictor.

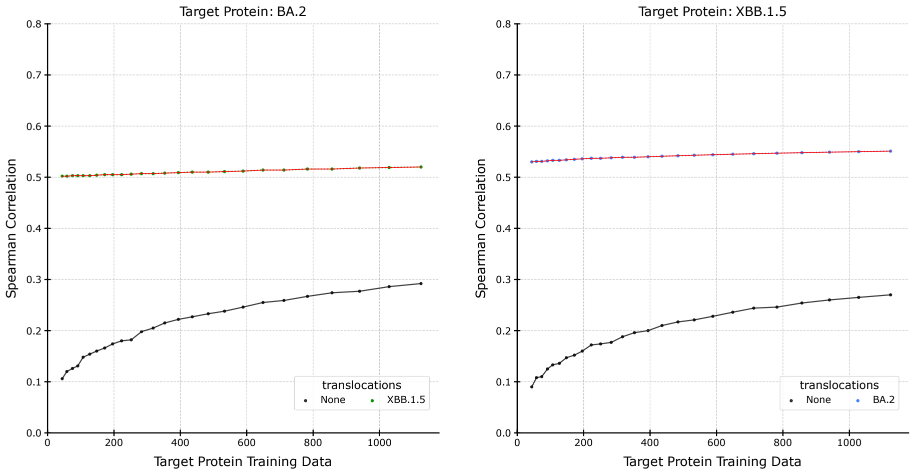


Figure S34.3: SARS-CoV-2 Spike protein Cell Entry, ESM2 pLM, and SVR predictor.

#### Supplemental S35: SARS-CoV-2 Spike protein Cell Entry, ESM2, Lasso, Statistical-Greedy - Individual-Greedy - Individual-Select

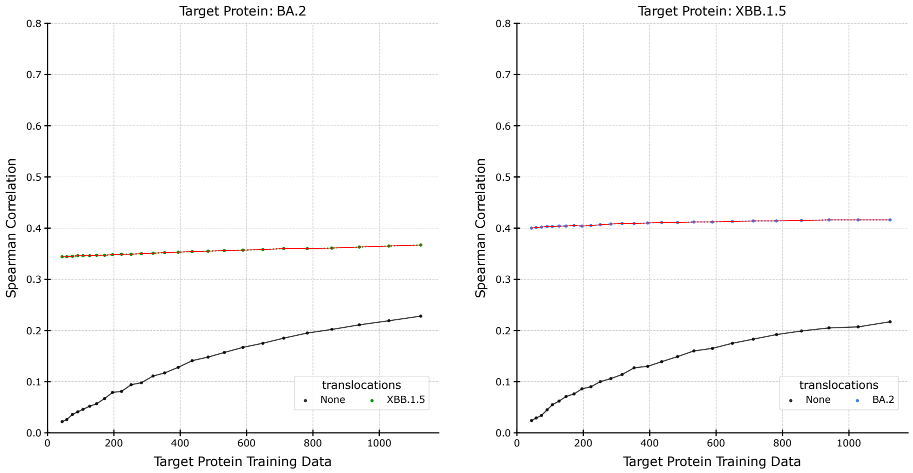


Figure S35.1: SARS-CoV-2 Spike protein Cell Entry, ESM2 pLM, and lasso predictor.

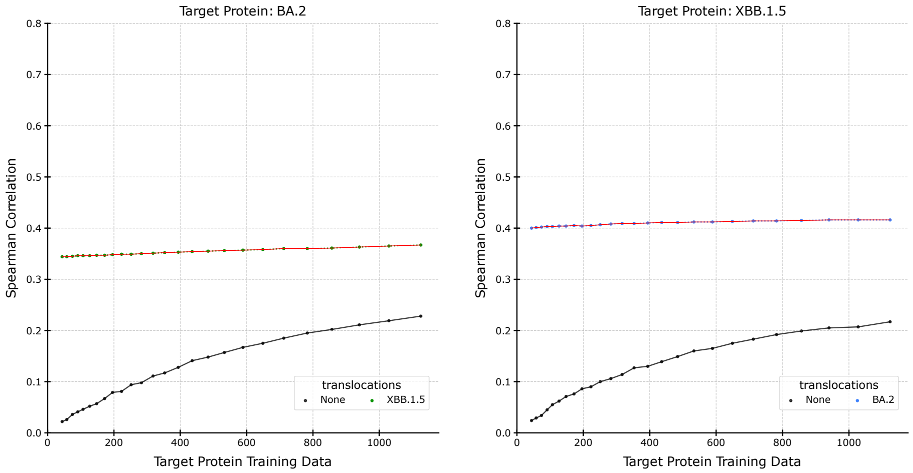


Figure S35.2: SARS-CoV-2 Spike protein Cell Entry, ESM2 pLM, and Lasso predictor.

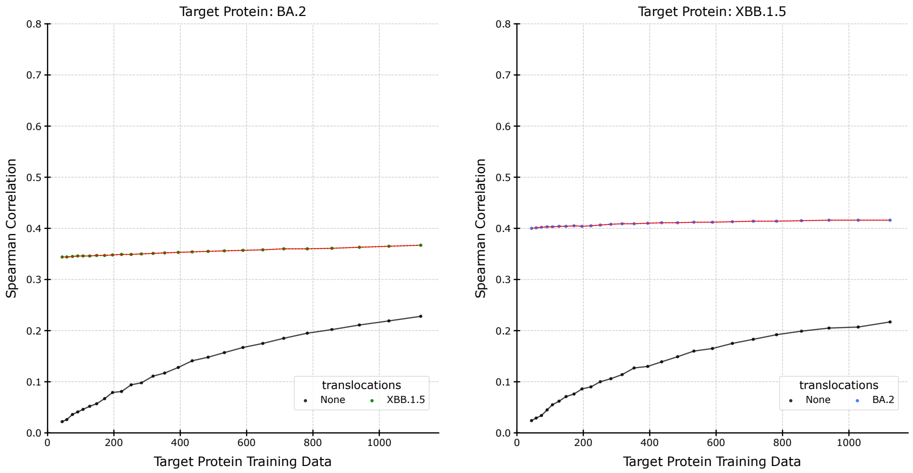


Figure S35.3: SARS-CoV-2 Spike protein Cell Entry, ESM2 pLM, and lasso predictor.

#### Supplemental S36: SARS-CoV-2 Spike protein Cell Entry, ESM2, RF, Statistical-Greedy - Individual-Greedy - Individual-Select

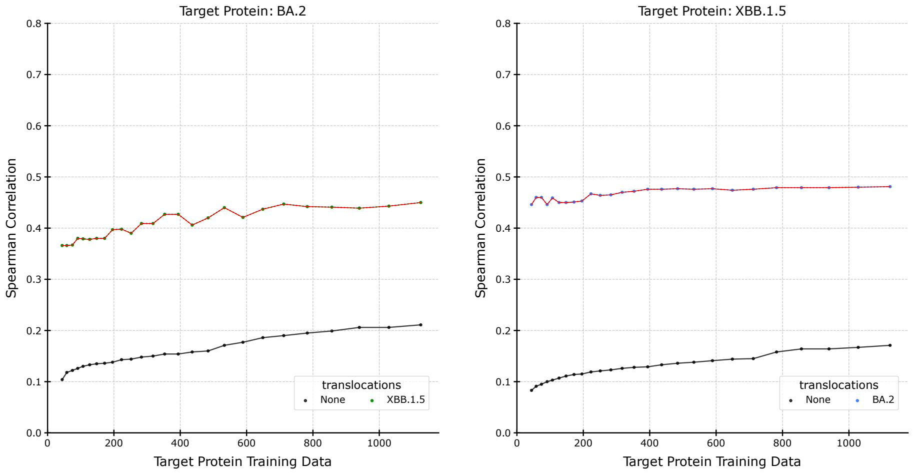


Figure S36.1: SARS-CoV-2 Spike protein Cell Entry, ESM2 pLM, and RF predictor.

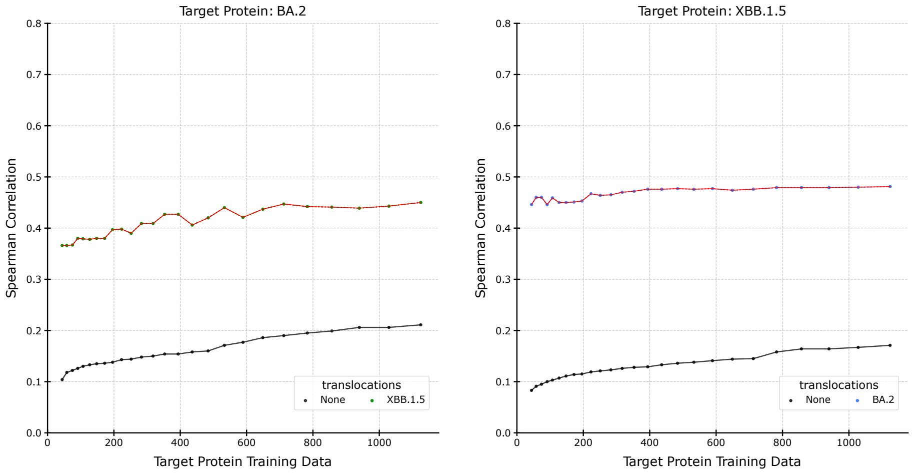


Figure S36.2: SARS-CoV-2 Spike protein Cell Entry, ESM2 pLM, and RF predictor.

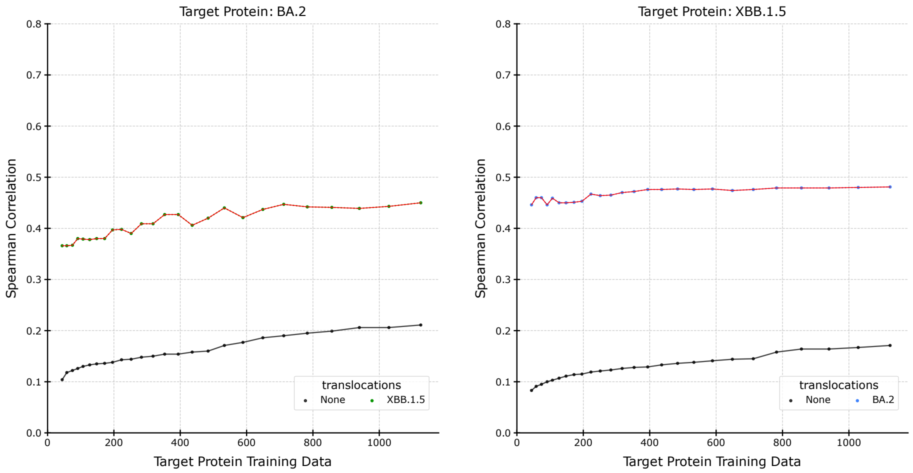


Figure S36.3: SARS-CoV-2 Spike protein Cell Entry, ESM2 pLM, and RF predictor.

#### Supplemental S37: SARS-CoV-2 Spike protein ACE2 Binding, ESM2, SVR, Statistical-Greedy - Individual-Greedy - Individual-Select

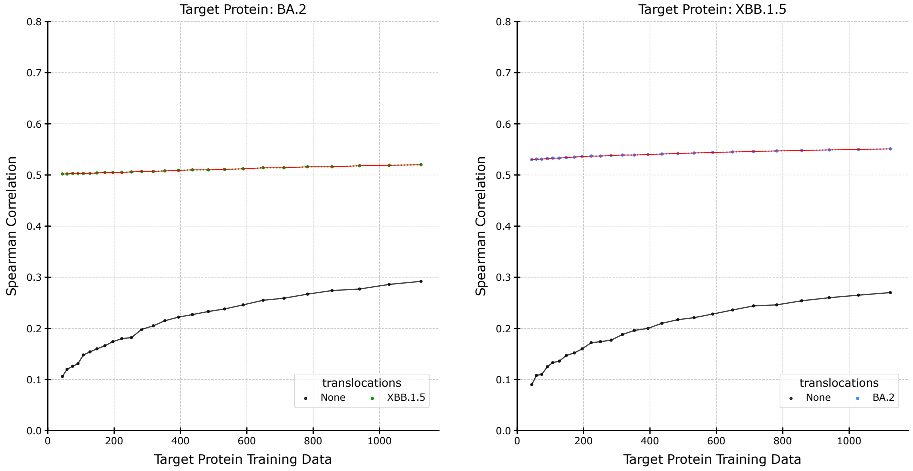


Figure S37.1: SARS-CoV-2 Spike protein ACE2 Binding, ESM2 pLM, and SVR predictor.

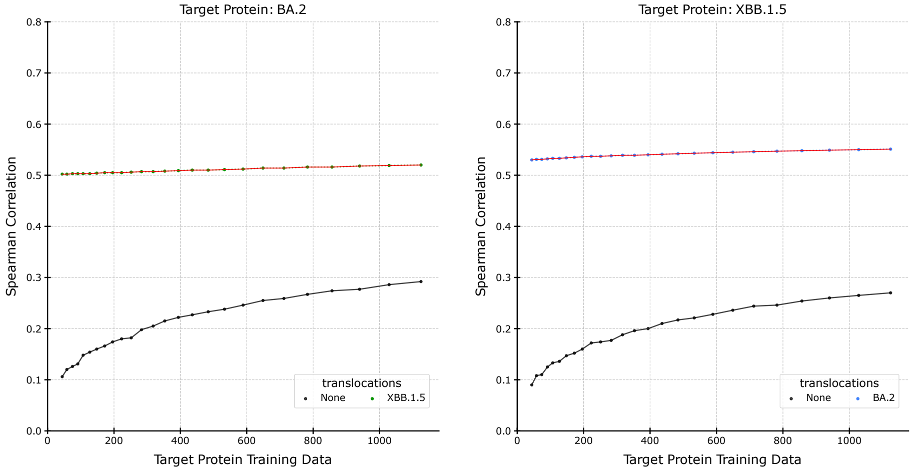


Figure S37.2: SARS-CoV-2 Spike protein ACE2 Binding, ESM2 pLM, and SVR predictor.

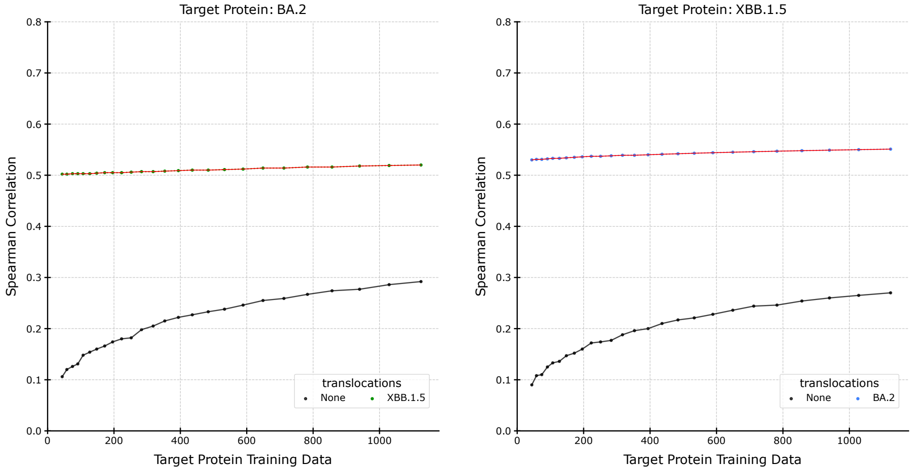


Figure S37.3: SARS-CoV-2 Spike protein ACE2 Binding, ESM2 pLM, and SVR predictor.

#### Supplemental S38: SARS-CoV-2 Spike protein ACE2 Binding, ESM2, Lasso, Statistical-Greedy - Individual-Greedy - Individual-Select

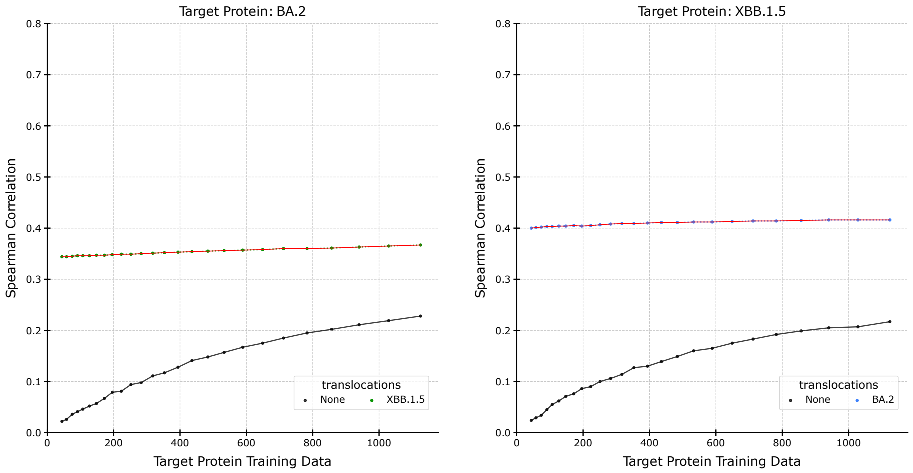


Figure S38.1: SARS-CoV-2 Spike protein ACE2 Binding, ESM2 pLM, and lasso predictor.

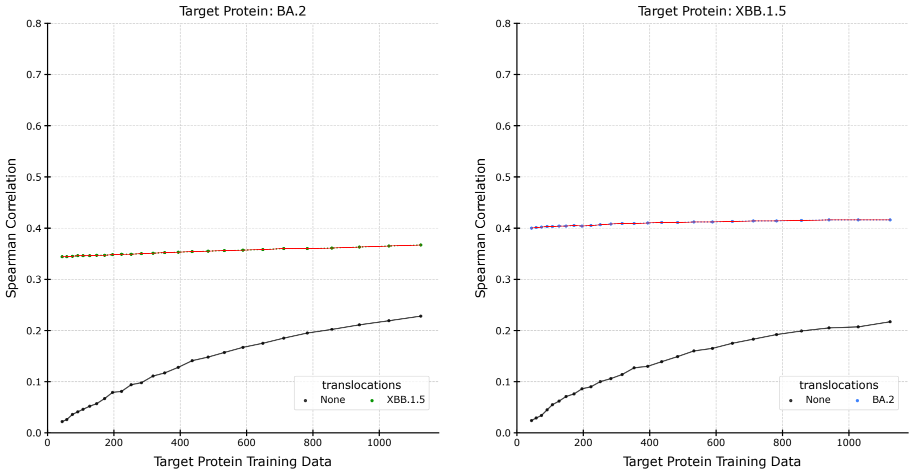


Figure S38.2: SARS-CoV-2 Spike protein ACE2 Binding, ESM2 pLM, and Lasso predictor.

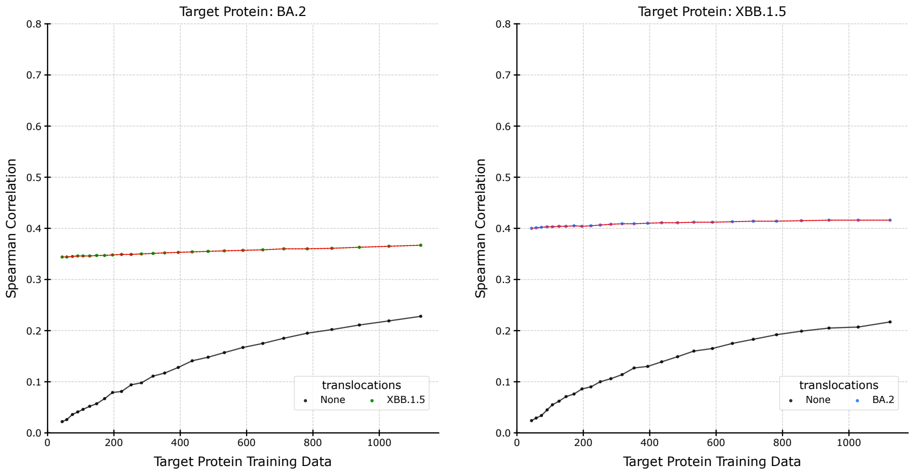


Figure S38.3: SARS-CoV-2 Spike protein ACE2 Binding, ESM2 pLM, and lasso predictor.

#### Supplemental S39: SARS-CoV-2 Spike protein ACE2 Binding, ESM2, RF, Statistical-Greedy - Individual-Greedy - Individual-Select

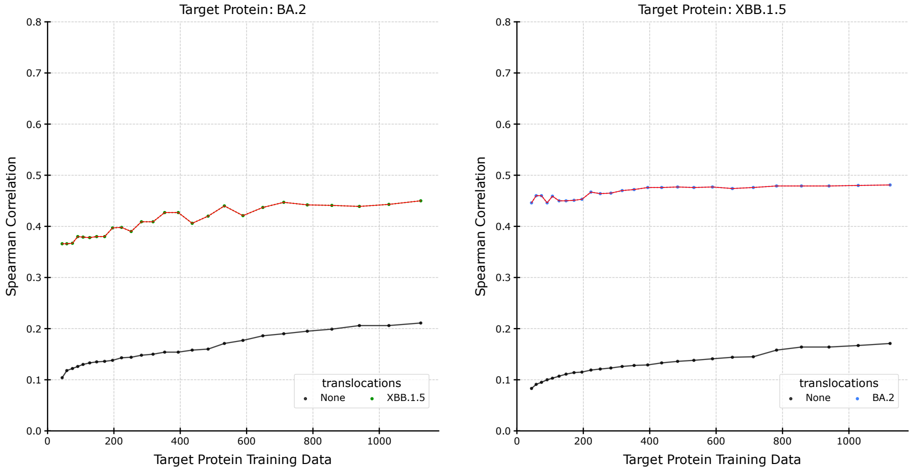


Figure S39.1: SARS-CoV-2 Spike protein ACE2 Binding, ESM2 pLM, and RF predictor.

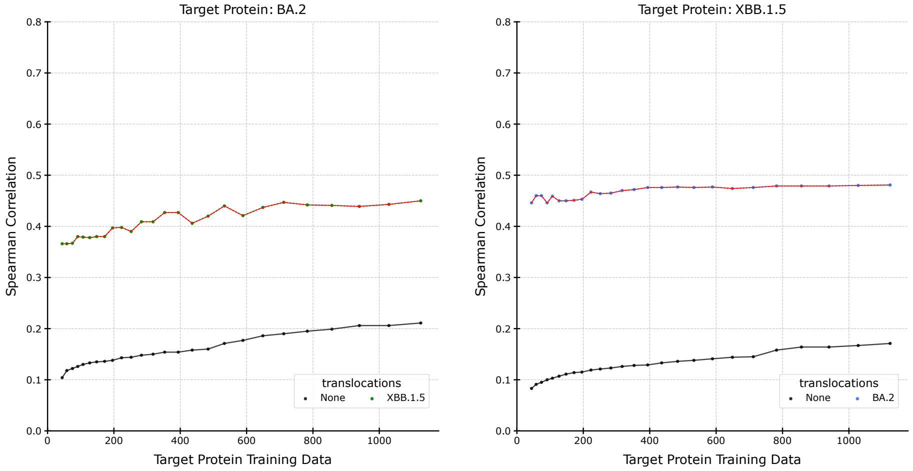


Figure S39.2: SARS-CoV-2 Spike protein ACE2 Binding, ESM2 pLM, and RF predictor.

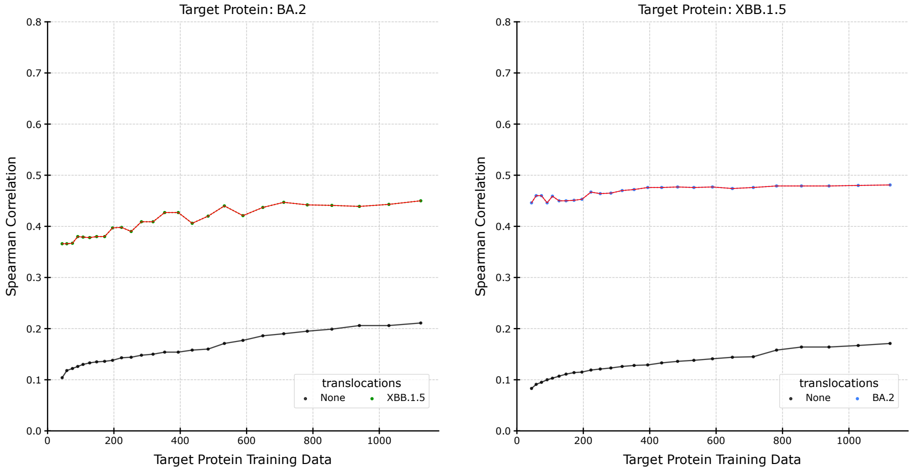


Figure S39.3: SARS-CoV-2 Spike protein ACE2 Binding, ESM2 pLM, and RF predictor.

### Supplemental S40: IGPS, ESM2, SVR, Statistical-Greedy - Individual-Greedy - Individual-Select

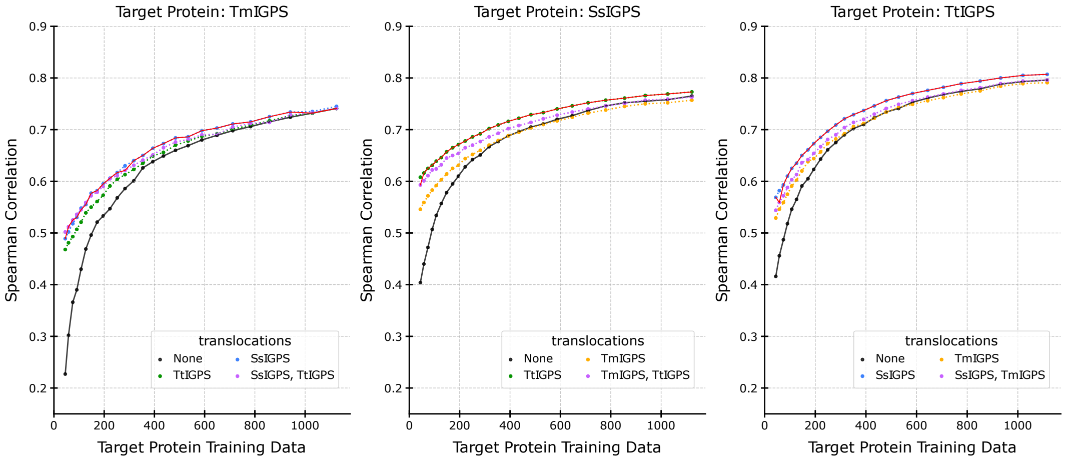


Figure S40.1: IGPS, ESM2 pLM, and SVR predictor.

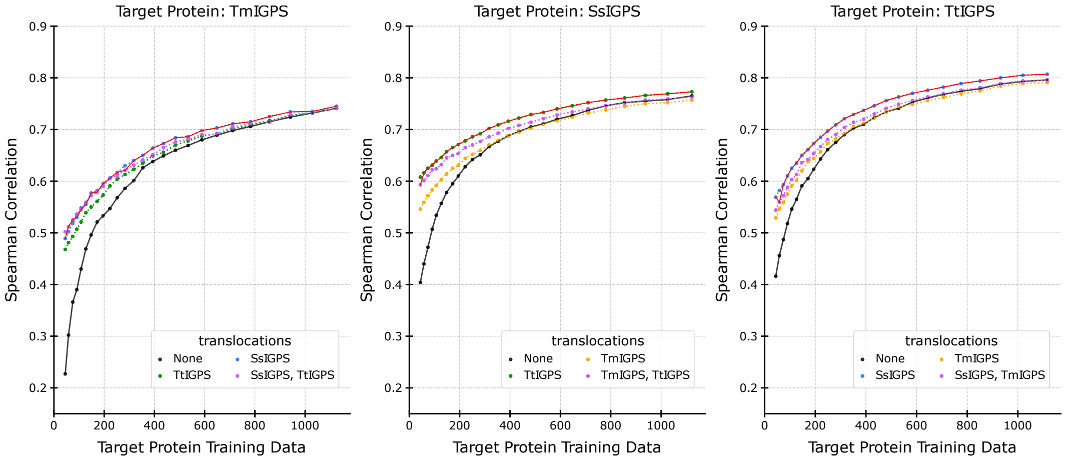


Figure S40.2: IGPS, ESM2 pLM, and SVR predictor.

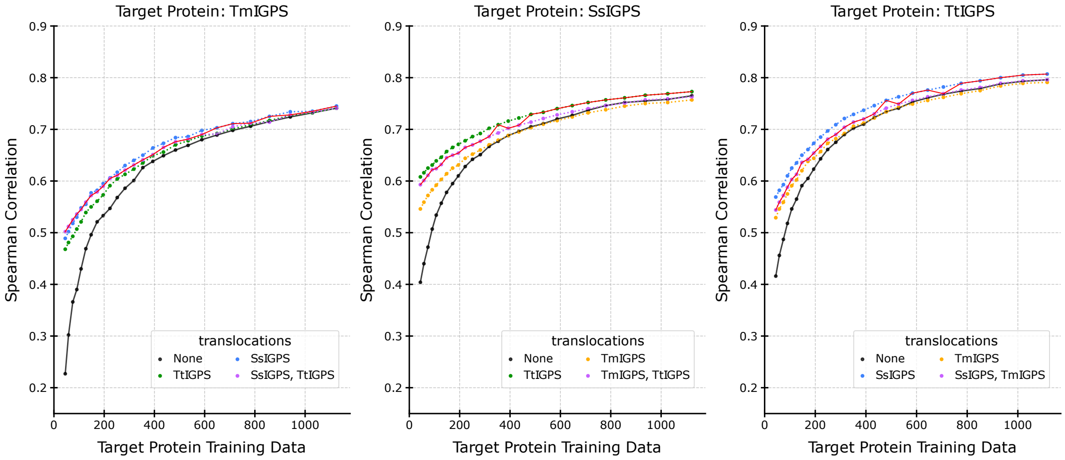


Figure S40.3: IGPS, ESM2 pLM, and SVR predictor.

#### Supplemental S41: IGPS, ESM2, Lasso, Statistical-Greedy - Individual-Greedy - Individual-Select

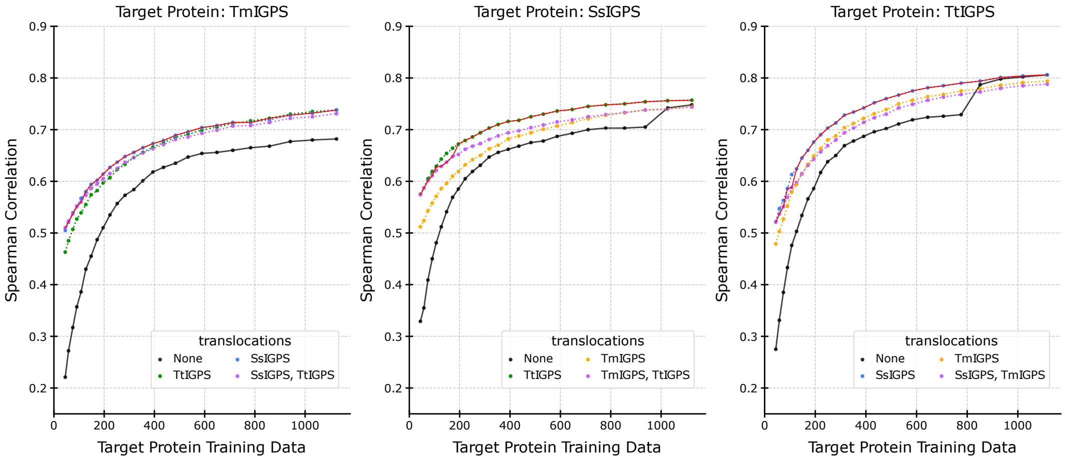


Figure S41.1: IGPS, ESM2 pLM, and Lasso predictor.

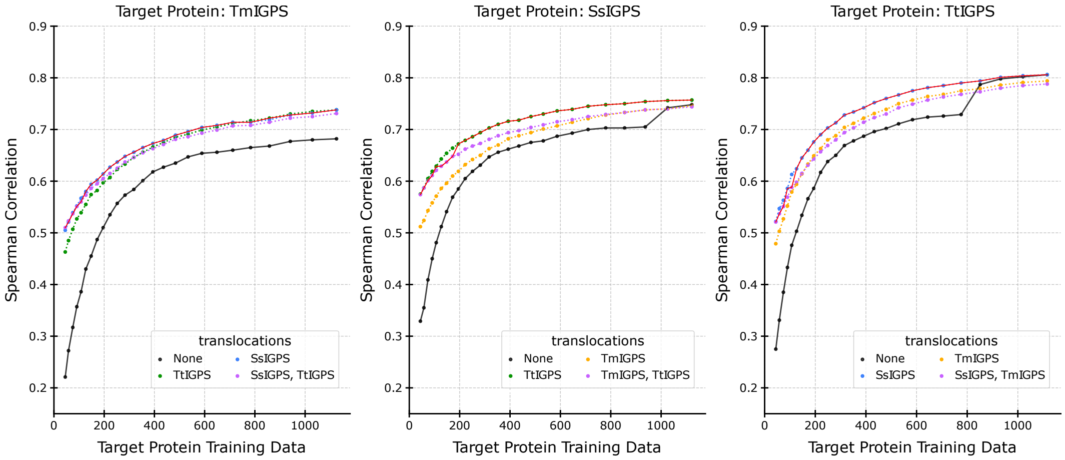


Figure S41.2: IGPS, ESM2 pLM, and Lasso predictor.

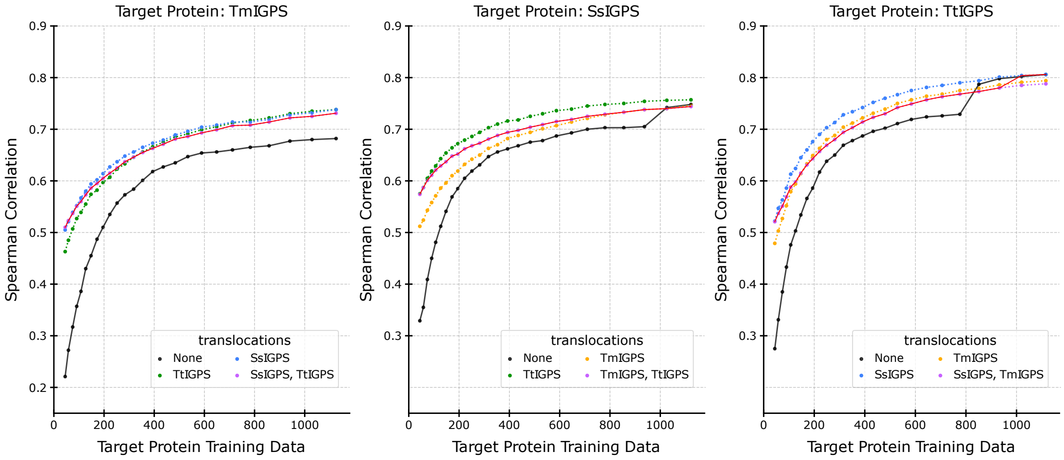


Figure S41.3: IGPS, ESM2 pLM, and Lasso predictor.

#### Supplemental S42: IGPS, ESM2, RF, Statistical-Greedy - Individual-Greedy - Individual-Select

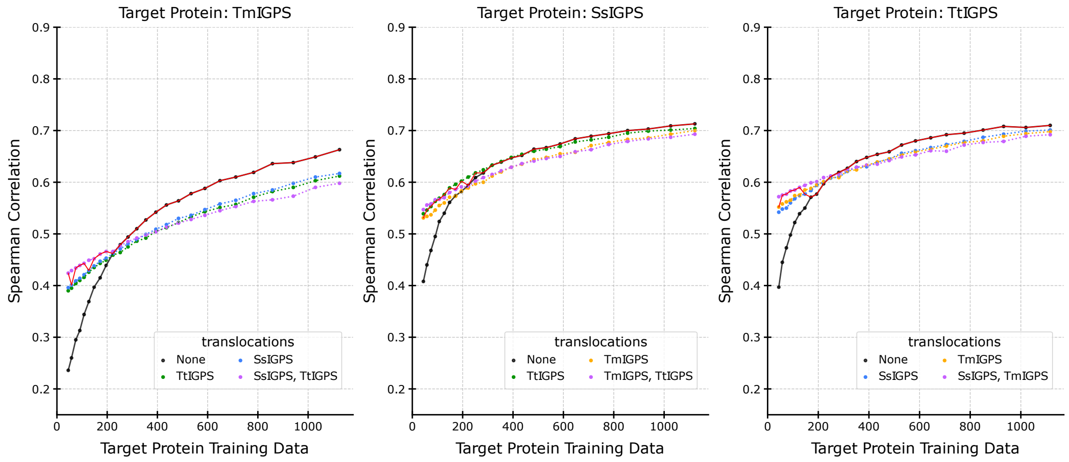


Figure S42.1: IGPS, ESM2 pLM, and RF predictor.

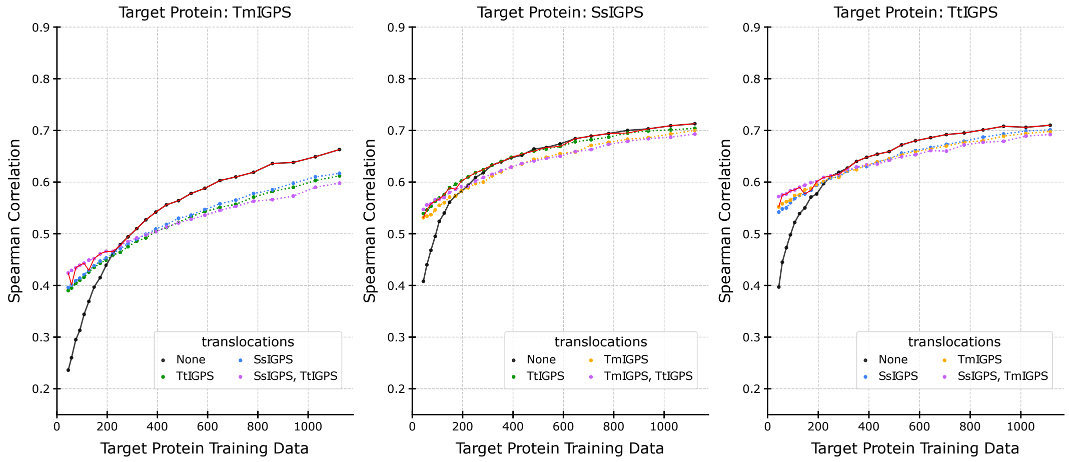


Figure S42.2: IGPS, ESM2 pLM, and RF predictor.

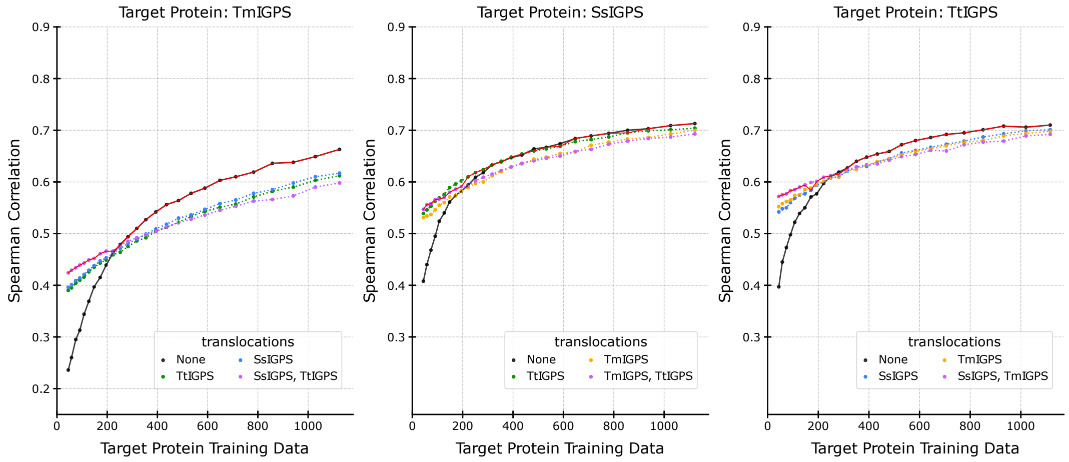


Figure S42.3: IGPS, ESM2 pLM, and RF predictor.

#### Supplemental S43: GFP, ESM2, SVR, Statistical-Greedy - Individual-Greedy - Individual-Select

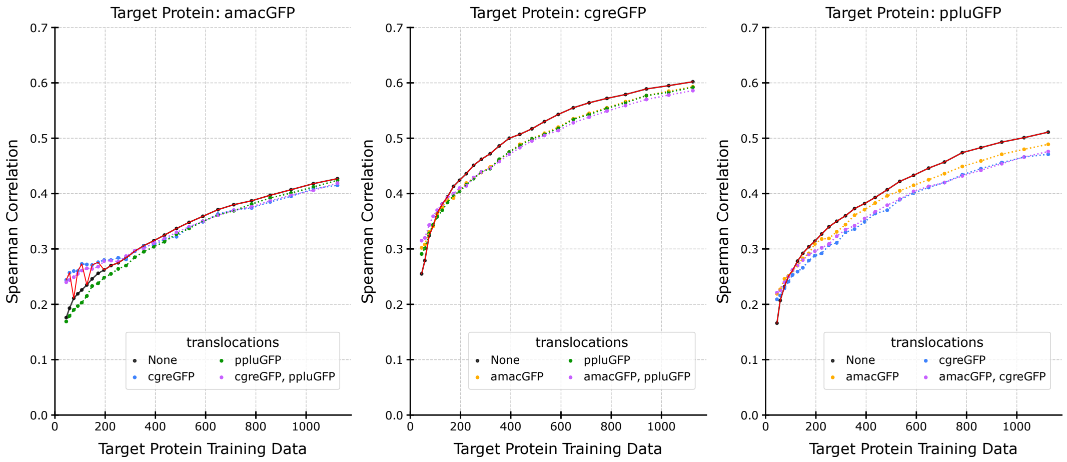


Figure S43.1: GFP, ESM2 pLM, and SVR predictor.

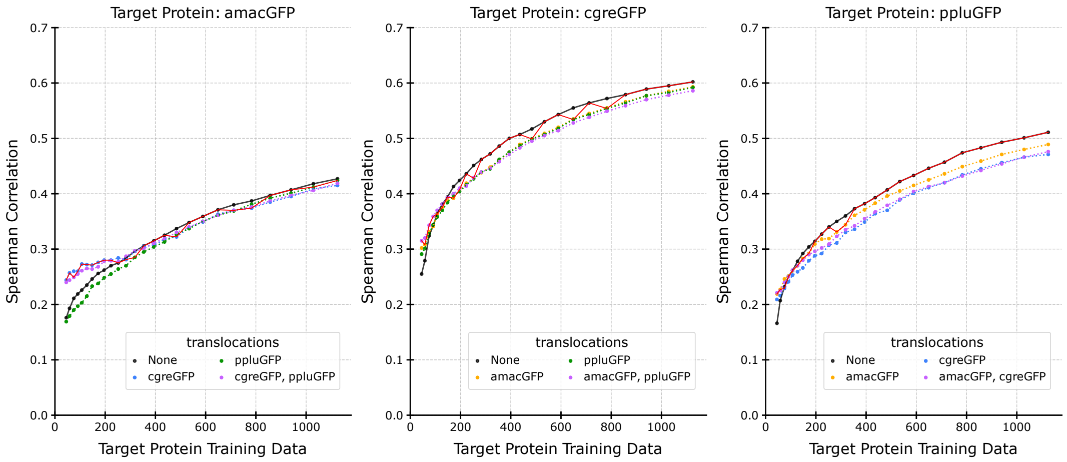


Figure S43.2: GFP, ESM2 pLM, and SVR predictor.

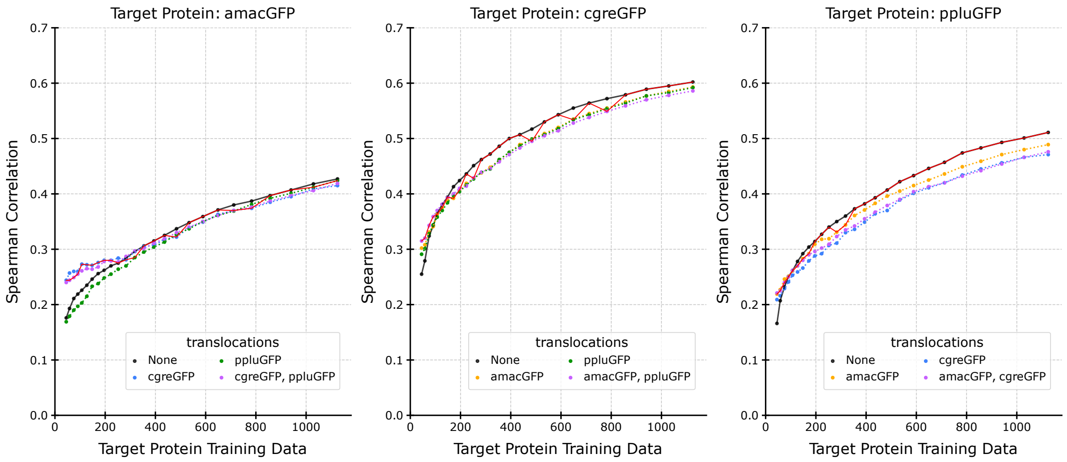


Figure S43.3: GFP, ESM2 pLM, and SVR predictor.

#### Supplemental S44: GFP, ESM2, Lasso, Statistical-Greedy - Individual-Greedy - Individual-Select

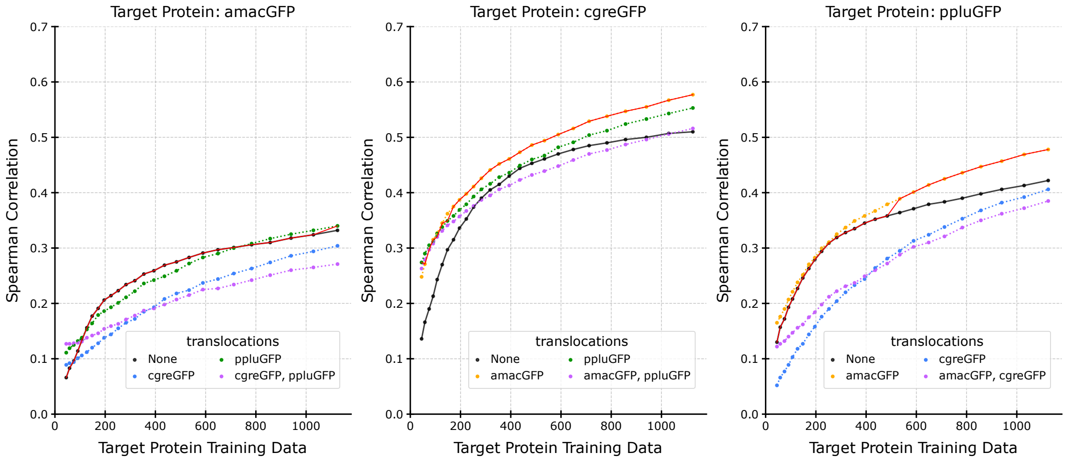


Figure S44.1: GFP, ESM2 pLM, and Lasso predictor.

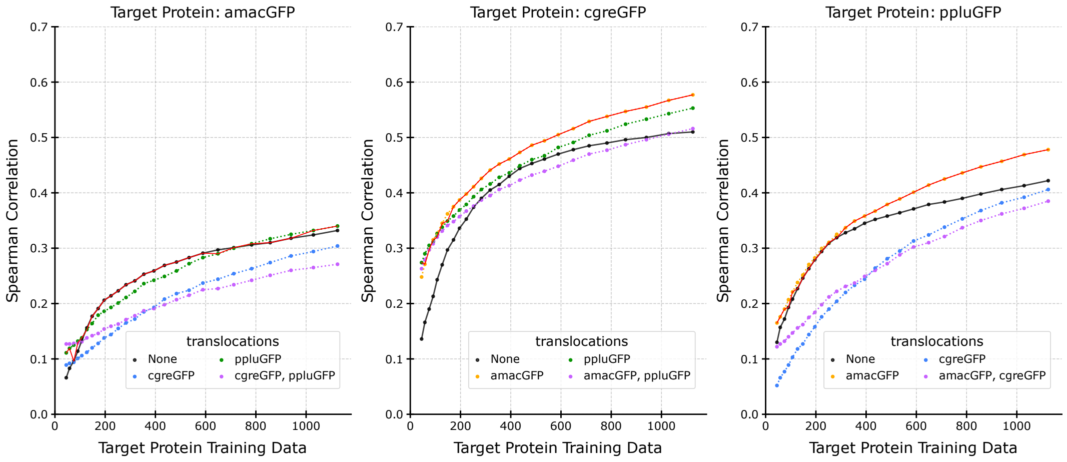


Figure S44.2: GFP, ESM2 pLM, and Lasso predictor.

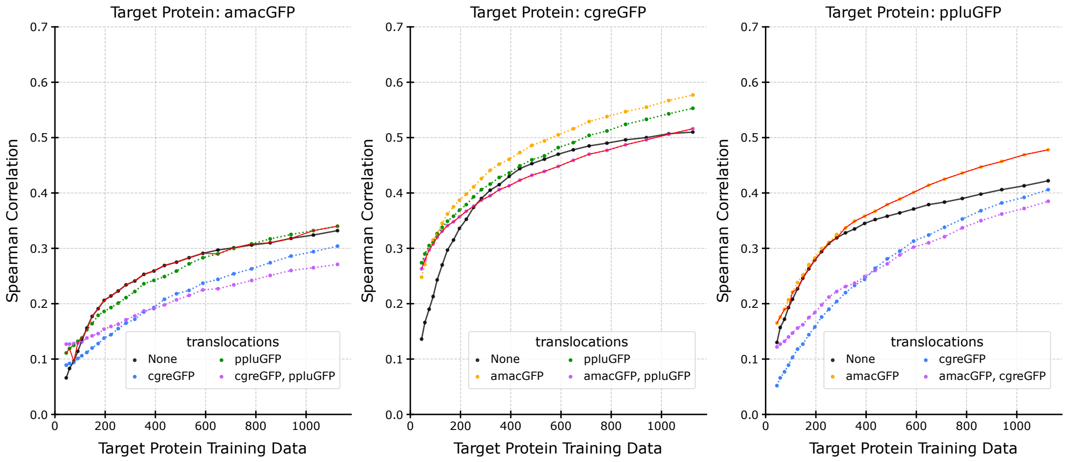


Figure S44.3: GFP, ESM2 pLM, and Lasso predictor.

#### Supplemental S45: GFP, ESM2, RF, Statistical-Greedy - Individual-Greedy - Individual-Select

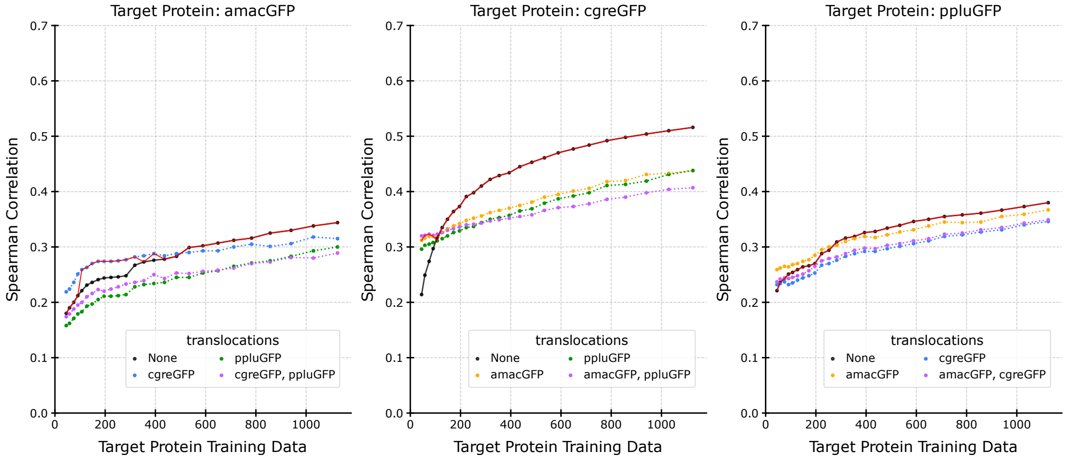


Figure S45.1: GFP, ESM2 pLM, and RF predictor.

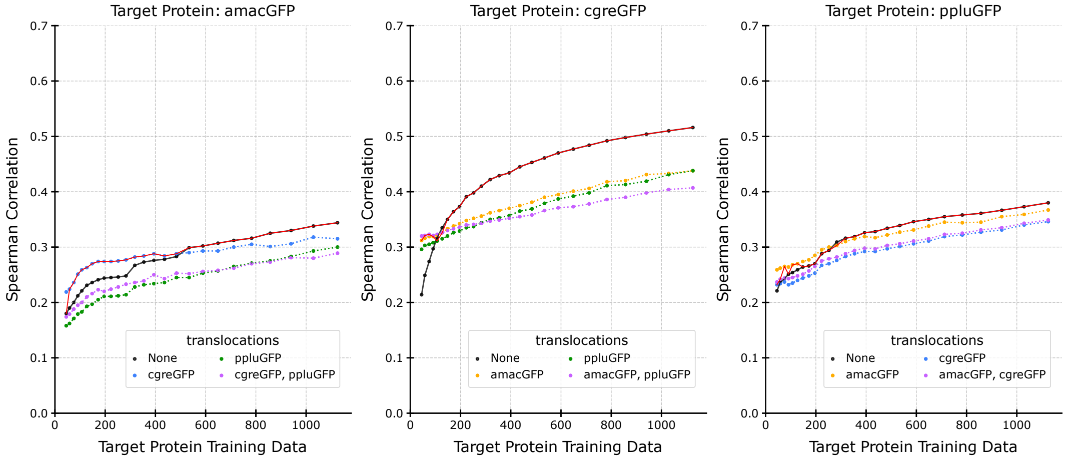


Figure S45.2: GFP, ESM2 pLM, and RF predictor.

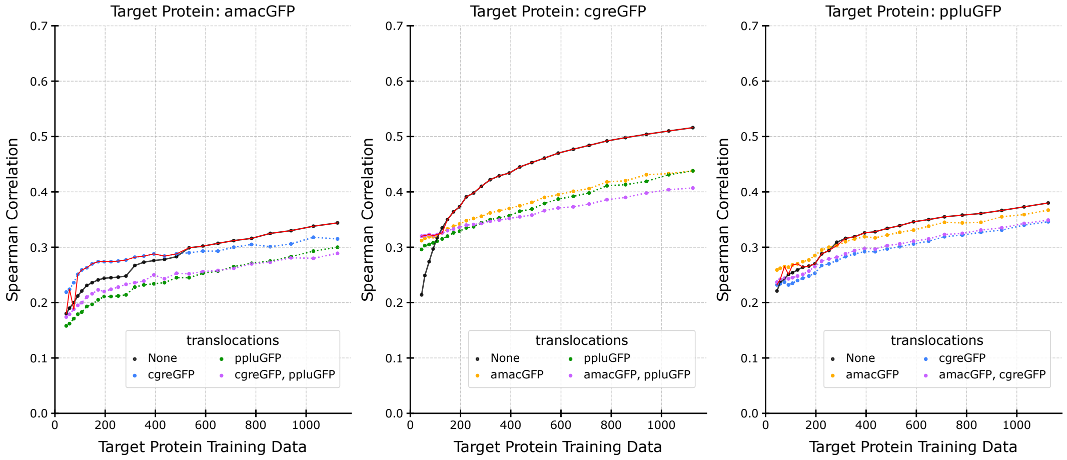


Figure S45.3: GFP, ESM2 pLM, and RF predictor.
